# Distinct goal location beta-frequency dynamics in hippocampus and prefrontal cortex across learning

**DOI:** 10.1101/2025.08.11.669642

**Authors:** Glingna Wang, Nan Zhou, Zachary M. Leveroni, Pinar Toptas, Audrey E. Kaye, Jaquelin Gutierrez, Amanda Rodriguez-Leon, Jai Y. Yu

## Abstract

Neural activity at goal locations contributes to learning by providing feedback on the success of preceding actions. This period engages neocortical and hippocampal networks, which serve distinct functions in reward processing and in forming associations between experience and reward. A neocortical network signature for reward feedback processing is beta oscillations (15-30 Hz). Beta oscillations are thought to coordinate distributed neural processes across brain regions. However, it is unknown whether beta oscillations coordinate hippocampal-neocortical networks during the goal period, or how their dynamics relate to learning. Here, we show that beta oscillations occur in both hippocampal CA1 and the prefrontal cortex (PFC) when rats reach goal locations in spatial navigation tasks. Despite the presence of beta oscillations in both regions after goal entry, beta activity in each region differed in spectral and temporal properties. These differences suggest that the hippocampus and PFC are weakly coupled at the beta frequency. We found that across learning, the strengths of PFC and CA1 beta oscillations were inversely related: PFC beta power increased and CA1 beta power decreased. Beta burst properties in PFC also had an inverse relationship to those of hippocampal sharp-wave ripples (SWRs), a prominent hippocampal process required for learning. We found a subset of PFC neurons modulated by both beta and hippocampal SWRs, which had distinct task-related firing patterns. Our results suggest that during outcome processing at goal locations, the neocortex and hippocampus are locally modulated at the beta frequency and then become coordinated for memory-related processes during SWRs.

## Introduction

Learning leads to changes in distributed neural networks across the brain. During learning, diverse neural processes across brain networks need to be coordinated, which is hypothesized to rely on neural oscillations that rhythmically couple activity in connected brain regions. This coupling between groups of brain regions is thought to be dynamic, creating functionally interacting subnetworks to support specific neural processes when required. A neural oscillation that is hypothesized to support these functions is beta (15-30 Hz) (Bibbig et al., 2002; Gourévitch et al., 2010; Igarashi et al., 2014; Jayachandran et al., 2023; Kopell et al., 2011; Lundqvist et al., 2024; Miles et al., 2023; Spitzer & Haegens, 2017; Symanski et al., 2022). Beta oscillations occur in many brain regions, including sensory (Gervais et al., 2007; Kay et al., 1996; Martin et al., 2004), neocortical (Baker et al., 1997; Buschman et al., 2012; Cohen et al., 2007; Cohen et al., 2015; Dubey et al., 2023; HajiHosseini & Holroyd, 2015; HajiHosseini et al., 2020; Iturra-Mena et al., 2023; Kawasaki & Yamaguchi, 2013; Koloski, Hulyalkar, et al., 2024; MacKay & Mendonça, 1995; Pesaran et al., 2008; Voloh et al., 2020; Walsh et al., 2025; Xiao et al., 2024), subcortical (DeCoteau et al., 2007; Feingold et al., 2015; Howe et al., 2011; Lansink et al., 2009; Leventhal et al., 2012; Quinn et al., 2010; Sturman & Moghaddam, 2012) and limbic regions (Amaya et al., 2024; Berke et al., 2008; Fourcaud-Trocmé et al., 2019; França et al., 2014; França et al., 2021; Igarashi et al., 2014; Jackson et al., 2024; Jayachandran et al., 2023; Kay & Freeman, 1998; Rangel et al., 2015; Rangel et al., 2016; Samson et al., 2017; Symanski et al., 2022). Beta oscillations are associated with a wide range of cognitive functions, including sensory perception (Kay, 2014), reward processing (Cohen et al., 2007), top-down control (Engel & Fries, 2010) and novelty detection (Marco-Pallarés et al., 2015). Despite their widespread occurrence and association with diverse cognitive functions, a less-understood aspect of beta oscillations is their contribution to modulating or coordinating brain networks during learning, when rules are not fully understood and performance is improving. Most studies investigating beta dynamics across the brain have primarily focused on performance after initial learning. However, three lines of work in sensory-limbic, striatal, and amygdala networks point to beta playing a role in supporting learning. First, sensory-association learning studies suggest that beta activity coordinates sensory-limbic networks to enable the formation of associations between the sensory cue and its behavioral relevance (Fourcaud-Trocmé et al., 2019; Martin et al., 2007; Martin et al., 2004). Second, beta oscillation dynamics in the ventral striatum change as habits are formed (Howe et al., 2011). Lastly, beta oscillations in the basolateral amygdala can drive reward preference (Amaya et al., 2024). However, the patterns of learning-related changes in beta oscillations across brain networks remain unclear, since the dynamics of beta during learning outside of olfactory and striatal networks are not known. Here, we asked how beta dynamics contribute to spatial learning in hippocampal-neocortical networks, whose component regions are known to play central roles in supporting memory processing and guiding behavior.

To understand how beta oscillations in hippocampal-neocortical networks change with learning, we focused on an important period for driving learning: when the reward outcome is revealed. We refer to this period as the “goal” phase of a task. Beta oscillations have been observed during equivalent task periods in both human and non-human animal studies. Neocortical beta oscillations during this period have been associated with processing outcomes, where feedback from the presence or absence of the reward can inform future actions and drive learning (Cohen et al., 2011). Goal-related beta oscillations have been observed in frontal cortical regions (Cohen et al., 2007; Kawasaki & Yamaguchi, 2013), including the anterior cingulate cortex (ACC) (Iturra-Mena et al., 2023; Xiao et al., 2024), the prefrontal cortex (PFC) (HajiHosseini & Holroyd, 2015; HajiHosseini et al., 2020; Voloh et al., 2020) and the orbitofrontal cortex (OFC) (Koloski, Hulyalkar, et al., 2024; Koloski, O’Hearn, et al., 2024), in primate and rodent species. Beta oscillations in these frontal cortical regions have been hypothesized to provide top-down cortical control of connected networks (Babapoor-Farrokhran et al., 2017; Benchenane et al., 2011; Bressler & Richter, 2015; Brovelli et al., 2004; Buschman et al., 2012; Dubey et al., 2023; Engel & Fries, 2010; Pesaran et al., 2008).

Despite frontal cortical beta activity being implicated in outcome feedback processing, whether frontal cortical and hippocampal networks are coordinated in the beta-frequency range during the goal period remains unknown. Beta-frequency coordination of hippocampal-neocortical networks has been reported in olfactory sensory decision-making tasks during the cue presentation period (Igarashi et al., 2014; Jayachandran et al., 2023; Martin et al., 2007; Symanski et al., 2022), and is hypothesized to support the use of memory-based sensory associations for decision making (Spitzer & Haegens, 2017). However, goal-related beta-frequency oscillations have not been reported in the hippocampus. Beta oscillations in the hippocampus have so far been observed during exploration (Ahmed & Mehta, 2012), especially in novel environments (Berke et al., 2004), interaction with novel objects (França et al., 2014; França et al., 2021; Rangel et al., 2015), approaches to goals (Iwasaki et al., 2021; Lansink et al., 2009) as well as sensory stimulus sampling (Fourcaud-Trocmé et al., 2019; Gourévitch et al., 2010; Jayachandran et al., 2023; Kay & Freeman, 1998; Kay et al., 1996; Martin et al., 2007; Rangel et al., 2016; Symanski et al., 2022). Thus, it is unknown if and how hippocampal-neocortical networks are coordinated at the beta frequency during the goal phase of a task, and how this coordination relates to learning.

To understand the relationship between goal-related beta-frequency activity in hippocampal-neocortical networks and learning, we recorded local field potentials (LFPs) and single units from the hippocampus (dorsal CA1) and the PFC in rats learning spatial navigation tasks over five days. We observed beta oscillations in both regions at the goal locations of the tasks. However, we were surprised to find that beta oscillations in the PFC and CA1 had distinct spectral and temporal properties. The mismatch in frequency range and timing is consistent with each region engaging in distinct patterns of beta activity, likely with other brain regions, rather than being coupled with each other by beta. Over days of training, beta oscillations in the PFC and CA1 had distinct correlations with performance, where PFC beta increased in strength and CA1 beta decreased in strength. We also found an inverse relationship between the learning-related properties of PFC beta and a prominent hippocampal marker of learning, the sharp-wave ripple (SWR). Lastly, we found distinct task correlates in a subpopulation of PFC cells that showed modulation by both beta oscillations and hippocampal SWRs.

In contrast to previous results demonstrating coordinated hippocampal-neocortical beta modulation during sensory decision-making (Gourévitch et al., 2010; Jayachandran et al., 2023; Kay & Freeman, 1998; Kay et al., 1996; Martin et al., 2007; Rangel et al., 2016; Symanski et al., 2022), we show that beta oscillations in these networks are likely modulating distinct network processes during the goal phase of a task. Further, beta dynamics in the PFC and CA1 undergo opposite changes across learning. After goal entry, the neocortex and hippocampus are weakly coupled at the beta frequency but later coordinate during hippocampal SWRs. Our results support the hypothesis that at the goal, the neocortex is initially engaged in processing outcome-related information before coordinating with hippocampal networks for memory-related processes.

## Results

### Region-specific beta oscillations at goal locations

Rats (n=6) were trained on two spatial alternation tasks, interleaved across sessions, for five days (Table 1-1). In each task, reward was delivered when the three goal locations were visited in an alternating sequence (Fig. 1 A). We defined the goal location as the reward delivery device at the end of each maze arm (Fig. 1 B). Goal entry was defined as the time when the animal arrived at the goal and triggered the infrared sensor on the device. We first asked whether beta oscillations could be observed in both CA1 and PFC (Fig. 1-1 and Table 1-2). We observed power in the beta-frequency range (15-30 Hz) in both PFC and CA1 after the animal reached the goal locations (Fig. 1 B, Fig. 1-2). This was confirmed by comparing the power spectral density (PSD) between periods when the animal was moving on the maze arms (On Maze) or immobile at the goal location (At Goal) (Fig. 1 C-H and Fig. 1-3). The PSD for PFC showed a peak at approximately 20 Hz (Fig. 1 C), which was significantly stronger at the goal compared with on the maze (Fig. 1 E and G). In CA1, beta-frequency power was elevated during periods at the goal compared with on the maze (Fig. 1 D, F and H). We repeated these analyses using continuous wavelet transform and found the same pattern: beta power was elevated after but not before goal entry (Fig. 1-4).

**Figure 1.**
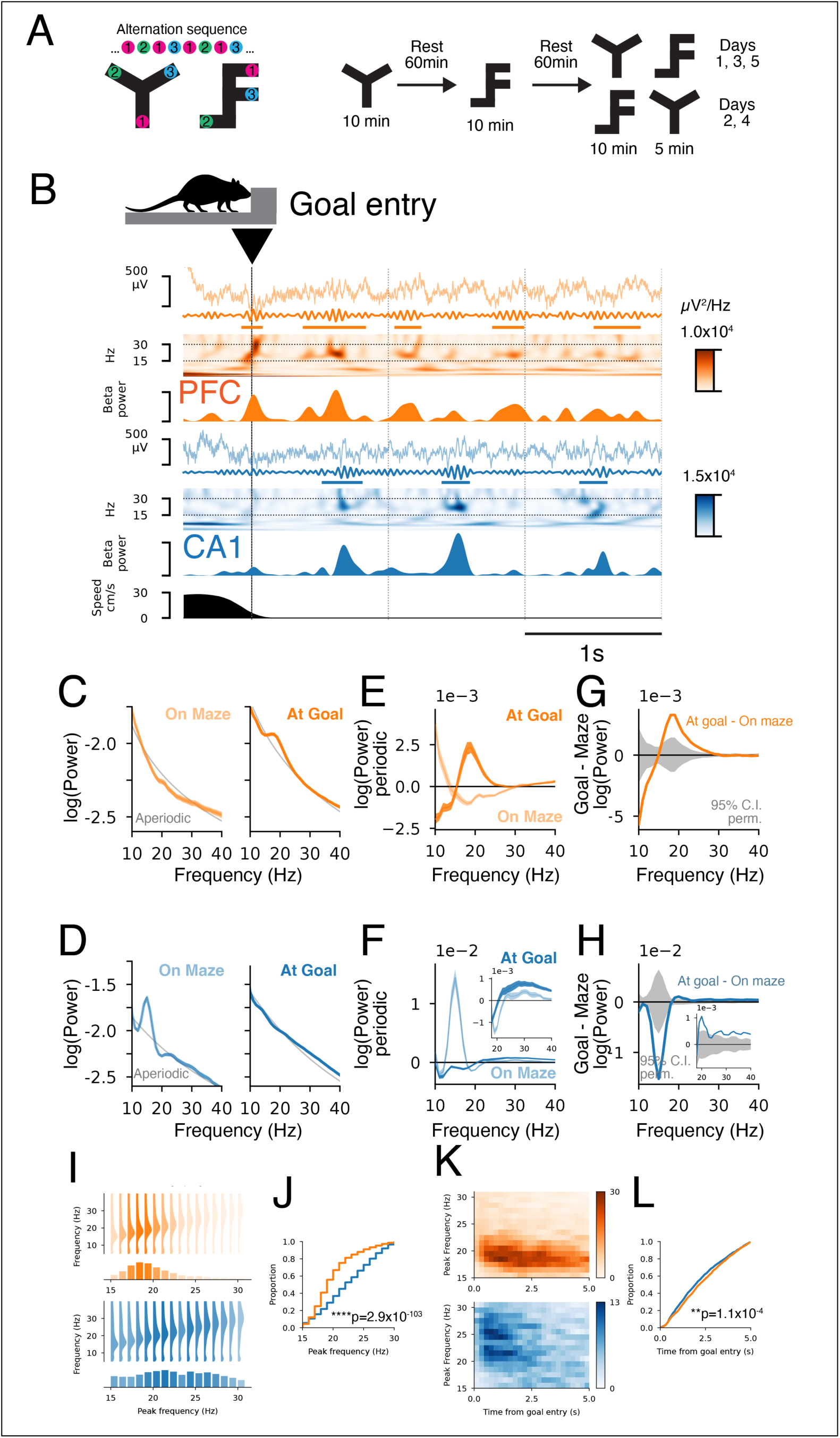
Distinct PFC and CA1 beta-frequency activity after goal location entry during spatial navigation tasks. **A.** Spatial learning tasks on Y- and F-shaped mazes with spatial alternation rules. Animals were trained on the mazes across interleaved sessions. Reward locations (1, 2 and 3) are indicated. The order of visits required to ensure reward on every trial is shown above the maze schematics. The daily training schedule shows the duration, order and separation between sessions. **B.** Example PFC (orange) and CA1 (blue) LFP traces aligned to goal entry (black vertical line). Goal entry is the time when the animal reaches the reward delivery device at the end of each maze arm and breaks the infrared beam. This time is registered by an infrared sensor on the device. The beta-frequency-filtered LFP is shown below the raw LFP. The heatmap shows the corresponding continuous wavelet transform spectrogram (<40 Hz). An extended frequency spectrogram is shown in Fig. 1-2. Beta bursts are indicated by horizontal lines. Dotted vertical lines mark 1 s intervals. **C.** PFC PSD (mean±SEM) for time intervals when the animal is running on the maze (On Maze) (left) or is immobile at the goal (At Goal) (right). The aperiodic component is shown in gray. An extended frequency version is shown in Fig. 1-3. **D.** CA1 power spectral density (mean±SEM) for time intervals when the animal is running on the maze (left) or is immobile at the goal (right). The aperiodic component is shown in gray. An extended frequency version is shown in Fig. 1-3. The peak marked by * in the CA1 PSD at ∼15 Hz corresponds to the first harmonic of theta (Scheffer-Teixeira & Tort, 2016). **E.** PFC PSDs with aperiodic component subtracted. **F.** CA1 PSDs with aperiodic component subtracted. The beta-frequency range is shown in the inset. **G.** PFC difference between goal and maze PSDs (orange) with 95% confidence interval shown in gray. The beta-frequency range is shown in the inset. **H.** CA1 difference between goal and maze PSDs (blue) with 95% confidence interval shown in gray. **I.** Frequency profile for beta bursts grouped by peak power frequency for PFC (orange) and CA1 (blue). Top: the vertical histogram of the mean frequency profiles for bursts at each peak power frequency. Bottom: histograms with the count distribution for bursts within 15-30 Hz, grouped by peak power frequency. Shading corresponds to the probability value at each frequency. Bursts occurring within 5 s after goal location entry are included. **J.** Cumulative distribution of peak burst frequencies for PFC (orange) and CA1 (blue). Kolmogorov-Smirnov test, p=2.9×10^-103^. **K.** Distribution of burst times relative to goal location entry grouped by peak frequency for PFC (orange) and CA1 (blue). **L.** Cumulative distribution of burst occurrence relative to goal location entry for PFC (orange) and CA1 (blue). Kolmogorov-Smirnov test, p=1.1×10^-4^.

Although the PSD in CA1 did not show a peak in the beta-frequency range like the PFC, the continuous wavelet transform from single trials showed that CA1 beta power increased in bursts (Fig. 1 B), which is consistent with reported properties of beta oscillations (Sherman et al., 2016; Shin et al., 2017). To characterize the properties of beta-frequency bursts in PFC and CA1, we extracted burst times from the continuous wavelet transform (Fig. 1-5, Methods). We found the spectral properties of PFC and CA1 bursts differed significantly. PFC bursts were tightly distributed in frequency (median peak frequency: 19 Hz), whereas the peak frequencies of CA1 bursts were distributed across a wider frequency range (median peak frequency: 22 Hz) (Fig. 1 I-J, Fig. 1-3 and Fig. 1-4). This explained the peak in the PFC PSD and the lack of such a peak in the CA1 PSD. The burst amplitude profiles also differed between the two regions. In PFC, lower-frequency bursts tended to be higher in amplitude, whereas CA1 bursts were more uniform in size (Fig. 1-6 A). Burst durations were similar across the two regions (Fig. 1-6 B).

The timing of bursts relative to goal entry also differed between PFC and CA1 (Fig. 1 K and L). Bursts tended to start approximately 400 ms after goal location entry, but CA1 bursts were more concentrated within the first two seconds compared with PFC bursts. The heatmap shows a clear difference in the burst frequency distribution between the two regions and their distribution over time. Although beta is generally defined as oscillations in the 15-30 Hz range, it has been separated into a lower-frequency beta1 (∼15 Hz) and a higher-frequency beta2 (20-30 Hz), based on work in cortical regions (Baker et al., 1997; MacKay & Mendonça, 1995; Roopun et al., 2008; Roopun et al., 2006). However, the frequency of PFC beta bursts described here is higher than beta1 and lower than beta2. The frequency of CA1 beta bursts identified here (∼22 Hz) is consistent with beta2 previously described in the hippocampus (Berke et al., 2008; Igarashi et al., 2014; Iwasaki et al., 2021). These results indicate that PFC and CA1 beta bursts at the goal have distinct spectral and temporal properties.

Beta oscillations are hypothesized to coordinate neural activity across brain regions (Spitzer & Haegens, 2017) to support cognitive processes including sensory-guided decisions and reinstatement of context-dependent memory patterns (Igarashi et al., 2014; Jayachandran et al., 2023; Kay & Freeman, 1998; Symanski et al., 2022). We therefore asked whether PFC and CA1 beta oscillations are coordinated at the goal. If so, we would expect coherence in the beta-frequency range between PFC and CA1, and similar burst timing in both regions. Visual inspection of LFP signals from PFC and CA1 tetrodes showed that some beta bursts were coincident, while others appeared independent in time (Fig. 2 A). To quantify beta-frequency coordination between regions, we first calculated the LFP coherence between tetrodes within each region as well as across regions, for periods when the animal was at the goal. We found higher coherence within PFC or CA1 compared to coherence between PFC and CA1 (Fig. 2 B-C, Table 2-1). The PFC coherence within the beta range showed a peak centered at approximately 20 Hz, whereas the CA1 coherence was similar across the entire beta-frequency range. The across-region coherence was significantly weaker but still showed a small peak at ∼20 Hz. Interestingly, coherence was stronger within each PFC hemisphere than across hemispheres, indicating beta dynamics are more coordinated locally within each hemisphere (Fig. 2-1 A-D). There was similar coherence for ipsi- or contralateral PFC and CA1 comparisons (Fig. 2-1 E-G). We then analyzed burst-wise timing and found beta bursts were more coincident within each region compared with across regions (Fig. 2 D-E). The cross-correlation of burst peak times showed higher values around 0 s lag within each brain region compared with the cross-correlation between PFC and CA1. Comparing across hemispheres, PFC burst timing was more coincident within each hemisphere than across (Fig. 2-1 H-K) but there was weak coincidence with CA1 irrespective of laterality (Fig. 2-1 L-N).

**Figure 2.**
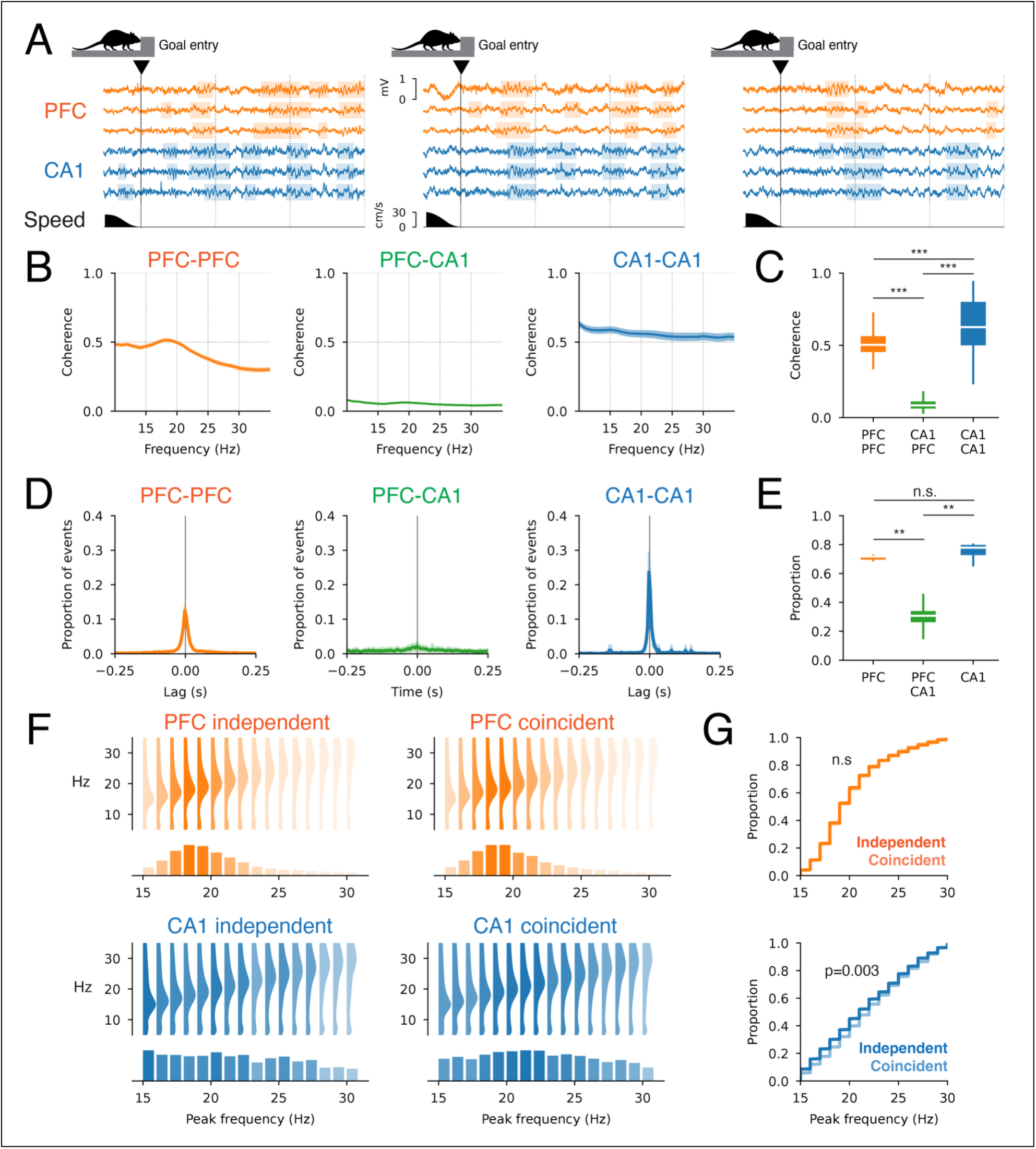
Beta bursts occur independently in both regions. **A.** Beta-frequency bursts (shaded) detected across multiple electrodes in PFC (orange) and CA1 (blue) relative to goal location entry (black vertical line). The speed profile is shown in black. Three example trials are shown. **B.** Coherence (mean±SEM) within PFC (orange), between PFC and CA1 (green), or within CA1 (blue) for periods at the goal location. **C.** Mean coherence (15-30 Hz) across the groups. Pairwise rank-sum test with Benjamini-Hochberg false discovery rate correction: p<0.001 for all pairs. **D.** Cross-correlation (mean±SD) of burst peak times for bursts detected on tetrodes within PFC (orange), across PFC-CA1 (green) and within CA1 (blue). **E.** Proportion of the cross-correlation density within ±50 ms lag for PFC (orange), across PFC-CA1 (green) and within CA1 (blue). Pairwise rank-sum test with Benjamini-Hochberg false discovery rate correction: PFC versus PFC-CA1 (p=0.003**), PFC versus CA1 (p=0.24n.s.) and PFC-CA1 versus CA1 (p=0.003**). **F. Frequenc**y distribution of bursts by peak frequency for independent and coincident bursts in each region. **G.** Cumulative distribution of burst peak frequency for coincident and independent bursts. Kolmogorov-Smirnov test: PFC (p=0.96n.s.) and CA1 (p=0.002**).

Our results so far suggest beta-frequency oscillations are distinct across PFC and CA1 based on their frequency distributions and low burst coincidence. Despite these differences, it is possible that a subset of coincident PFC and CA1 bursts could coordinate activity across the two regions. This hypothesis predicts that coincident bursts have similar frequency distributions and are more coherent between the two regions. We therefore classified each burst as “coincident” across regions if it overlapped in time with a burst in the other brain region. Bursts that did not overlap were classified as “independent” across the regions. Based on this method, we asked whether coincident bursts were more closely matched in their frequency distribution across the regions compared with independent bursts. However, this was not the case, as coincident bursts in CA1 were higher in peak frequency than coincident bursts in PFC (Fig. 2 F-G). The difference in frequency distribution for the coincident bursts provides evidence against a common beta-frequency range coordinating PFC and CA1, since there was a significant frequency mismatch. We confirmed our classification of independent and coincident bursts using burst-aligned power and coherence analysis (Fig. 2-2). Independent bursts in one region were not accompanied by significant increases in beta power in the other region (Fig. 2-2 A-B). Coincident bursts, by definition, showed the expected increase in power in both regions. We found a similar pattern for coherence (Fig. 2-2 D-E). During independent bursts, coherence increased within the region with the burst but not in the other region. Coherence in both regions increased during coincident bursts. Coherence across regions also increased for coincident bursts more than independent bursts (Fig. 2-2 G-H); however, the coherence value across brain regions was lower in comparison with coherence within each region (Fig. 2-2 E vs. Fig. 2-2 H).

### Experience-dependent goal location beta dynamics

Having established the properties of beta oscillations in PFC and CA1, we next asked how goal-related hippocampal-neocortical beta oscillations changed across task learning. We therefore determined the relationship between beta properties in PFC and CA1 with either days of training or performance in each task. Animals performed better on the Y alternation task than the F alternation task over 5 days (Fig. 3 A). By day 5, animals showed clear evidence of learning the Y maze task, with performance exceeding 80%. Animals struggled to learn the F maze, with performance remaining close to 50%, which is at chance for the alternation task, making the F maze task a non-learning control condition. Although the performance on the F maze task showed a statistically significant increase across days, the low marginal R^2^ indicates that the change across days was weak. This performance difference allowed us to compare beta dynamics between tasks with distinct learning outcomes under comparable levels of familiarity, since the animals were trained on both tasks over the same 5-day period. In both tasks, PFC beta burst power increased over the five days, whereas CA1 beta burst power decreased (Fig. 3 B). We repeated the same set of analyses by partitioning the data based on performance quintiles (Fig. 3 C-D), which allowed us to directly relate beta activity properties with performance. PFC beta burst power increased with improved performance. CA1 beta burst power decreased with improvements in performance. These changes were observed on the Y maze task, which animals learned, but not on the F maze, which the animals failed to learn. These effects were not due to changes in the animal’s movement over days since we ensured the animal was immobile at the time of the bursts (Fig. 3-1). Thus, we found complementary changes in beta dynamics in PFC and CA1 with familiarity and performance.

**Figure 3.**
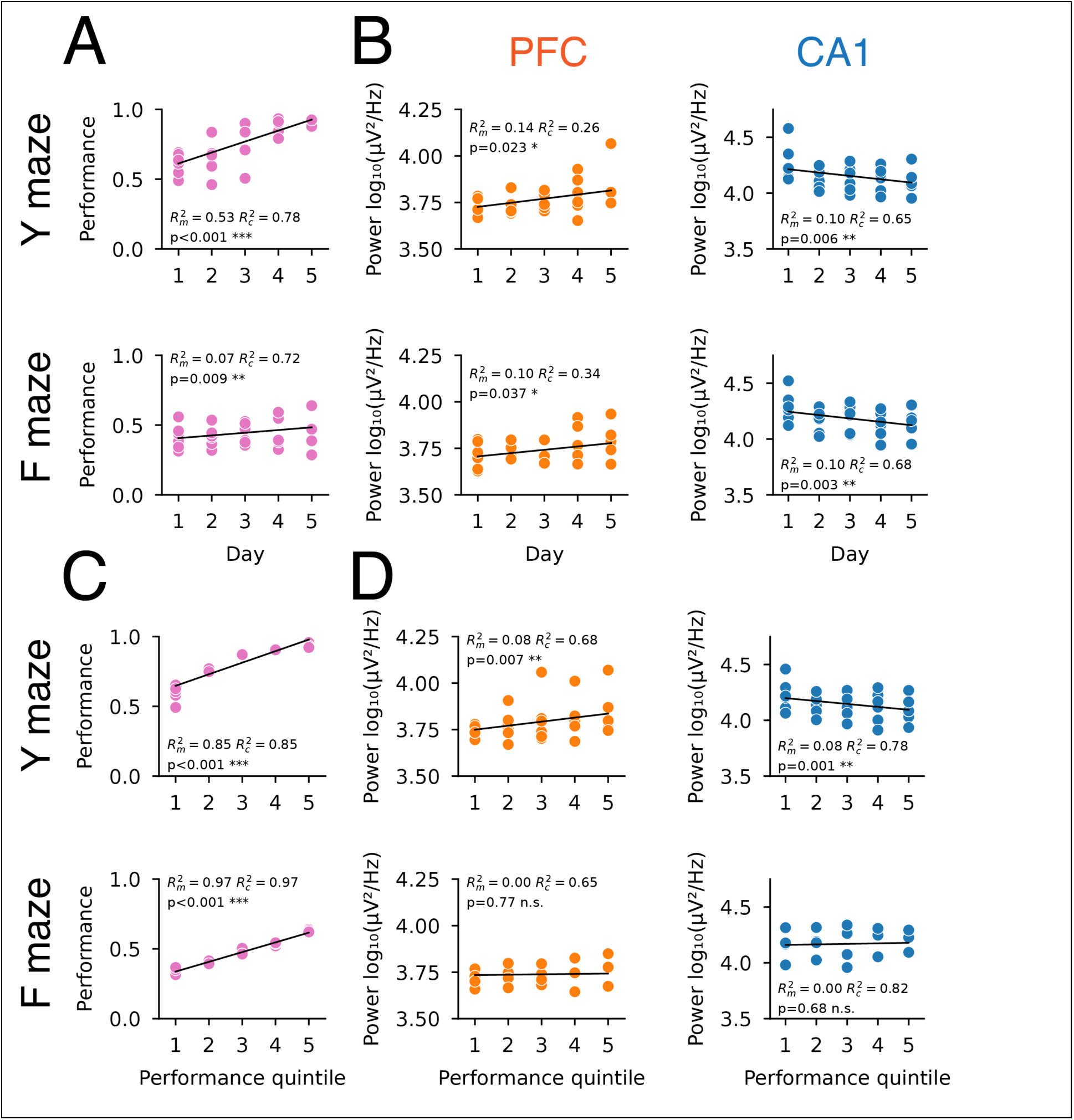
Distinct changes in PFC and CA1 burst properties across days of learning. **A.** Performance in each maze task by day. Linear model with fixed (day) and random (animal) effects. The marginal R^2^ (R^2^_m_) and conditional R^2^ (R^2^_c_) are shown. **B.** Burst power in each maze task by day. Linear model with fixed (day) and random (animal) effects. **C.** Performance in each maze task by performance quintile. Linear model with fixed (performance quintile) and random (animal) effects. **D.** Burst power in each maze task by performance quintile. Linear model with fixed (performance quintile) and random (animal) effects.

In addition to changes in the strength of beta bursts across days and with improved performance, we asked whether beta coordination between PFC and CA1 changed as a function of day or performance. We found weak effects, indicating PFC and CA1 beta coherence (Fig. 3-2 B, E, H and K) and burst coincidence (Fig. 3-3 B, E, H and K) did not change as a function of day or performance. Although some of the linear mixed-effects models produced statistically significant results, the effect of day or performance on coherence or burst coincidence was weak given the marginal R^2^ was very low compared with the conditional R^2^ (Fig. 3-2 B, F and J). The most robust effect was in burst coincidence within CA1, which decreased across days and with improved performance in the Y maze task (Fig. 3-3 C and I).

We next identified temporal changes in beta-frequency-modulated processes in PFC and CA1 with learning. Over days of training, the timing of beta bursts relative to goal entry on unrewarded trials remained unchanged in both PFC and CA1 and on both mazes (Fig. 4 A). However, bursts on rewarded trials shifted closer to the time of goal entry for PFC on the Y maze and CA1 on both mazes (Fig. 4 A-B). Further, burst timing differed between unrewarded and rewarded trials, with bursts on rewarded trials occurring later than those on unrewarded trials relative to goal entry (Fig. 4 A-B, right columns). Similar trends were observed when burst timing was correlated with performance (Fig. 4 C-D). We ensured these trends were not due to differences in the movement of the animal at the goal for rewarded or unrewarded trials (Fig. 4-1). These timing properties were observed in Y and F mazes (Fig. 4-2). Interestingly, the PFC burst peak frequency was lower on rewarded compared with unrewarded trials, which was not the case for CA1 bursts (Fig. 4-3). Across the 5 days, the burst peak frequency also became more consistent, but only when performance improved. The variance in PFC and CA1 burst frequency decreased over days of training for the Y but not the F maze (Fig. 4-4 A and C). We ensured this result was not due to differences in burst event counts across days (Fig. 4-4 B and D). These results demonstrate that beta-frequency modulation in PFC and CA1 has distinct correlates with performance across the five days of training, suggesting changes to beta-frequency activity reflect learning-related dynamics in PFC and CA1 networks.

**Figure 4.**
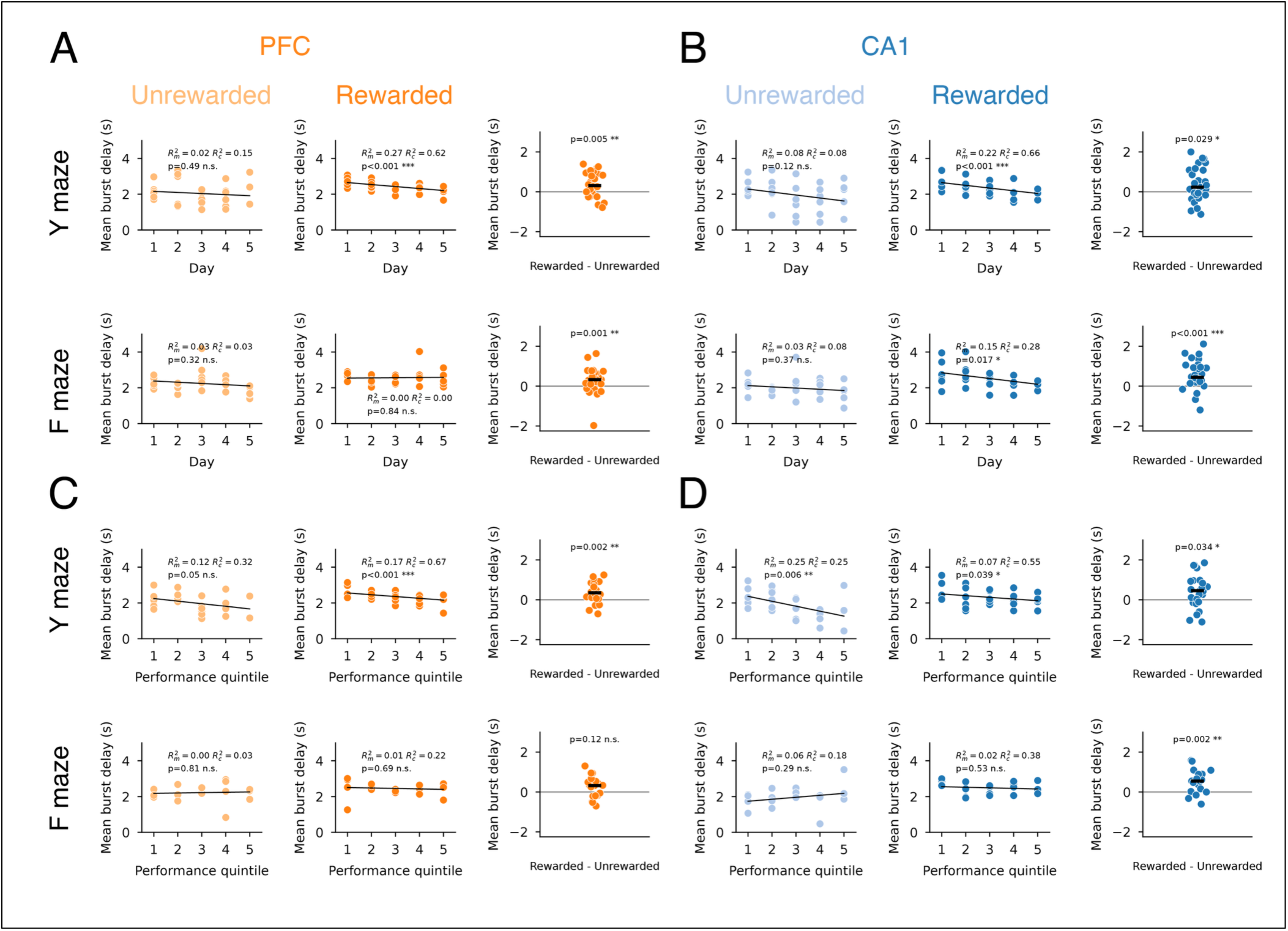
Bursts occur sooner after goal entry over days of training. **A.** PFC burst delay distribution relative to goal entry versus days for unrewarded (left column) or rewarded (center column) trials, for the two maze tasks (top or bottom row), and their comparison (right column). **B.** CA1 burst delay distribution relative to goal entry versus days. **C.** PFC burst delay distribution relative to goal entry versus performance. **D.** CA1 burst delay distribution relative to goal entry versus performance.

### Spiking entrainment to goal location beta oscillations

We next asked how PFC and CA1 spiking activity is modulated at the beta-frequency range. Example PFC and CA1 cells (Fig. 5 A-B, Tables 5-1, 5-2 and 5-3) showed spike phase-locking preference for the beta-frequency range (Fig. 5 C-D and Table 5-4). We show example phase-aligned spike rasters to illustrate each cell’s phase locking at each goal location (Fig. 5 E-F), as well as the corresponding spiking pattern relative to goal entry (Fig. 5 G-H). Both PFC (23%) and CA1 (32%) populations showed significant phase locking to beta within each region (Fig. 5 I-N). This finding was based on two analysis methods, the Rayleigh test and the von Mises fit (see Methods: Spike phase-locking analysis). The PFC and CA1 cell populations showed phase preferences to the trough of local beta, with PFC cells biased toward spiking during the rising phases, whereas CA1 cells were slightly biased toward the falling phases (Fig. 5 K and N). These properties were highly similar between the two maze tasks (Fig. 5-1). Further, we found cells in each region showed higher phase locking to local beta compared with remote beta (Fig. 5 O-R). This indicates that spiking in each region is modulated preferentially by local beta oscillations rather than remote beta oscillations. These results provide further evidence that the PFC and CA1 are independently modulated at the beta frequency during the goal period.

**Figure 5.**
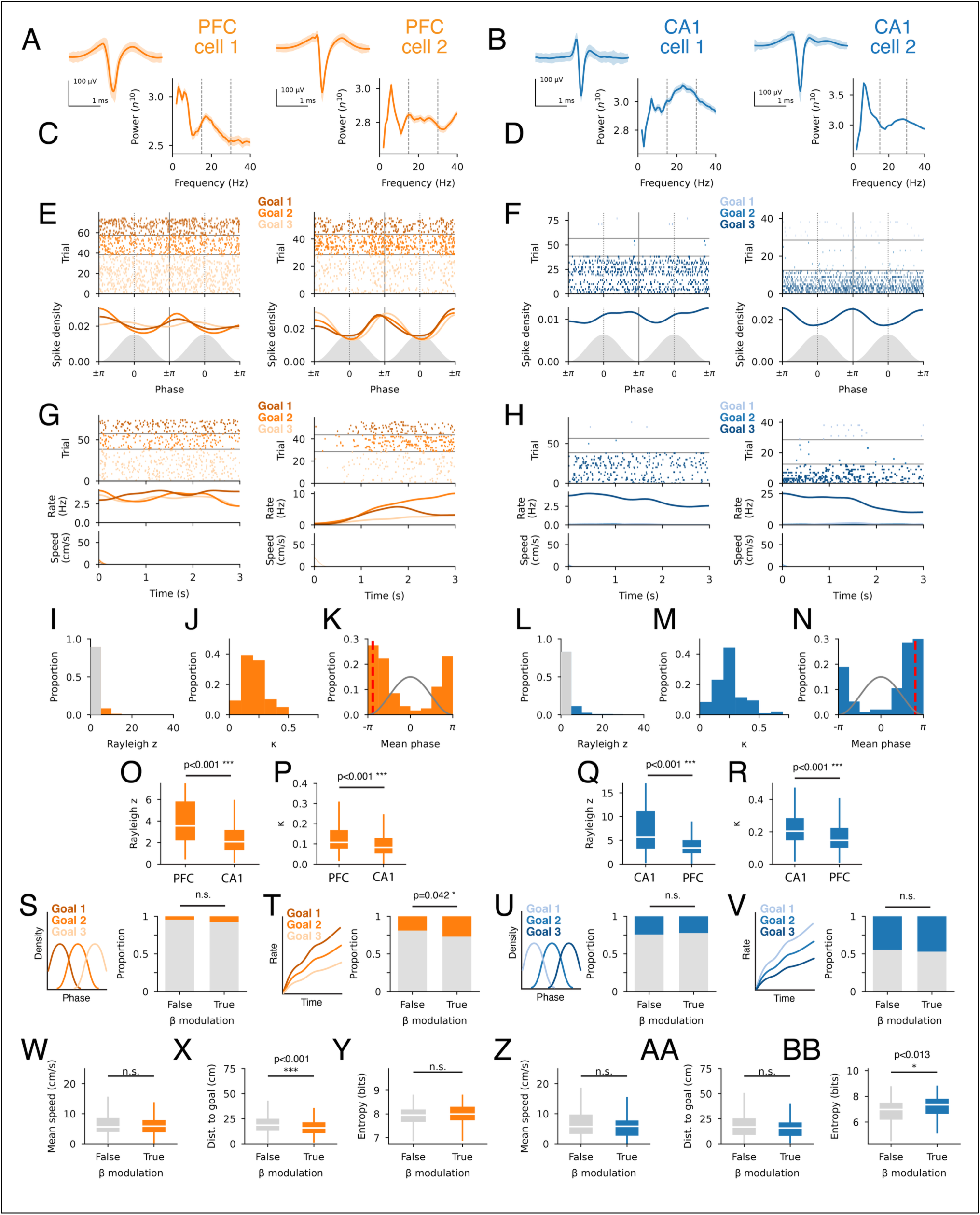
PFC and CA1 cells show stronger spiking modulation to local beta. **A.** Spike waveforms for example PFC cells. **B.** Spike waveforms for example CA1 cells. **C.** Spike-aligned power spectrum for example PFC cells. **D.** Spike-aligned power spectrum for example CA1 cells. **E.** Spike beta-phase-locking raster and histogram for example PFC cells. Spike raster is arranged by trial and goal location (color shading). The corresponding mean histogram for each goal location is shown below. The phase of beta is shown in gray. **F.** Spike beta-phase-locking raster and histogram for example CA1 cells. **G.** Goal location-aligned spiking raster for the example PFC cells. The mean firing rate for each goal location is shown below the spike raster. The corresponding speed at each goal is in the lower row. **H.** Goal location-aligned spiking raster for the example CA1 cells. **I.** Distribution of Rayleigh z for PFC. **J.** Distribution of k from a von Mises distribution fit for significantly beta-modulated cells (Rayleigh p <0.05 in panel I) in PFC. **K.** Distribution of phase preference for significantly beta-modulated cells in PFC. Red dotted line shows the mean population phase preference. The gray line indicates the phase of beta. **L.** Distribution of Rayleigh z for CA1. **M.** Distribution of k from a von Mises distribution fit for significantly beta-modulated cells (Rayleigh p <0.05 in panel L) in CA1. **N.** Distribution of phase preference for significantly beta-modulated cells in CA1. **O. P**FC beta-phase-locking strength using the Rayleigh test (Rayleigh z) to PFC or CA1 beta. Wilcoxon signed-rank test: p=5.4×10^-50^. **P.** PFC beta-phase-locking strength using the von Mises fit (kappa) to PFC or CA1 beta. Wilcoxon signed-rank test: p=5.6×10^-43^. **Q.** CA1 beta-phase-locking strength using the Rayleigh test (Rayleigh z) to CA1 or PFC beta. Wilcoxon signed-rank test: p=5.3×10^-55^. **R.** CA1 beta-phase-locking strength using the von Mises fit (kappa) to CA1 or PFC beta. Wilcoxon signed-rank test: p=1.5×10^-45^. **S.** Goal location classification based on mean phase preference at each goal (schematic on the left). Bar plot shows the proportion of the PFC population (non-beta or beta-modulated) that distinguished goal identity (orange) or did not (gray) based on mean phase preference at each goal. Fisher’s exact test: p=0.19. **T.** Goal location classification based on mean firing rate at each goal (schematic on the left). Bar plot shows the proportion of the PFC population (non-beta or beta-modulated) that distinguished goal identity (orange) or did not (gray) based on mean firing rate at each goal. Fisher’s exact test: p=0.042. **U.** Bar plot shows the proportion of the CA1 population (non-beta or beta-modulated) that distinguished goal identity (blue) or did not (gray) based on mean phase preference at each goal. Fisher’s exact test: p=0.70. **V.** Bar plot shows the proportion of the CA1 population (non-beta or beta-modulated) that distinguished goal identity (blue) or did not (gray) based on mean firing rate at each goal. Fisher’s exact test: p=0.66. **W.** Mean spiking speed for PFC cells based on beta modulation. Wilcoxon rank-sum test: p=0.41. **X.** Spiking distance to nearest goal for PFC cells based on beta modulation. Wilcoxon rank-sum test: p=0.0006. **Y.** Spatial firing entropy for PFC cells based on beta modulation. Wilcoxon rank-sum test: p=0.18. **Z.** Mean spiking speed for CA1 cells based on beta modulation. Wilcoxon rank-sum test: p=0.29. **AA.** Spiking distance to nearest goal for CA1 cells based on beta modulation. Wilcoxon rank-sum test: p=0.13. **BB.** Spatial firing entropy for CA1 cells based on beta modulation. Wilcoxon rank-sum test: p=0.013.

Given PFC and CA1 cells showed clear goal-entry-modulated spiking (Fig. 5-2), we next determined whether beta-phase-locked or non-phase-locked cells had distinct task correlates. We first asked whether phase-locking preference or firing rate distinguished between the goal locations, which could suggest phase or rate coding of goal identity. Using a Gaussian Naïve Bayes model to classify goal identity based on the beta phase for spikes, we found the phase of PFC spiking was a poor differentiator of goal identity (Fig. 5 S). In contrast, PFC firing rate differed between goals, which was more likely for beta-phase-locked cells than non-phase-locked cells (Fig. 5 T). For CA1, a larger proportion of the cell population differentiated the goals based on spiking phase (Fig. 5 U). We note that for CA1, even cells that were classified as not modulated by beta showed differences in phase locking between goals. This is explained by our beta-phase modulation classification being based on spiking from all goals rather than individual goals, where a non-beta-modulated cell could show spike phase preference at one of the goals. Similarly, for firing rate, a large proportion of CA1 cells had different firing rates at each goal, and the proportions were similar between phase-locked and non-phase-locked cells (Fig. 5 V). In additional analyses of task firing correlates, we found PFC beta-modulated cells fired closer to goals compared with non-phase-locked cells (Fig. 5 X), an effect driven by the F maze (Fig. 5-3 B). PFC beta-phase-locked cells had similar spatial firing entropy compared with non-phase-locked cells (Fig. 5 Y and Fig. 5-3 C, left two columns). CA1 beta-phase-locked cells had similar proximity to goal firing compared with non-beta-phase-locked cells (Fig. 5 AA) but had higher spatial firing entropy than non-beta-phase-locked cells only on the Y maze (Fig. 5 BB and Fig. 5-3 C, right two columns). Beta-phase-locking status did not identify differences in the speed preference of the cells (Fig. 5 W and Z, and Fig. 5-3 A).

### Distinct temporal coordination between beta oscillations and hippocampal SWRs

We next asked how changes in PFC and CA1 beta oscillations are related to another prominent neural process supporting learning: awake hippocampal SWRs. Hippocampal SWRs also occur at goal locations (Ambrose et al., 2016; Carey et al., 2019; Diba & Buzsaki, 2007; Foster & Wilson, 2006; Kleinman & Foster, 2025; Singer & Frank, 2009) and are hypothesized to coordinate distributed representations (Jadhav et al., 2016; Shin et al., 2019; Tang et al., 2021; Yu et al., 2017; Yu et al., 2018) to support the formation and use of memory during spatial learning (Cheng & Frank, 2008; Deceuninck & Kloosterman, 2024; Dupret et al., 2010; Gupta et al., 2010; Jadhav et al., 2012; Pfeiffer & Foster, 2013; Singer et al., 2013; Singer & Frank, 2009). Since both beta and SWRs occur at goal locations, we asked how beta-frequency dynamics within PFC and CA1 networks are related to hippocampal SWR dynamics. Consistent with previous findings, we found SWRs tended to occur several seconds after the animal reached the goal (Fig. 6 A-B) (Ambrose et al., 2016; Carey et al., 2019; Diba & Buzsaki, 2007; Foster & Wilson, 2006; Kleinman & Foster, 2025; Singer & Frank, 2009). SWR power (Fig. 6-1 A) and rate (Fig. 6-1 B), but not duration (Fig. 6-1 C), decreased across days, consistent with prior findings (Cheng & Frank, 2008). We ensured the animals were immobile (Fig. 6-1 D), and the goal visit count (Fig. 6-1 E) and duration (Fig. 6-1 F) were comparable over days.

**Figure 6.**
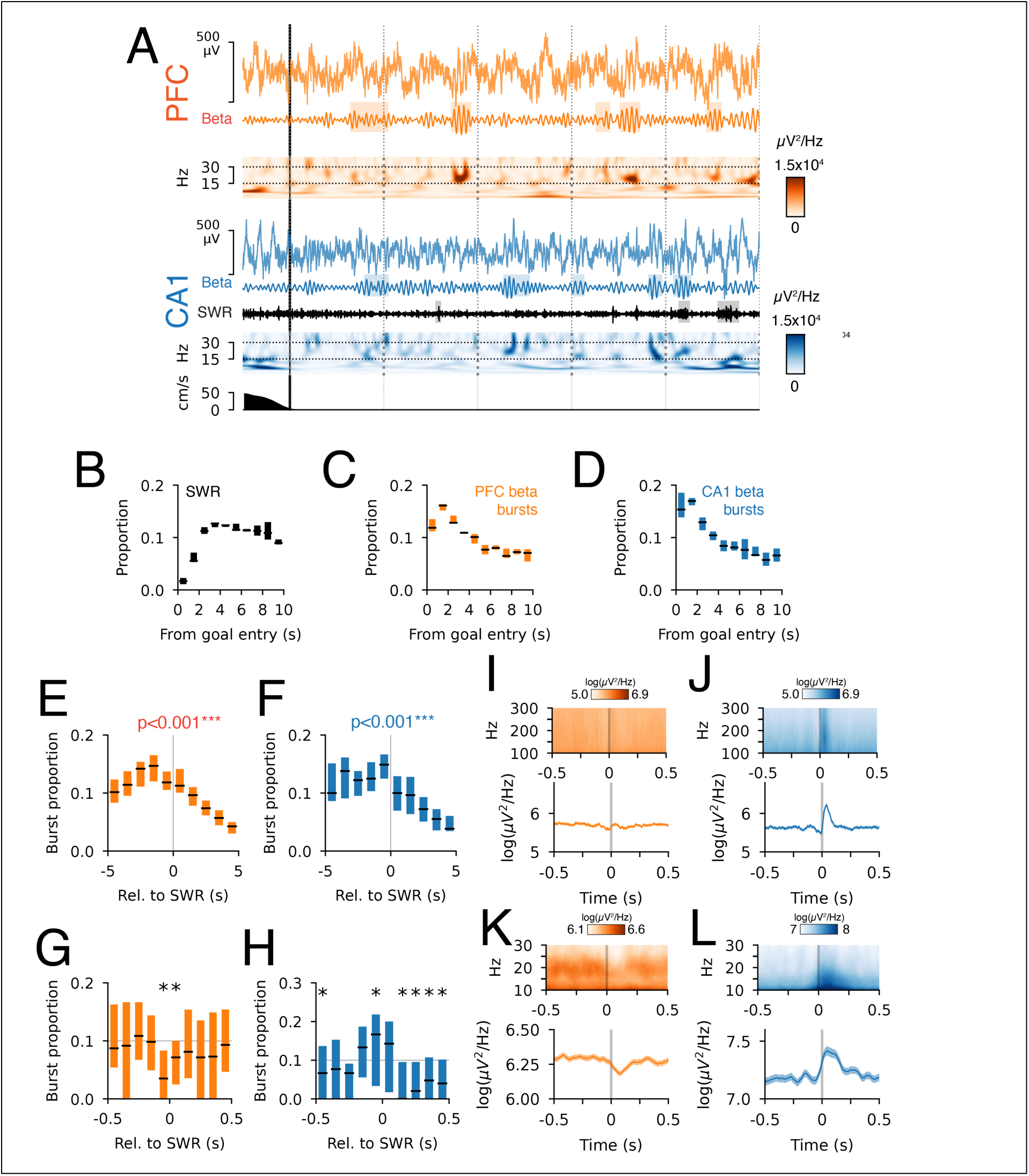
Goal location beta bursts and SWRs are coordinated at multiple timescales. **A.** PFC (orange) and CA1 (blue) LFP relative to goal entry (black vertical line). Beta-frequency-filtered signal is shown below the LFP. Ripple frequency (150-250 Hz) filtered CA1 signal is shown in black. The corresponding speed of the animal is shown in the bottom row. **B.** Goal-entry-aligned histogram for CA1 SWRs. Median and interquartile range are shown. **C.** Goal-entry-aligned histogram for PFC beta bursts. **D.** Goal-entry-aligned histogram for CA1 beta bursts. **E.** Cross-correlation between PFC beta bursts and CA1 SWRs (±5 s, 1 s bins). Beta bursts occur before SWRs, Wilcoxon signed-rank test: p=5.2×10^-8^ for burst time relative to SWRs. **F.** Cross-correlation between CA1 beta bursts and CA1 SWRs (±5 s, 1 s bins). Beta bursts occur before SWRs, Wilcoxon signed-rank test: p=7.1×10^-8^ for burst time relative to SWRs. **G.** Cross-correlation between PFC beta bursts and CA1 SWRs (±500 ms, 100 ms bins). Bins with significant deviations (with Benjamini-Hochberg false discovery rate correction) from the mean are indicated with *. **H.** Cross-correlation between CA1 beta bursts and CA1 SWRs (±500 ms, 100 ms bins). Bins with significant deviations (with Benjamini-Hochberg false discovery rate correction) from the mean are indicated with *. **I.** SWR-aligned PFC spectrogram (100-300 Hz) and corresponding ripple band power (150-250 Hz). **J.** SWR-aligned CA1 spectrogram (100-300 Hz) and corresponding ripple band power (150-250 Hz). **K.** SWR-aligned PFC spectrogram (10-30 Hz) and corresponding beta-band power (15-30 Hz). **L.** SWR-aligned CA1 spectrogram (10-30 Hz) and corresponding beta-band power (15-30 Hz).

We then found distinct temporal coordination patterns between beta bursts and SWRs that emerged on two timescales. In contrast to the delayed onset of hippocampal SWRs, beta bursts in PFC and CA1 occurred closer to the time of goal entry (Fig. 6 C-D); thus, on the timescale of seconds, beta bursts in each region occurred earlier than SWRs. We confirmed this relationship occurred on a single-trial timescale by computing the cross-correlation between the timing of beta bursts in each region and SWRs (Fig. 6 E-F). On this timescale, the beta bursts continued to occur earlier than SWRs throughout days of training (Fig. 6-2 A and C). We next determined whether beta bursts and SWRs were temporally coordinated on a finer timescale. On a shorter timescale of tens of milliseconds, each brain region had the opposite coordination pattern between beta bursts and hippocampal SWRs (Fig. 6 G-H). PFC beta bursts were less frequent at the time of hippocampal SWRs (Fig. 6 G), as shown by the dip in the cross-correlation histogram at 0 s lag, whereas CA1 beta bursts and SWRs were more coincident (Fig. 6 H), as shown by the peak at 0 s lag. Over days of training, this pattern persisted in PFC, whereas CA1 beta bursts became less coincident with SWRs (Fig. 6-2 B and D), suggesting an experience-dependent change specific to CA1.

This pattern of beta burst and SWR coordination was further supported by SWR-aligned spectrograms (Fig. 6 I-L). PFC beta-band power showed a decrease at the time of hippocampal SWRs (Fig. 6 K), whereas CA1 beta-band power showed an increase (Fig. 6 L). These patterns remained consistent when the Y and F maze data were analyzed separately (Fig. 6-3). It is possible that the CA1 beta-band power around the time of SWRs results from accompanying sharp waves (Buzsaki et al., 1983); however, our burst timing analysis suggests beta and SWRs are distinct. Although CA1 beta bursts can coincide with SWRs, the cross-correlation shows some beta bursts can occur earlier than SWRs (Fig. 6 H). We also found SWR band power showed phase coupling to CA1 but not PFC beta (Fig. 6-4). In CA1, SWR power was maximum at the peak of local beta (Fig. 6-4 C-D). SWRs and CA1 beta bursts can coincide and SWR amplitude is phase-coupled to local beta oscillations. Thus, the temporal coordination between beta bursts and SWRs occurs on the second and millisecond timescales and supports both markers being complementary neural signatures for learning-related processes.

### Beta-phase locking and SWR modulation identify a unique PFC subpopulation

Since our results suggest that goal location beta-frequency activity and SWRs correspond to temporally coordinated processes, do they modulate distinct populations of PFC and CA1 cells? In PFC and CA1, we found cells modulated by both SWRs and beta (during the 5 s after goal entry) (Fig. 7 A, B, G and H). In PFC, 22% of cells showed modulation to beta and ∼30% showed modulation to SWRs. The proportion of PFC cells that were modulated by both SWR and beta (8%) was not significantly different from the expected proportion (6.6%) under the assumption that the population is independently modulated by SWR and beta (30%×22%) (Fisher’s exact test, p=0.09). We found PFC cells modulated by both beta and SWRs (SWR^+^β^+^) had distinct firing properties. The SWR^+^β^+^ population had a positive relationship between the spike beta-phase preference and the direction of SWR modulation (Fig. 7 C). PFC cells with spiking preference for the falling phases of the beta cycle tended to show spiking inhibition during SWRs (negative SWR-modulation index) (Fig. 7 A, bottom row), whereas cells with spiking preference for the rising phases of beta tended to be excited during SWRs (positive SWR-modulation index) (Fig. 7 A, top row). As a control, this relationship was not observed for PFC cells that were not modulated by beta but modulated by SWRs (SWR^+^β^-^) (Fig. 7 C, lower panel, Fig. 7-1 A). The SWR^+^β^+^ PFC cell population was further distinguished from the SWR^+^β^-^ population by its strong positive relationship with goal location firing. Negatively SWR-modulated cells were active closer to goals whereas positively SWR-modulated cells were active further away from goals (Fig. 7 D, upper panel). This relationship was absent for SWR^+^β^-^ cells (Fig. 7 D, lower panel). SWR-modulated PFC cells showed a positive relationship with speed, where negatively modulated cells spiked when the animal was at a lower speed and positively modulated cells spiked when the animal was moving faster (Fig. 7 E), which is consistent with prior work (Jadhav et al., 2016; Yu et al., 2017). This relationship did not depend on beta modulation status. Similarly, SWR-modulated PFC cells had comparable spatial firing entropy irrespective of beta modulation status (Fig. 7 F). Additional task correlate analysis showed that beta modulation did not further segregate SWR-modulated PFC cells based on phase or firing rate coding of goal locations (Fig. 7-2 A and B).

**Figure 7.**
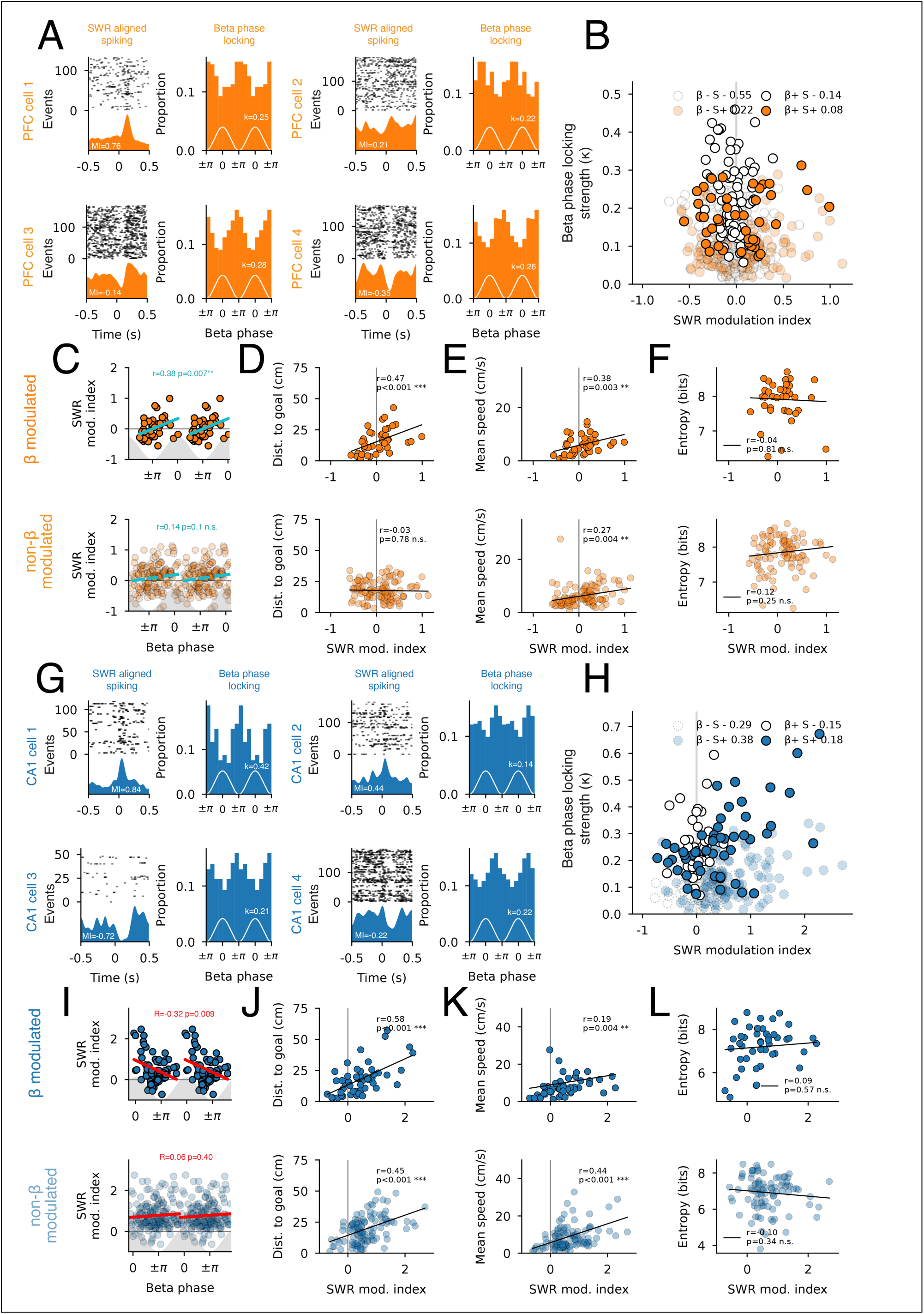
SWR and beta modulation in PFC and CA1. **A.** Four example PFC cells showing SWR-aligned spiking raster and histogram, and spike beta-phase-locking histogram. Top two cells show spiking excitation around SWRs, and bottom two cells show spiking inhibition around SWRs. The SWR-modulation index (MI) is displayed. In the spike beta-phase-locking histogram, the distribution is duplicated to improve visualization. The peaks and troughs for the beta cycle are shown in white and the phase-locking strength (k) is displayed. **B.** PFC beta-modulation strength (k) versus SWR-modulation index. The four combinations of SWR (S) and beta (β) modulation (+: modulated and -: not modulated) and their corresponding proportions are indicated. **C.** PFC SWR-modulation index versus beta-phase-locking preference for beta and SWR-modulated cells (S+β+) (upper) and SWR-modulated but not beta-modulated cells (S+β-) (lower). The regression line is shown in cyan. The gray shading indicates the peak and trough of beta. Two cycles are repeated for clarity. **D.** Spiking distance to nearest goal versus SWR-modulation index. **E.** Spiking speed versus SWR-modulation index. **F.** Spatial firing entropy versus SWR-modulation index. **G.** Four example CA1 cells showing SWR-aligned spiking raster and histogram, and spike beta-phase-locking histogram. **H.** CA1 beta modulation strength (k) versus SWR-modulation index. The four combinations of SWR (S) and beta (β) modulation (+: modulated and -: not modulated) and their corresponding proportions are indicated. **I.** CA1 SWR-modulation index versus beta-phase-locking preference for beta and SWR-modulated cells (S+β+) (upper) and SWR-modulated but not beta-modulated cells (S+β-) (lower). The regression line is shown in red. The gray shading indicates the peak and trough of beta. Two cycles are repeated for clarity. **J.** Spiking distance to nearest goal versus SWR-modulation index. **K.** Spiking speed versus SWR-modulation index. **L.** Spatial firing entropy versus SWR-modulation index.

For CA1, we found modulation patterns distinct from PFC. We repeated these analyses for the CA1 population and found the opposite relationship between beta-phase preference and SWR modulation compared with PFC cells (Fig. 7 I). In CA1, ∼33% of cells showed beta-phase modulation and ∼56% of cells showed modulation to SWRs. The subset of CA1 cells that were modulated by both SWR and beta (SWR^+^β^+^, 18%) was similar to the expected proportion (19%) under the assumption of independence (33%×56%) (Fisher’s exact test, p=0.62). For this population, cells with spiking preference for the falling phases of the beta cycle tended to have a positive SWR-modulation index, indicating stronger excitation during SWRs (Fig. 7 I). As a control, this relationship was not observed for the SWR^+^β^-^ CA1 cells (Fig. 7 I, lower panel, Fig. 7-1 B). Unlike PFC cells, SWR-modulated CA1 cells showed a strong positive correlation between the direction of SWR modulation and spiking distance relative to the goal and speed, irrespective of beta modulation patterns (Fig. 7 J-K). This is consistent with known properties of dorsal CA1 cells, where cells with place fields on the maze leading to goals are reactivated and immobility cells active at goals are suppressed (Jin et al., 2024; Kay et al., 2016; Yu et al., 2017). SWR^+^β^-^ CA1 cells showed a negative correlation with spatial firing entropy; however, the relationship was weak (Fig. 7 L, lower panel). We also determined whether SWR and beta modulation status identified cells that could distinguish goal identity, either based on phase-locking preference or firing rate at goals. However, these were similar irrespective of beta or SWR-modulation status (Fig. 7-2 A-B).

Our results show that beta oscillations modulate a subpopulation of PFC neurons active at the goal, and some of these cells are coordinated with hippocampal SWRs. The PFC subset may not be coordinated with the hippocampus at the beta frequency, but it can still be linked with relevant hippocampal representations during SWRs. This subset of PFC cells expressed a more faithful recapitulation of the task structure relative to goals than the rest of the PFC population, which may represent a functional subpopulation engaged in representing the ongoing task structure.

## Discussion

Our results show beta-frequency oscillations occurred after entry to goal locations in both CA1 and PFC. However, several properties suggest that the two brain regions are weakly coordinated at the beta frequency. First, we found that beta bursts had distinct spectral and temporal properties in each brain region, where PFC beta bursts were lower in frequency than CA1 bursts. Second, PFC burst amplitude increased with performance, whereas CA1 burst amplitude decreased. This was observed when animals learned the task (Y maze) but not when they failed to learn the task (F maze). The pattern of learning-related changes for PFC bursts was also opposite to those of SWRs. Lastly, beta- and SWR-modulation jointly identified a specific subpopulation with distinct task correlates in PFC but not in CA1.

Our results suggest that the neocortex and hippocampus may participate in distinct functional networks at the goal, and beta oscillations may support each region’s interactions with its respective network partners at distinct frequencies within the beta band. This contrasts with other task periods where the PFC and hippocampus are part of the same functional network modulated by beta or other frequencies. For example, during spatial navigation, PFC cells show strong phase locking to hippocampal theta (Hyman et al., 2005; Jones & Wilson, 2005; Siapas et al., 2005; Yu & Frank, 2021; Zielinski et al., 2019), a frequency band that has been hypothesized to reflect coordination of distinct task-related representations across these structures during the execution of cognitive tasks. During olfactory sensory decision making, beta-frequency coherence is observed across neocortical and hippocampal networks, which is thought to coordinate associative memory representations (Fourcaud-Trocmé et al., 2019; Igarashi et al., 2014; Jayachandran et al., 2023; Kay & Freeman, 1998; Martin et al., 2007; Sheriff et al., 2021; Symanski et al., 2022). During awake quiescence, activity in neocortical and hippocampal networks is transiently synchronized during SWRs (Jadhav et al., 2016; Remondes & Wilson, 2015; Shin et al., 2019; Tang et al., 2017; Yu et al., 2017; Yu et al., 2018). Coordination at these frequencies and task periods demonstrates hippocampal-neocortical interactions play a central role in supporting diverse cognitive functions. Our results show that the neocortex and hippocampus may be weakly coupled during the goal period at the beta frequency, and each region may be engaged with its own set of functionally connected networks (Cohen et al., 2011). This was surprising given the extensive coordination across these brain structures during other important task periods. We had expected that stronger coordination during this period would be important for learning-related changes across neocortical-hippocampal networks, similar to SWRs that occur at the goal.

Our results also reveal complementary changes in hippocampal-neocortical beta dynamics across multiple timescales. First, we found an inverse pattern in the hippocampus and PFC during learning, which spans the timescale of days. These results indicate that beta-frequency oscillations in the neocortex and hippocampus are markers of complementary learning-induced changes in these networks, pointing to the contrasting contributions of the hippocampus and neocortex to learning. The hippocampus is hypothesized to play an important role in rapid changes during early learning, whereas the neocortex gradually changes on a longer timescale (Buzsáki, 1996; McClelland et al., 1995). In the hippocampus, SWRs are a neural signature of early learning, where larger amplitude events are observed during novel experience (Cheng & Frank, 2008). With learning, these events become less prominent. We found similar patterns for goal-related beta activity in the hippocampus, which decreased with performance. This mirrors novelty-related beta activity in the hippocampus that occurs during exploration of new environments or objects, which decreases with familiarity (Berke et al., 2008; França et al., 2014; França et al., 2021; Rangel et al., 2015). In contrast to hippocampal beta oscillations, we hypothesize that beta activity in the PFC reflects the reconfigured state for a given task as learning progresses, where beta amplitude and frequency uniformity increase with experience and performance. This increase parallels findings in the olfactory bulb and piriform cortex, where the strength of beta during odor sampling increases with performance proficiency in olfactory sensory decision-making tasks (Cohen et al., 2015; Fourcaud-Trocmé et al., 2019; Gervais et al., 2007; Martin et al., 2007; Martin et al., 2004), in the ventral striatum, where beta strengthens as habits form (Howe et al., 2011), and the emergence of beta oscillations in the basolateral amygdala with the development of reward preference (Amaya et al., 2024).

On the timescale of seconds, we found beta bursts typically preceded hippocampal SWRs after goal entry (Fig. 6 A-F) across days of training (Fig. 6-2 A and C). Beta bursts in PFC and CA1 occurred most frequently within a second after goal entry and then became less frequent. In contrast, hippocampal SWRs became more frequent over several seconds after goal entry. The temporal segregation likely points to distinct goal-related processes. Immediately after goal entry, PFC and CA1 are separately engaged with their own beta-modulated networks; this then transitions to strong coordination between the two regions during hippocampal SWRs to support coordinated hippocampal-neocortical reactivation. Interestingly, on the timescale of milliseconds, ongoing PFC beta oscillations were reduced during hippocampal SWRs (Fig. 6 K), whereas CA1 beta power increased around the time of SWRs (Fig. 6 L). Across days of training, the timing relationship remained similar for PFC but weakened for CA1 (Fig. 6-2 B and D). This is again consistent with the hypothesis that PFC and CA1 beta are signatures for functionally distinct and uncoordinated processes between the two regions. When hippocampal SWRs occur, a switch in the hippocampal-neocortical state occurs, reducing ongoing neocortical beta oscillations in favor of reactivation-related coordination across hippocampal-neocortical networks (Jadhav et al., 2016; Yu et al., 2017). Despite our identification of goal period timing relationships between beta oscillations and hippocampal SWRs, further investigation is needed to determine the functional contribution of beta modulation during this period.

We found that beta modulation status identified a specific subpopulation of SWR-responsive PFC cells with distinct task firing correlates. Prior work shows that PFC firing during awake hippocampal SWRs can reactivate movement representations (Shin et al., 2019), where SWR-excited PFC cells prefer to spike during movement and SWR-inhibited PFC cells prefer to spike during immobility (Jadhav et al., 2016; Yu et al., 2017). Our results are consistent with these findings (Fig. 7 E). Interestingly, beta and SWR-modulated PFC cells had a distinguishing feature, where they better recapitulated the distance to goal relationship than non-beta-modulated PFC cells (Fig. 7 D). This suggests PFC cells with beta modulation may represent a cell subpopulation participating in the expression of goal-related states, beyond movement coding alone. These properties were specific to PFC since we did not find that beta modulation status was associated with distinct functional properties for SWR-modulated CA1 cells.

The difference in the frequency distribution of hippocampal and PFC beta raises the question of whether cortical and hippocampal beta have similar origins. Given that beta oscillations have been found in connected cortical, subcortical, and hippocampal regions, do they arise from similar mechanisms? Experimental evidence and computational models indicate that beta oscillations could be generated locally within a network (Bitzenhofer et al., 2017; David et al., 2015; Fourcaud-Trocmé et al., 2011; Osinski & Kay, 2016; Roopun et al., 2008; Roopun et al., 2006; Sherman et al., 2016; Womelsdorf et al., 2014). Beta was originally partitioned into a lower-frequency ∼15 Hz (beta1) band and a higher-frequency 20-30 Hz (beta2) band (Baker et al., 1997; MacKay & Mendonça, 1995; Roopun et al., 2008; Roopun et al., 2006). Our observations in CA1 are consistent with beta2 (20-30 Hz) (Berke et al., 2008; Igarashi et al., 2014; Iwasaki et al., 2021), with our observed beta bursts having a median peak frequency of approximately 22 Hz. The PFC beta we observed centered around 19 Hz with a narrow frequency distribution. This is closer to beta1 but is higher than the ∼15 Hz beta1 reported in various neocortical regions. Could the distinct beta-frequency distribution in these regions point to distinct origins? PFC beta may be generated locally or from a brain region with a lower-frequency beta generator. The wider distribution of CA1 beta burst peak frequencies suggests hippocampal beta may arise from multiple local or remote inputs from a variety of brain regions, each with a range of central frequencies above 20 Hz. These oscillations may be relayed to CA1, possibly via the midline thalamic regions, including the nucleus reuniens (Jayachandran et al., 2023), or via the entorhinal cortex (Kay & Freeman, 1998; Kay et al., 1996).

There is increasing evidence for the contribution of beta oscillations to a wide range of cognitive processes (Lundqvist et al., 2024; Miles et al., 2023), expanding from prior work focused on sensorimotor control (Barone & Rossiter, 2021; Engel & Fries, 2010) and sensory perception (Kay, 2014). While beta has been implicated in coordinating neural activity across brain networks, including the hippocampus and neocortex (Igarashi et al., 2014; Jayachandran et al., 2023; Spitzer & Haegens, 2017; Symanski et al., 2022), we identified an important period for learning during which the hippocampus and PFC could be modulated independently by distinct frequency bands within the beta range. Identifying functional subnetworks and when they coordinate can provide further insight into the temporal dynamics of distributed neural processes supporting cognition. An important future direction will be to test the causal contribution of beta-frequency dynamics across distributed brain networks in learning.

## Acknowledgements

We thank Leslie Kay for providing helpful feedback on the manuscript and Krish Khanna and Rajat Gupta for assisting with DeepLabCut.

## Author contributions

Experiment design and conceptualization (JYY, GW and NZ). Data collection (NZ, ZL, PT, AEK, JG and ARL). Data analysis (JYY and GW). Manuscript preparation (JYY) and editing (JYY, NZ, GW and ZL).

## Funding

Brain Research Foundation seed grant (JYY). Whitehall Foundation research grant (JYY). The University of Chicago Neuroscience Early-Stage Scientist Training Program (NINDS 5R25NS117360, AEK). The University of Chicago Post-Baccalaureate Research Education Program (NIGMS 3R25GM066522-18, JG). Leadership Alliance Summer Research Early Identification Program (ARL).

## Methods

### Animals

Long-Evans rats (Charles River Laboratories, 6 males, 4-10 months) were used for the study. All procedures were performed under approval by the university’s Institutional Animal Care and Use Committee, according to the guidelines of the Association for Assessment and Accreditation of Laboratory Animal Care.

### Neural implants

Animals were chronically implanted with neural recording devices (Voigts et al., 2020) with up to 32 adjustable tetrodes. Up to 16 tetrodes targeted the CA1 region of the left hippocampus (AP -3.8, ML -2.75 relative to bregma), and up to 16 tetrodes targeted both hemispheres of the prelimbic region of rat PFC (AP +3.5, ML ±1.5 relative to bregma). The PFC bundle was angled at 15° towards the midline to avoid the superior sagittal sinus. A ground screw was placed in the skull above the right cerebellum. Tetrodes were gradually adjusted into the target regions over two to three weeks after surgery, based on LFP markers and estimated depth of the tetrodes. Once the tetrodes were within the target brain regions, small adjustments of 25 μm, when necessary, were made to isolate single cells. Tetrode locations were verified at the end of the experiment. To mark the tip of the tetrode, electrolytic lesions were induced for each tetrode (100 μA, 2 s). Nissl staining was used to verify the location of the lesion was within the intended area.

### Spatial learning apparatus

All tracks and mazes were constructed from acrylic (black DP-9, 8 cm wide, 4 cm tall walls, 76 cm above floor level). Reward delivery in all tasks was automated. Infrared sensors were placed at the ends of the maze arms to detect the rat’s arrival and to trigger the reward. Reward was delivered by a syringe pump (100 μl at 20 ml/min, NE-500, New Era Pump Systems Inc, New York, USA).

Goal entry was defined as the time when the animal reached the reward delivery device and broke the infrared beam. All timestamps for beam breaks and for when the beam was restored at each goal were logged by the recording system.

### Behavioral training

Animals were food restricted to 85-90% of their baseline body weight. For pretraining, animals were first trained to forage for reward in a black open field box (H: 31 cm, W: 61 cm, L: 61 cm), for 10 minutes per day for 3 days. The reward was 70% evaporated milk (Carnation) with 5% sucrose. The reward was randomly dropped inside the open box to encourage foraging. The next pretraining phase involved the rats learning to run back and forth on an elevated linear track (60 cm) to consume reward from the ends. Animals were trained until they reached a criterion of 20 rewards per session (10-minute sessions, three times daily, 4 to 7 days).

For the tasks on the Y- and F-shaped mazes, the animals needed to learn a specific sequence of goal visits to be rewarded on every trial. There were three goal locations at the end of the arms in each maze. The animal needed to alternate visits between goals 2 and 3 via goal 1 (1, 2, 1, 3, 1, 2, 1, 3 …), similar to previously described spatial alternation tasks (Frank et al., 2000; Jadhav et al., 2012; Kim & Frank, 2009; Maharjan et al., 2018). For example, if the rat was at goal 1, it would be rewarded next at goal 2 only if it had previously visited goal 3. Similarly, if the rat was at goal 1, it would be rewarded next at goal 3 only if it had previously visited goal 2. If the rat was at goals 2 or 3, it would be rewarded only by going to goal 1. Reward delivery was controlled using custom scripts written for the data recording system.

### Data recording

Data were collected using the Trodes data acquisition system (SpikeGadgets LLC, California, USA). Neural data were recorded using a 128-channel digitizing headstage. The headstage was tethered via a commutator to the main data acquisition system. The commutator and tether were supported by a passive suspension system. Raw data were recorded at 30 kHz. Local field potential (<300 Hz) was extracted at 1500 Hz. Time-synchronized video (1456 x 1088, 30 Hz) was recorded using Allied Vision Manta G-158C cameras.

### Behavioral analysis

DeepLabCut (Mathis et al., 2018) was used to track the animal in the video. The spatial coordinate for the neck of the animal was used as the animal’s location and to calculate the animal’s speed. Performance of the animals was derived using a state-space model that estimated the probability of the animal making a correct choice (Smith et al., 2004). We then calculated the mean performance for each day, and also grouped performance by quintiles.

### Data and statistical analysis

We used Python packages for signal processing: Elephant electrophysiology toolkit (elephant.readthedocs.io), fooof tools (fooof-tools.github.io), GhostiPy (github.com/kemerelab/ghostipy); circular statistics: PyCircStat (github.com/circstat/pycircstat); and statistics: SciPy (scipy.org), statsmodels (statsmodels.org) and pingouin (pingouin-stats.org).

We used linear mixed-effects models (statsmodels mixedlm) to perform regression analysis to account for the contribution of fixed effects from the independent variable and the random effects from individual variation between animals. We report the marginal R^2^ (R^2^_m_), which is the proportion of variance explained by the fixed effects and the conditional R^2^ (R^2^_c_), which is the proportion of variance explained by the entire model (fixed and random effects) (Nakagawa & Schielzeth, 2013).

### Spectral analysis

We calculated the power spectral density (PSD) using the Welch function, with 1 s segments with 50% overlap (scipy.signal.welch). We calculated coherence using the coherence function with 1 s segments and 50% overlap (scipy.signal.coherence). We isolated the periodic component of the PSD by subtracting the aperiodic component using Fitting Oscillations & One-Over-F (fooof-tools) (Donoghue et al., 2020). This was done for each behavior session, and the mean was calculated across all sessions per day or for sessions on each maze separately. For periods at the goal, we only included 1 s segments when the animal was immobile (speed < 0.5 cm/s). For periods on the maze, we only included 1 s segments when the animal was mobile (speed ≥ 4 cm/s). We calculated the difference in PSD between goal and maze periods by subtracting the periodic component PSDs. A 95% confidence interval for the difference was generated using a permutation test (1000 shuffles) where we permuted the maze or goal identity of each PSD.

We repeated the spectral analysis using continuous wavelet transform (CWT) for a ±3 s window centered on goal entry, using a generalized Morse wavelet (γ = 4, β = 7.5) for 2-100 Hz in 1 Hz steps (GhostiPy cwt). We calculated the mean PSD for the 3 s window on the maze before goal entry and the 3 s after goal entry, as well as the difference between the two. This was done for each behavior session, and the mean was calculated across all sessions per day or for sessions on each maze separately. A 95% confidence interval for the difference was generated using a permutation test (1000 shuffles) where we permuted the behavioral state (maze or goal) of each spectrogram.

### Beta burst detection

Local field potential (LFP) analysis was performed using the Elephant electrophysiology toolkit (Denker et al., 2024). We applied CWT to extract time-resolved spectral power for 0.5-130 Hz in 1 Hz steps. This was done using the GhostiPy package (Chu & Kemere, 2021). The average power in the beta band (15-30 Hz) isolated from the CWT spectrogram was used to detect bursts using a dual-threshold amplitude method (Feingold et al., 2015). Bursts were defined as periods when the beta power envelope exceeded 1 standard deviation (SD) above the mean power with a minimum duration of 100 ms. The onset and offset of each burst were marked by the first upward and downward crossings of 0.5 SD above the mean, respectively. Burst peaks were characterized by the z-scored amplitude at the power maximum. Events separated by less than 100 ms were combined. Periods where the maximum LFP signal amplitude across all tetrodes exceeded 10 standard deviations were classified as noise and excluded from analyses.

We identified the time at which the burst had its peak power, the magnitude of the peak power, and the frequency at the time of peak power. This was done by isolating the CWT spectrogram for each burst and extracting the time and frequency corresponding to the maximum power value within the burst. Bursts were classified as independent or coincident based on overlap. Independent bursts are those without overlap in time, whereas coincident bursts overlap in time.

### Burst-aligned power and coherence analysis

Event-aligned power and coherence within and across PFC and CA1 (Fig. 2-2) were computed using the multi-taper method (2 tapers with a time-bandwidth product of 2) (Python implementation of the coherence function from chronux.org) (Mitra & Bokil, 2008). We calculated the coherence in a 1 s window centered at the peak of each burst (500 ms moving window, 5 ms steps). For within-region coherence, the coherence value was the mean for all pairs of tetrodes. To determine the change in coherence around the peak of the burst, a baseline was defined as the mean coherence in the −500 to −375 ms interval relative to the center of the burst.

### Sharp-wave ripple detection

Sharp-wave ripples (SWRs) were identified based on previous methods (github.com/Eden-Kramer-Lab/ripple_detection) (Kay et al., 2016). LFP from tetrodes in the dorsal CA1 region (oriens and pyramidale) was filtered in the ripple frequency band (150-250 Hz). The median of the smoothed envelope (Gaussian kernel σ = 4 ms) was calculated. We used a threshold of 2 standard deviations for detecting SWRs, with a minimum duration of 50 ms and separation of 50 ms. Only SWR events occurring at speed <4 cm/s and at the goal locations were included. We filtered out detection artifacts due to LFP noise by excluding time intervals in which the LFP signal magnitude exceeded 10 standard deviations across any tetrode.

### Spike sorting

We used MountainSort (Chung et al., 2017) with manual curation to isolate single cells. The quality of well-isolated cells was determined by having an interspike interval >2 ms, consistency of the waveforms, and the stability of the peak voltage values over the sessions.

### Cell-type classification

We classified each single unit as a pyramidal cell or an interneuron using established methods based on firing rate (Jadhav et al., 2016; Kay et al., 2016), trough-to-peak width (Petersen et al., 2021), and autocorrelation function mean (Kay et al., 2016). The thresholds for classification are shown in Table 5-1. Cells were only included if they had at least 20 spikes in the 5 s post-goal period across all trials in a session, with a minimum of 5 trials.

### Spike-aligned power spectrum

For each cell, the spike-aligned power spectrum (Fig. 5 C-D) was generated by extracting the CWT at the time of each spike and taking the average.

### Spike phase-locking analysis

For each tetrode, the phase of beta oscillations was extracted by first applying a band-pass filter to the raw LFP signal within the beta-band range (15-30 Hz, acausal 4^th^-order Butterworth), followed by a Hilbert transform using the Elephant toolkit (Denker et al., 2024). For each cell, we calculated the phase at each spike relative to the beta phase for every available tetrode, which allowed us to determine the cell’s phase-locking preference to beta within the same brain region (local) and the other region (remote). We used two metrics to quantify the strength of spike phase locking to beta. First, we performed the Rayleigh test for circular non-uniformity and calculated the Rayleigh-z value using PyCircStat (Berens, 2009). Second, we fitted a von Mises distribution to the phase preference histogram to estimate the mean phase and the concentration parameter (kappa) (scipy.stats.vonmises). To determine the local phase-locking strength of a given cell, we chose the maximum values of the phase-locking metric across all tetrodes within the same region as the cell. Similarly, we calculated the remote phase-locking strength as the maximum phase-locking value across all tetrodes in the other brain region. Only spikes that occurred when the animal was at the goal location were included for the phase-locking analysis.

### Task firing peri-event histogram and task modulation

To characterize the firing relative to goal entry, spike times of each unit were aligned to the time of goal entry for goal visits longer than 5 s. For this analysis, we only included units that spiked on more than 10 trials and had at least 50 spikes within ± 5 s of goal entry. Peri-event time histograms (PETHs) were constructed by binning spike times (± 5 s window centered on goal entry, 50 ms bins), dividing by the trial count and the bin width to obtain firing rate (Hz), and smoothing with a Gaussian kernel (σ = 150 ms). To test whether a unit’s firing was modulated relative to goal entry, we used a circular-shift permutation test. On each of 1000 iterations, the spike times of every trial were circularly shifted by a random offset, preserving the spike count and interspike interval structure of each trial while destroying the relation between spikes and goal entry, and the PETH was recomputed. The observed PETH was z-scored against the mean and standard deviation of the shuffled PETHs, and the test statistic was the maximum absolute z within ± 4.5 s (excluding 0.5 s at each edge to avoid smoothing artifacts). The p-value was the proportion of shuffles whose maximum absolute z was greater than or equal to the observed value. Units with p < 0.05 were classified as goal-entry-modulated.

### Reward outcome decoding

We decoded rewarded versus unrewarded goal visits from the population spike count in a 1 s window after goal entry. Sessions with fewer than 3 neurons or fewer than 3 visits of either outcome were excluded. We used logistic regression (L2-regularized, balanced class weights, features z-scored within training folds) and evaluated by ROC AUC under repeated stratified k-fold cross-validation (3 folds × 10 repeats). Significance was assessed per session by rerunning the same procedure on 1000 outcome-label shuffles (p = fraction of shuffled AUCs ≥ observed; significant at p < 0.05).

### Task correlates for spiking activity

To determine if each cell’s beta-phase preference could distinguish between goal locations, we used a Gaussian Naïve Bayes model (scikit-learn, GaussianNB). For the model, the input feature was the mean spike beta-phase preference (5 s after goal entry), and the label was the goal identity. We determined the statistical significance of the classification accuracy using a permutation test (scikit-learn, permutation_test_score, 1000 shuffles and 3-fold cross-validation) where the goal identity labels were permuted. We used the same method to determine whether a cell’s mean firing rate could distinguish between the goal locations.

The mean spiking speed is the mean speed for all spikes for each cell. The distance to goal is the mean of the shortest linear distance from the location at each spike to a goal, accounting for the path geometry of the maze. The spatial firing entropy is the Shannon entropy in bits of a cell’s distance-normalized firing pattern on all trajectories. Higher entropy values indicate greater uniformity in spatial firing patterns across trajectories, whereas lower entropy values indicate more spatially selective firing across trajectories.

### SWR-aligned spectrogram

We applied CWT to extract time-resolved spectral power for 0.5-300 Hz in 1 Hz steps using the same parameters as other CWT analyses described earlier. We then aligned the PFC or CA1 spectrogram relative to the start of CA1 SWRs for each behavior session, and calculated a session mean. The mean spectrogram for all sessions for each region is shown in Fig. 6. We repeated this for all sessions on each maze in Fig. 6-3. From these spectrograms, we quantified the mean power in the SWR band (150-250 Hz) or beta band (15-30 Hz).

### SWR modulation of spiking

SWR modulation was determined based on previously described methods (Jadhav et al., 2016; Rothschild et al., 2017; Yu et al., 2017; Yu et al., 2018). For each cell, an SWR event-aligned spiking histogram was created for a 1 s window centered on the start of each SWR. Only SWRs with duration greater than 50 ms were included. A modulation index was calculated based on the difference in spiking rate within a 200 ms window after the start of the SWR relative to the baseline within the 1 s window. A circular-shift permutation test was used to determine the significance of the modulation index. A positive modulation index corresponds to an increase in firing around the time of SWRs, whereas a negative modulation index corresponds to a decrease in firing.

### Phase amplitude coupling

For each session, one tetrode from the pyramidal layer of CA1 and one tetrode from PFC were selected for the analysis. We extracted the power in the SWR frequency band for the time intervals during SWRs, from the CA1 tetrode. Next, we extracted the corresponding beta phase from the CA1 or the PFC tetrode for those intervals. We binned the phase values (-π to π, π/8 bins) and assigned the SWR band power to their respective phase bins, and calculated the average for each bin.

We then converted the average to a z-score across bins before averaging across all sessions on each maze. To calculate the chance power distribution, we used a permutation test with 5000 shuffles. For each shuffle, we circularly permuted the session z-score average for each session and recalculated the mean across all sessions. The 99% confidence interval for each phase bin was derived from the permuted means.

### Correlation between beta-phase-locking preference and SWR-modulation index

We calculated the Pearson’s correlation between a cell’s beta-phase preference and its SWR-modulation index. To estimate the significance of the correlation coefficient, we performed a permutation test. Each cell was classified based on its phase-locking significance to local beta (β+ or β-) within the first 5 s after goal entry, for goal visits lasting at least 5 s, and its SWR-modulation significance (SWR+ or SWR-). Only cells with more than 20 spikes across at least 5 goal visits were included for this analysis. This produces four possible combinations in the population (SWR^-^β^-^, SWR^-^β^+^, SWR^+^β^-^, or SWR^+^β^+^). We then permuted the β and SWR significance labels for the population and repeated the calculation. This was done 5000 times to generate a permuted correlation coefficient distribution, which allowed us to estimate the significance to p<0.0002. The p value of the observed correlation coefficient was calculated as the proportion of the permuted distribution that is greater than or less than the observed value.

## Figures

**Figure 1-1.**
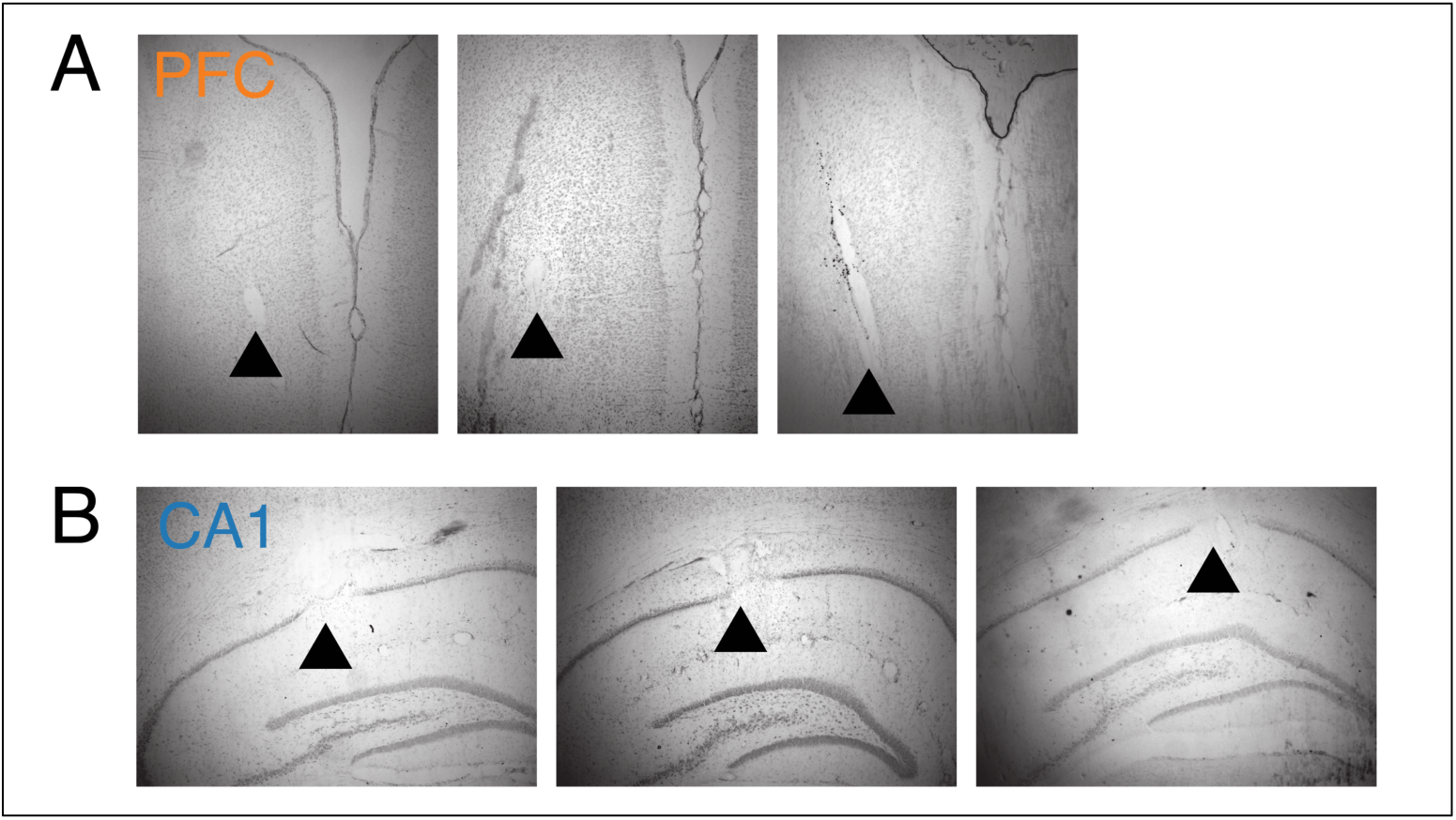
Location of recording tetrodes. **A.** Representative histology showing location of tetrodes in PFC. **B.** Representative histology showing location of tetrodes in CA1.

**Figure 1-2.**
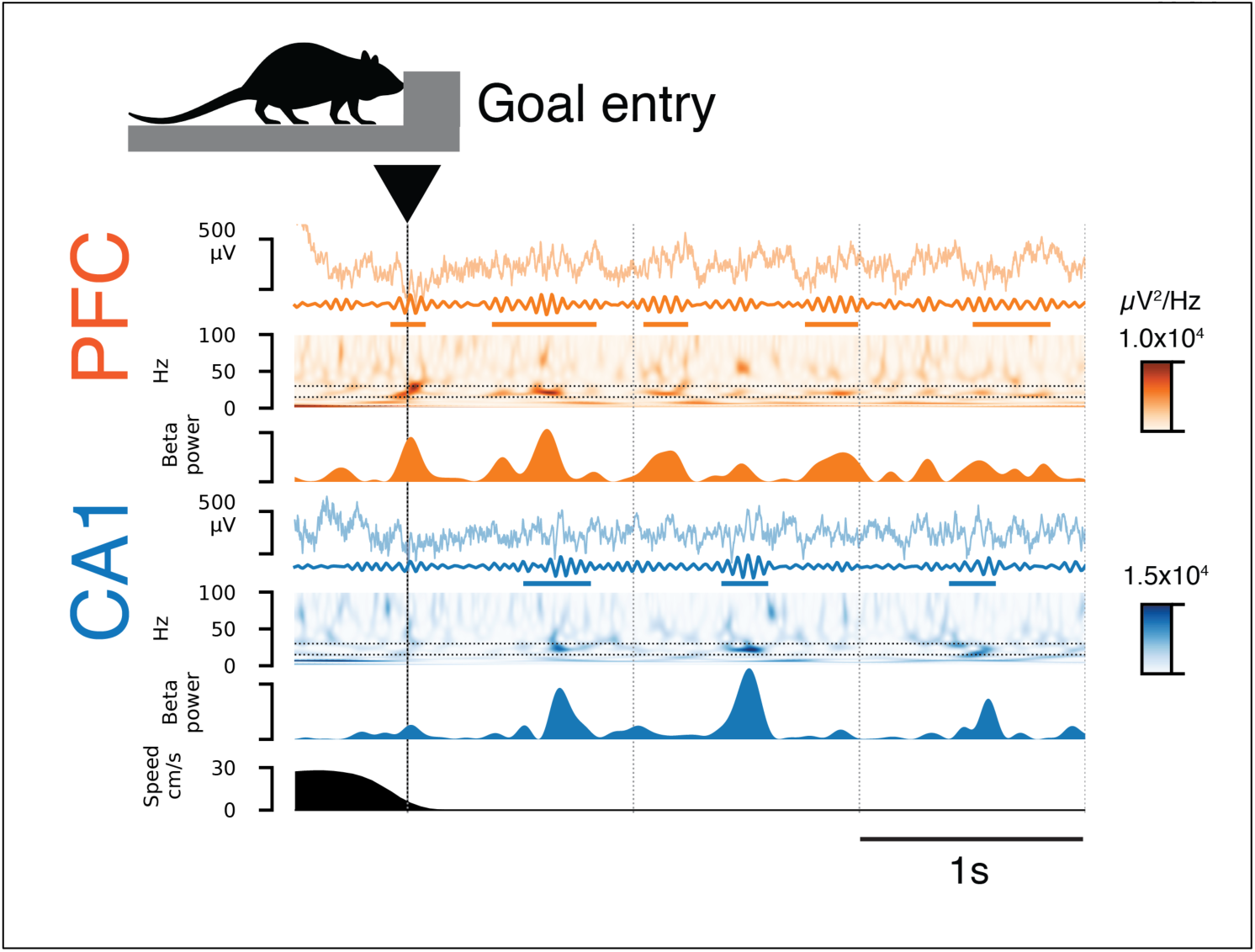
Extended frequency spectrogram for example goal-aligned PFC and CA1 LFP. Same example as Fig. 1 B with spectrogram up to 100 Hz.

**Figure 1-3.**
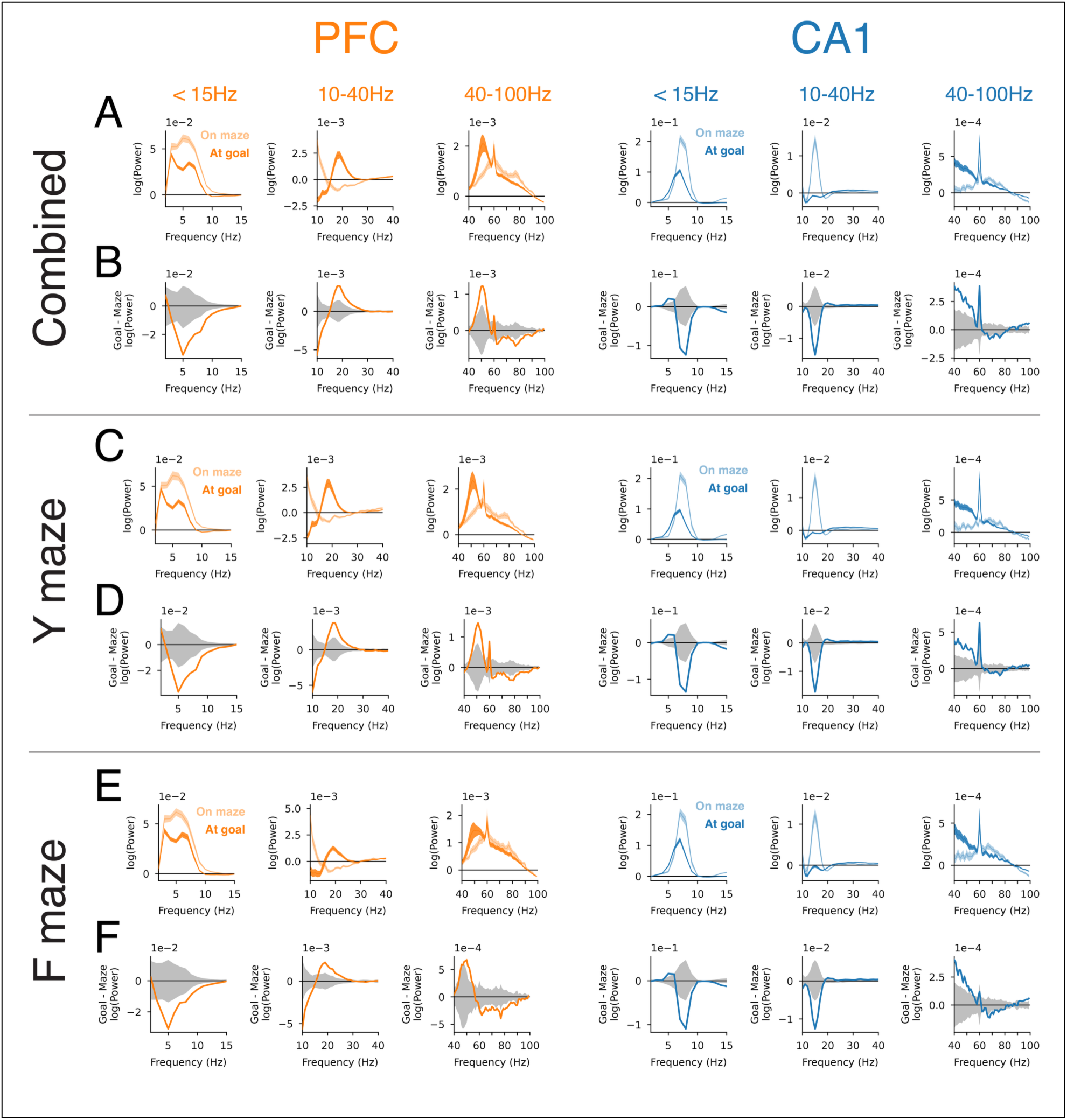
Power spectral density by brain region, frequency band and maze task. **A.** Average across both Y and F mazes. Periodic PSD component for PFC (orange) and CA1 (blue) for periods when the animal is running on the maze or immobile at the goal. Each column corresponds to a specific frequency range (marked above each plot) to display the appropriate power range. The sharp peak at 60 Hz corresponds to line noise. **B.** Average across both Y and F mazes. Difference between goal and maze periodic PSDs with 95% confidence interval shown in gray. **C.** Y maze only. Periodic PSD component for PFC (orange) and CA1 (blue). Same format as A. **D.** Y maze only. Difference between goal and maze periodic PSDs with 95% confidence interval shown in gray. **E.** F maze only. Periodic PSD component for PFC (orange) and CA1 (blue). Same format as A. **F.** F maze only. Difference between goal and maze periodic PSDs with 95% confidence interval shown in gray.

**Figure 1-4.**
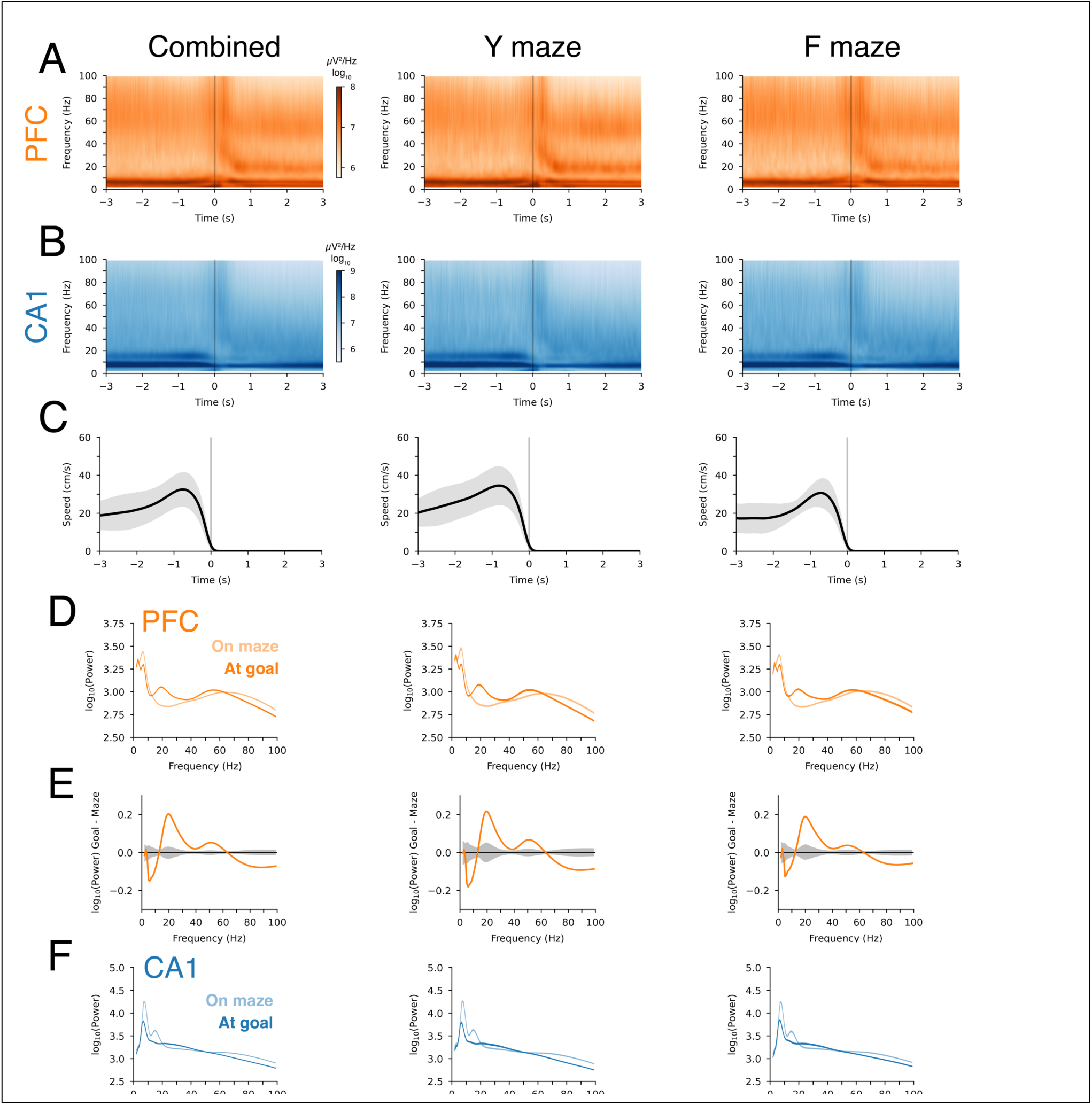
Goal-entry-aligned spectrogram. **A.** PFC spectrogram for all maze task sessions (left), Y maze (center) or F maze (right). Spectrogram generated using continuous wavelet transform. Session count shown in Table 1-1. **B.** CA1 spectrogram. **C.** Corresponding goal-entry-aligned speed profile (mean±std). **D.** Mean PFC power for 3 s before (maze) or after goal entry (goal). **E.** Difference in PFC power for 3 s before (maze) or after goal entry (goal). 95% confidence interval shown in gray. **F.** Mean CA1 power for 3 s before (maze) or after goal entry (goal). **G.** Difference in CA1 power for 3 s before (maze) or after goal entry (goal). 95% confidence interval shown in gray.

**Figure 1-5.**
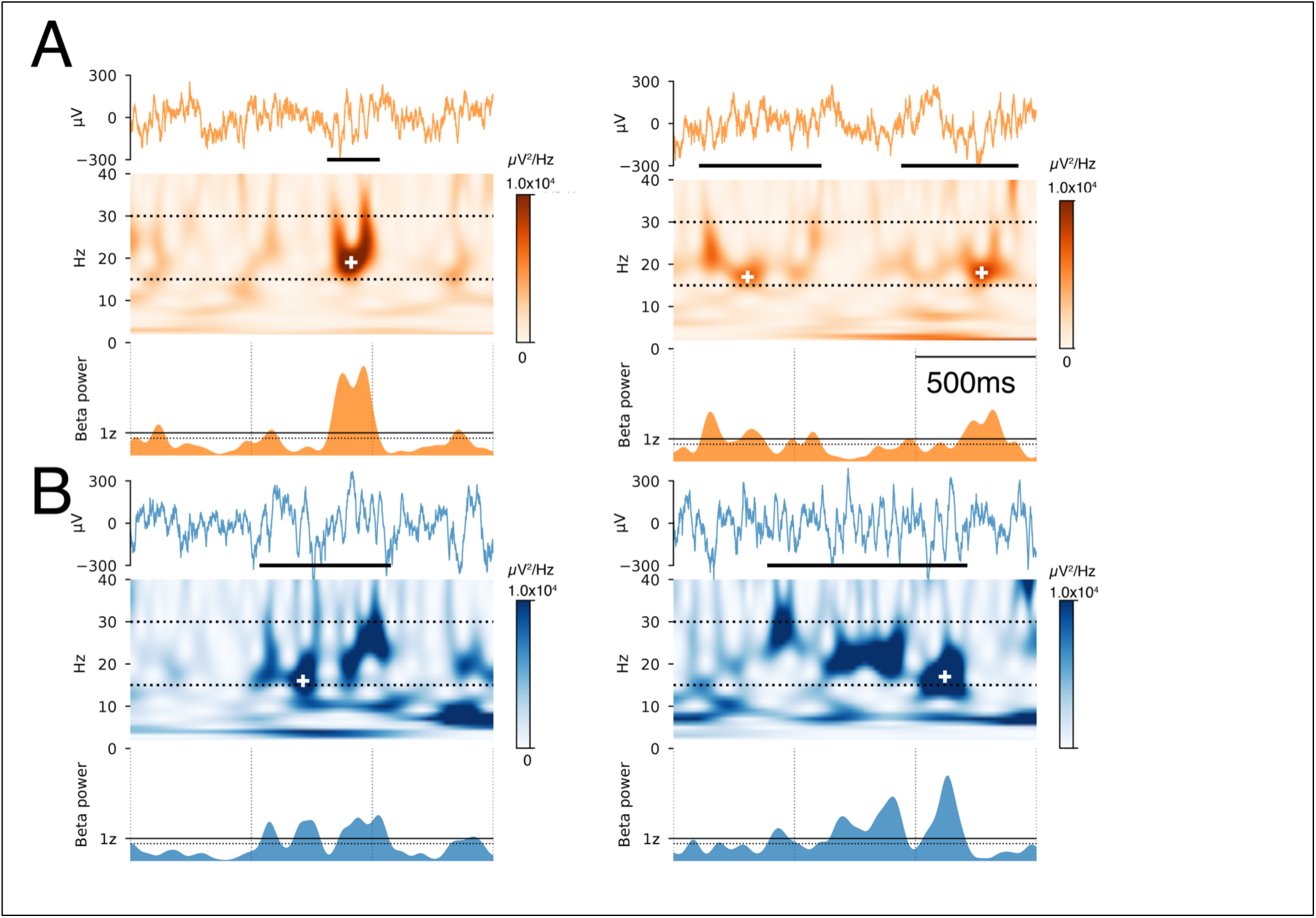
Burst detection. **A.** Two example bursts with corresponding LFP and continuous wavelet transform from PFC (1.5 s each, top row). The burst is marked by the black horizontal line. The corresponding continuous wavelet transform is shown in the middle row. The white cross marks the frequency and time corresponding to the maximum power in the burst interval. The mean power in the beta band (15-30 Hz) from the continuous wavelet transform is shown by the histogram in the bottom row. The solid black line corresponds to 1 standard deviation (z) above the mean (1z), and the dotted black line corresponds to 0.5 standard deviation above the mean (0.5z). Bursts were defined as periods when the beta power envelope exceeded 1z above the mean power with a minimum duration of 100 ms, with the start and end defined by crossings of the 0.5z threshold. Intervals separated by less than 100 ms are merged. The peak frequency for each burst is the frequency corresponding to the maximum power within the burst interval. **B.** Two example bursts from CA1.

**Figure 1-6.**
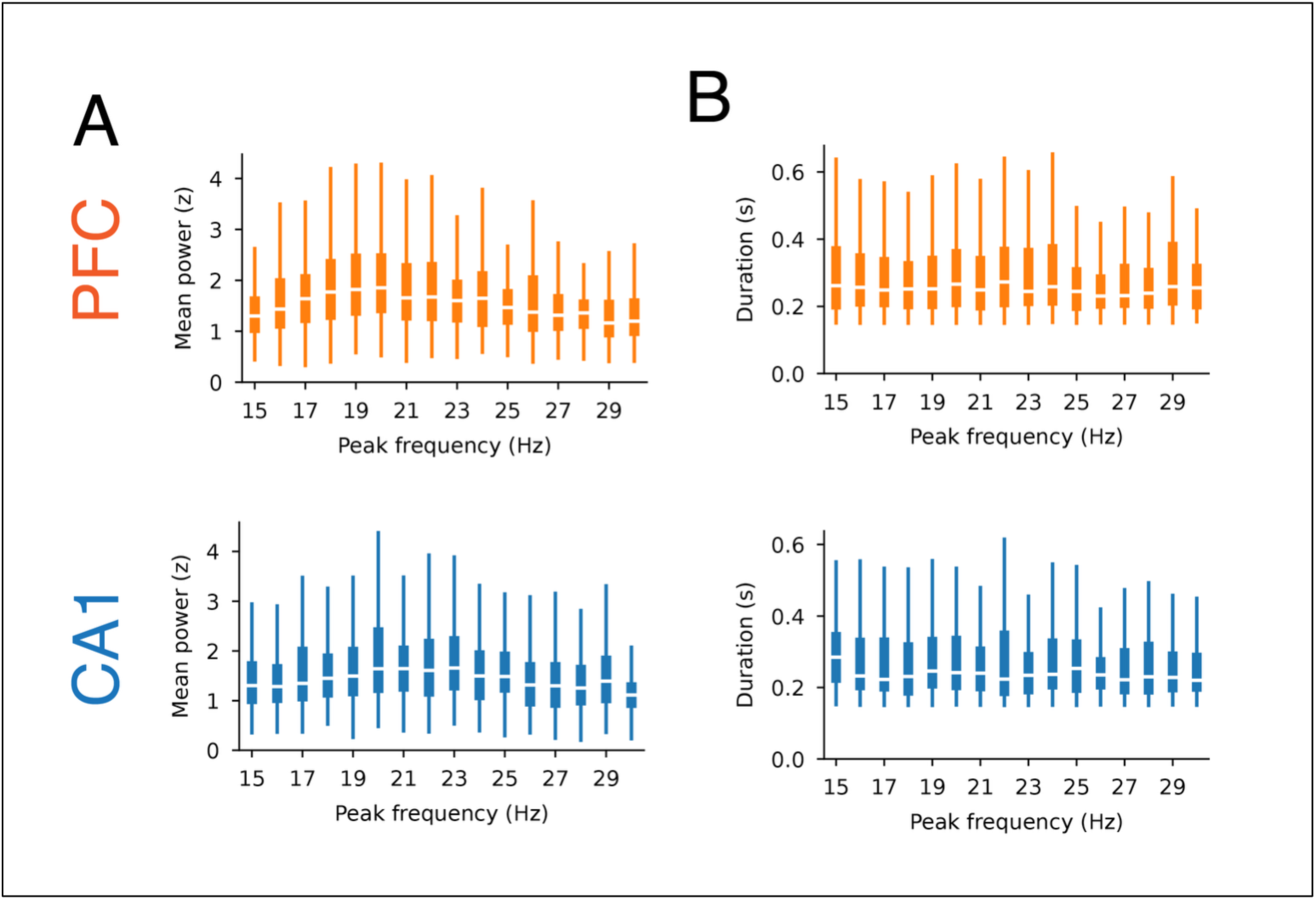
Power and duration for beta bursts by frequency. **A.** Beta burst power by peak burst frequency. **B.** Beta burst duration by peak burst frequency.

**Table 1-1.**
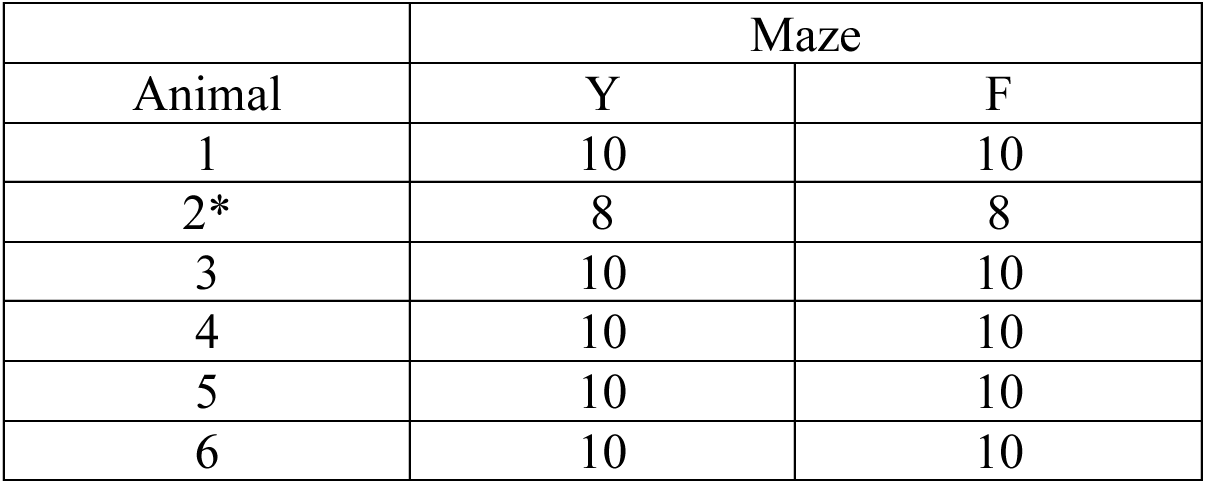
Session count. Count of sessions for each maze task. Day 5 for animal 2 was excluded due to noise in the recording.

|  | Maze |  |
| --- | --- | --- |
| Animal | Y | F |
| 1 | 10 | 10 |
| 2* | 8 | 8 |
| 3 | 10 | 10 |
| 4 | 10 | 10 |
| 5 | 10 | 10 |
| 6 | 10 | 10 |

**Table 1-2.** Tetrode count. Count of tetrodes in each brain region and hemisphere across days.

| Animal | Day | dCA1 | PFC |  |  |
| --- | --- | --- | --- | --- | --- |
|  |  | Left | Left | Right | Total |
| 1 | 1 | 14 | 6 | 6 | 12 |
|  | 2 | 14 | 6 | 6 | 12 |
|  | 3 | 11 | 6 | 6 | 12 |
|  | 4 | 13 | 6 | 6 | 12 |
|  | 5 | 11 | 6 | 6 | 12 |
| 2 | 1 | 13 | 0 | 7 | 7 |
|  | 2 | 12 | 0 | 7 | 7 |
|  | 3 | 13 | 5 | 6 | 11 |
|  | 4 | 14 | 5 | 7 | 12 |
| 3 | 1 | 3 | 0 | 6 | 6 |
|  | 2 | 5 | 0 | 6 | 6 |
|  | 3 | 4 | 0 | 6 | 6 |
|  | 4 | 4 | 0 | 6 | 6 |
|  | 5 | 4 | 0 | 6 | 6 |
| 4 | 1 | 12 | 7 | 7 | 14 |
|  | 2 | 11 | 7 | 7 | 14 |
|  | 3 | 12 | 7 | 7 | 14 |
|  | 4 | 6 | 7 | 7 | 14 |
|  | 5 | 4 | 7 | 7 | 14 |
| 5 | 1 | 1 | 6 | 6 | 12 |
|  | 2 | 2 | 6 | 6 | 12 |
|  | 3 | 1 | 5 | 4 | 9 |
|  | 4 | 1 | 6 | 3 | 9 |
|  | 5 | 1 | 6 | 2 | 8 |
| 6 | 1 | 2 | 5 | 8 | 13 |
|  | 2 | 8 | 5 | 7 | 12 |
|  | 3 | 10 | 5 | 7 | 12 |
|  | 4 | 10 | 6 | 8 | 14 |
|  | 5 | 9 | 6 | 6 | 12 |

**Figure 2-1.**
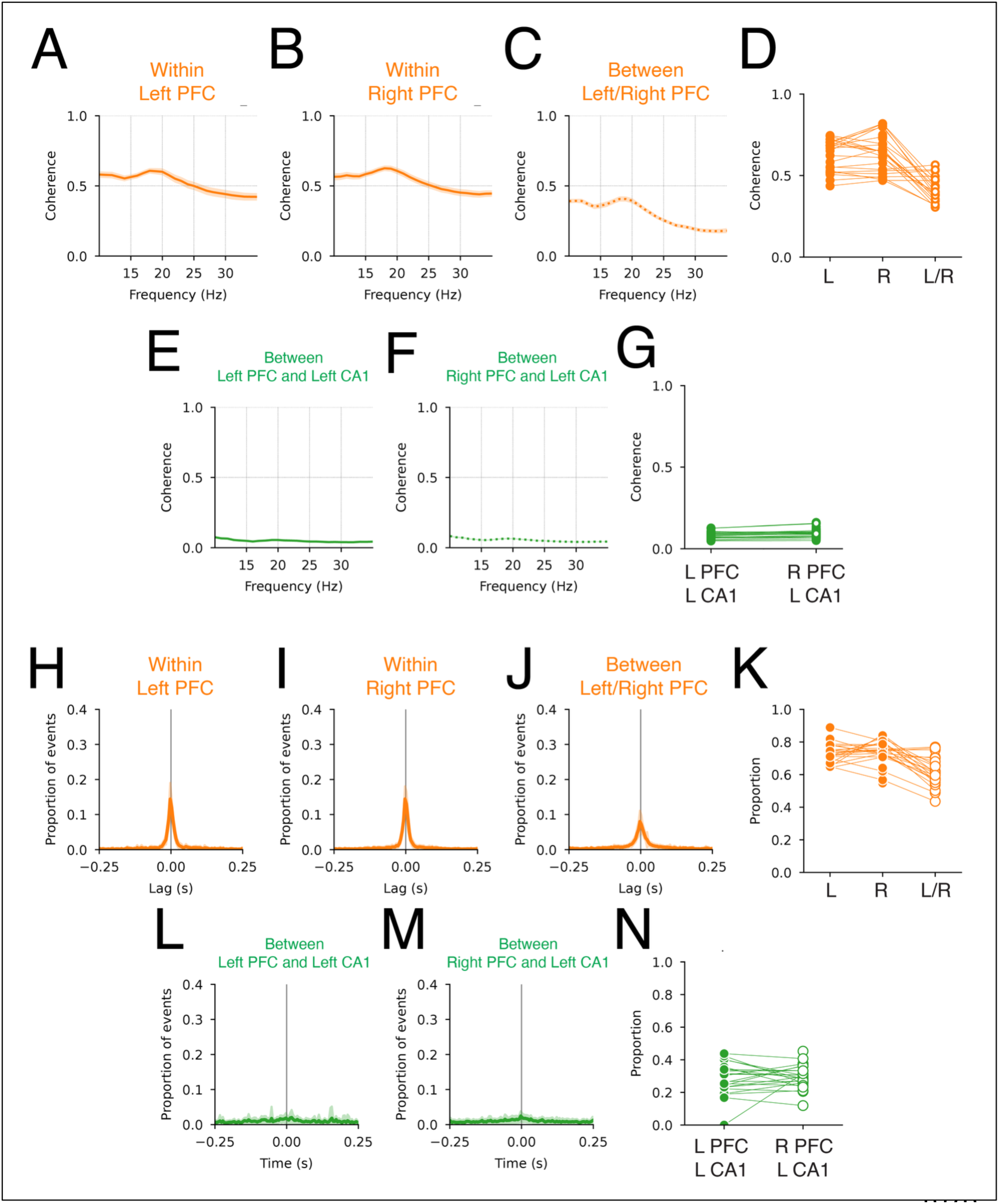
Coherence and burst coincidence within and across each hemisphere. **A.** Coherence within left PFC (mean±SEM). **B.** Coherence within right PFC (mean±SEM). **C.** Coherence between left and right PFC (mean±SEM). **D.** Beta-frequency coherence within each and across both PFC hemispheres. Each data point is the mean of each experiment day. **E.** Coherence between ipsilateral PFC and CA1. **F.** Coherence between contralateral PFC and CA1. **G.** Beta-frequency coherence for ipsi- and contralateral PFC and CA1. **H.** Cross-correlation (mean±SD) of burst peak times for bursts detected on tetrodes within left PFC. **I.** Cross-correlation (mean±SD) of burst peak times for bursts detected on tetrodes within right PFC. **J.** Cross-correlation (mean±SD) of burst peak times for bursts detected between left and right PFC. **K.** Proportion of the cross-correlation density within ±50 ms lag for left, right and across PFC hemisphere. **L. Cross**-correlation (mean±SD) of burst peak times for bursts detected between ipsilateral PFC and CA1. **M.** Cross-correlation (mean±SD) of burst peak times for bursts detected between contralateral PFC and CA1. **N.** Proportion of the cross-correlation density within ±50 ms lag for ipsi- and contralateral PFC and CA1.

**Figure 2-2.**
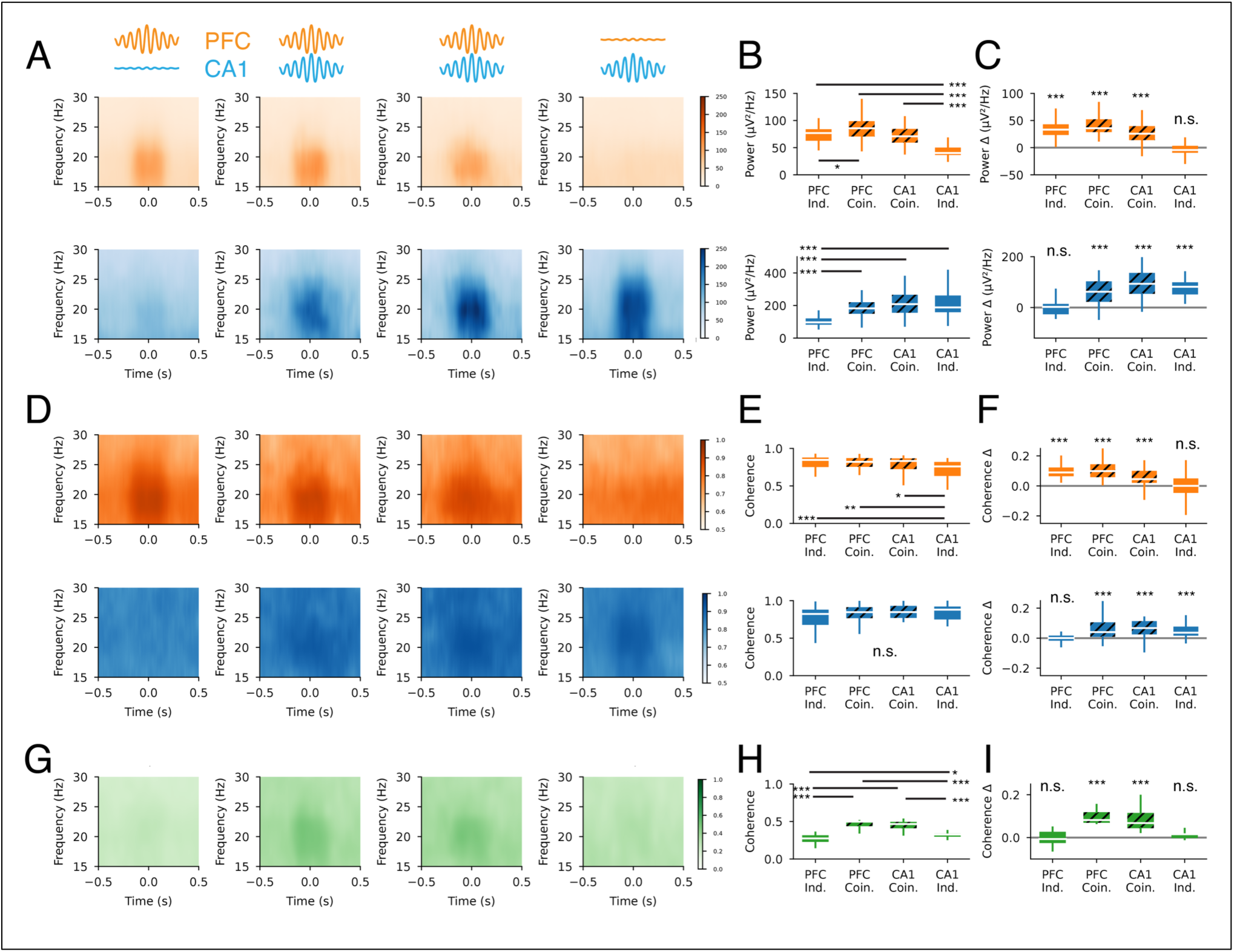
Beta burst-aligned power and coherence. **A.** Multi-taper power spectrum within a 1 s window centered on the peaks of independent or coincident beta bursts (columns) in the PFC (top row) and CA1 (bottom row). The heatmap shows the mean across all bursts in each category. **B.** Power (18-25 Hz) within ±100 ms of the peak. PFC (orange, top) and CA1 (blue, bottom). Pairwise rank-sum test with Benjamini-Hochberg false discovery rate correction: p<0.001*** and p<0.05* for all significant comparisons. **C.** Change in power within ±100 ms of the peak relative to baseline (125 ms from −0.5 s). PFC (orange, top) and CA1 (blue, bottom). Wilcoxon signed-rank test with Benjamini-Hochberg false discovery rate correction: p<0.001***. **D.** Multi-taper coherence in PFC (top row) and CA1 (bottom row) within a 1 s window centered on the peaks of independent or coincident beta bursts. The heatmap shows the mean across all bursts in each category. **E.** Coherence (18-25 Hz) within ±100 ms of the peak in PFC (top row) and CA1 (bottom row). Pairwise rank-sum test with Benjamini-Hochberg false discovery rate correction: p<0.001*** and p<0.05* for all significant comparisons. **F.** Change in coherence within ±100 ms of the peak relative to baseline (125 ms from -0.5 s). PFC (orange, top) and CA1 (blue, bottom). Wilcoxon signed-rank test with Benjamini-Hochberg false discovery rate correction: p<0.001***. **G.** PFC-CA1 coherence within a 1 s window centered on the peaks of independent or coincident beta bursts. The heatmap shows the mean across all bursts in each category. **H.** PFC-CA1 coherence (18-25 Hz) within ±100 ms of the peak in PFC (top row) and CA1 (bottom row). Pairwise rank-sum test with Benjamini-Hochberg false discovery rate correction: p<0.001*** and p<0.01** for all significant comparisons. **I.** Change in PFC-CA1 coherence within ±100 ms of the peak relative to baseline (125 ms from −0.5 s). PFC (orange, top) and CA1 (blue, bottom). Wilcoxon signed-rank test with Benjamini-Hochberg false discovery rate correction: p<0.001*** and p<0.01**.

**Table 2-1.** Session summary. Session and trial counts for each day for each animal.

| Animal | Day | Sessions | Trials |
| --- | --- | --- | --- |
| 1 | 1 | 4 | 56 |
|  | 2 | 4 | 44 |
|  | 3 | 4 | 45 |
|  | 4 | 4 | 41 |
|  | 5 | 4 | 73 |
| 2 | 1 | 4 | 46 |
|  | 2 | 4 | 59 |
|  | 3 | 4 | 79 |
|  | 4 | 4 | 117 |
| 3 | 1 | 4 | 96 |
|  | 2 | 4 | 96 |
|  | 3 | 4 | 119 |
|  | 4 | 4 | 104 |
|  | 5 | 4 | 150 |
| 4 | 1 | 4 | 88 |
|  | 2 | 4 | 63 |
|  | 3 | 4 | 65 |
|  | 4 | 4 | 77 |
|  | 5 | 4 | 84 |
| 5 | 1 | 4 | 66 |
|  | 2 | 4 | 68 |
|  | 3 | 4 | 92 |
|  | 4 | 4 | 86 |
|  | 5 | 4 | 106 |
| 6 | 1 | 4 | 60 |
|  | 2 | 4 | 74 |
|  | 3 | 4 | 102 |
|  | 4 | 4 | 92 |
|  | 5 | 4 | 109 |
| Total |  | 116 | 2357 |

**Figure 3-1.**
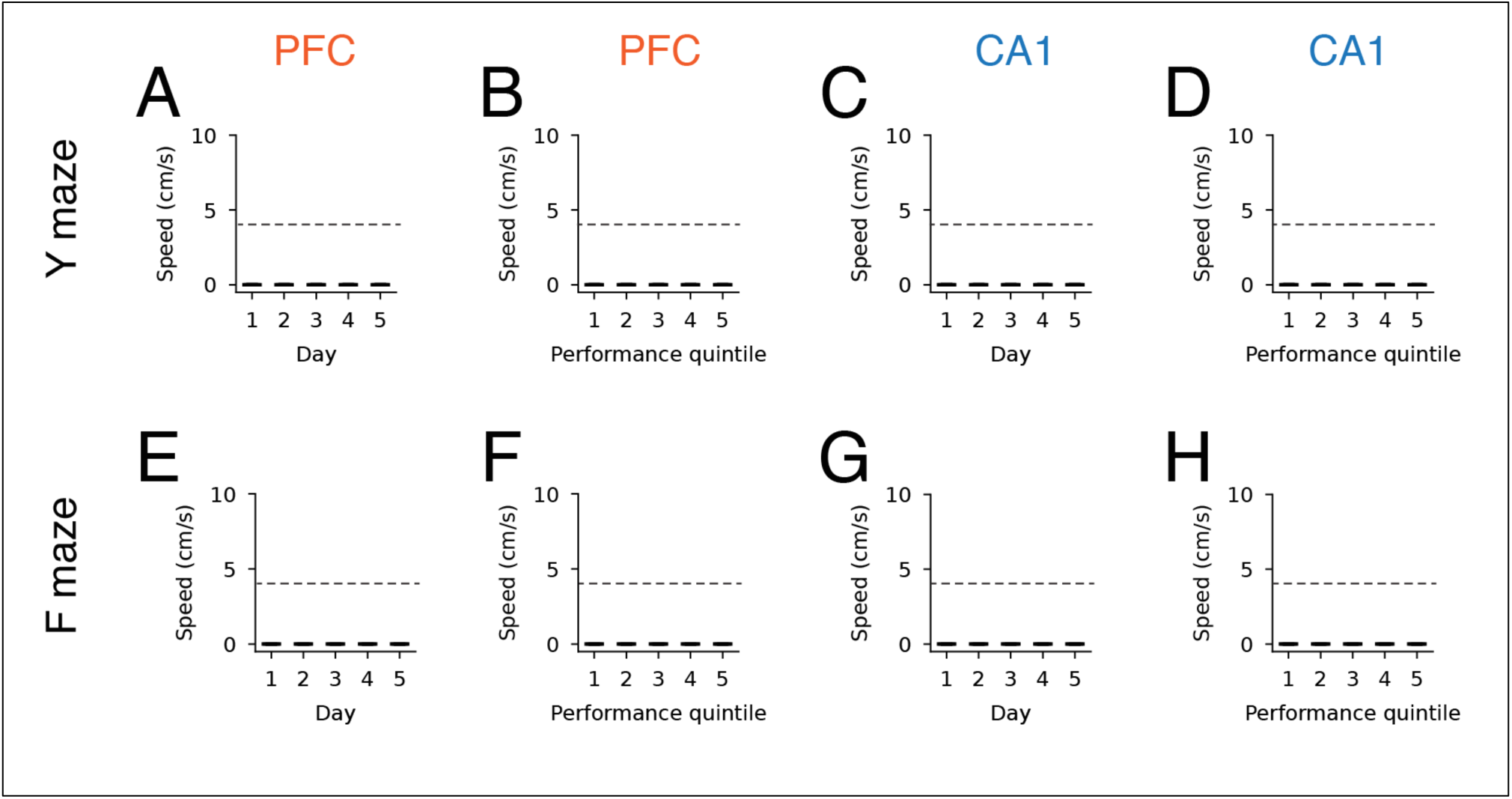
Animals are immobile at the time of beta bursts. **A.** Animal speed during PFC bursts in Y maze task by day. Dotted line shows the threshold for immobility (4 cm/s). **B.** Animal speed during PFC bursts in Y maze task by performance. **C.** Animal speed during CA1 bursts in Y maze task by day. **D.** Animal speed during CA1 bursts in Y maze task by performance. **E.** Animal speed during PFC bursts in F maze task by day. **F.** Animal speed during PFC bursts in F maze task by performance. **G.** Animal speed during CA1 bursts in F maze task by day. **H.** Animal speed during CA1 bursts in F maze task by performance.

**Figure 3-2.**
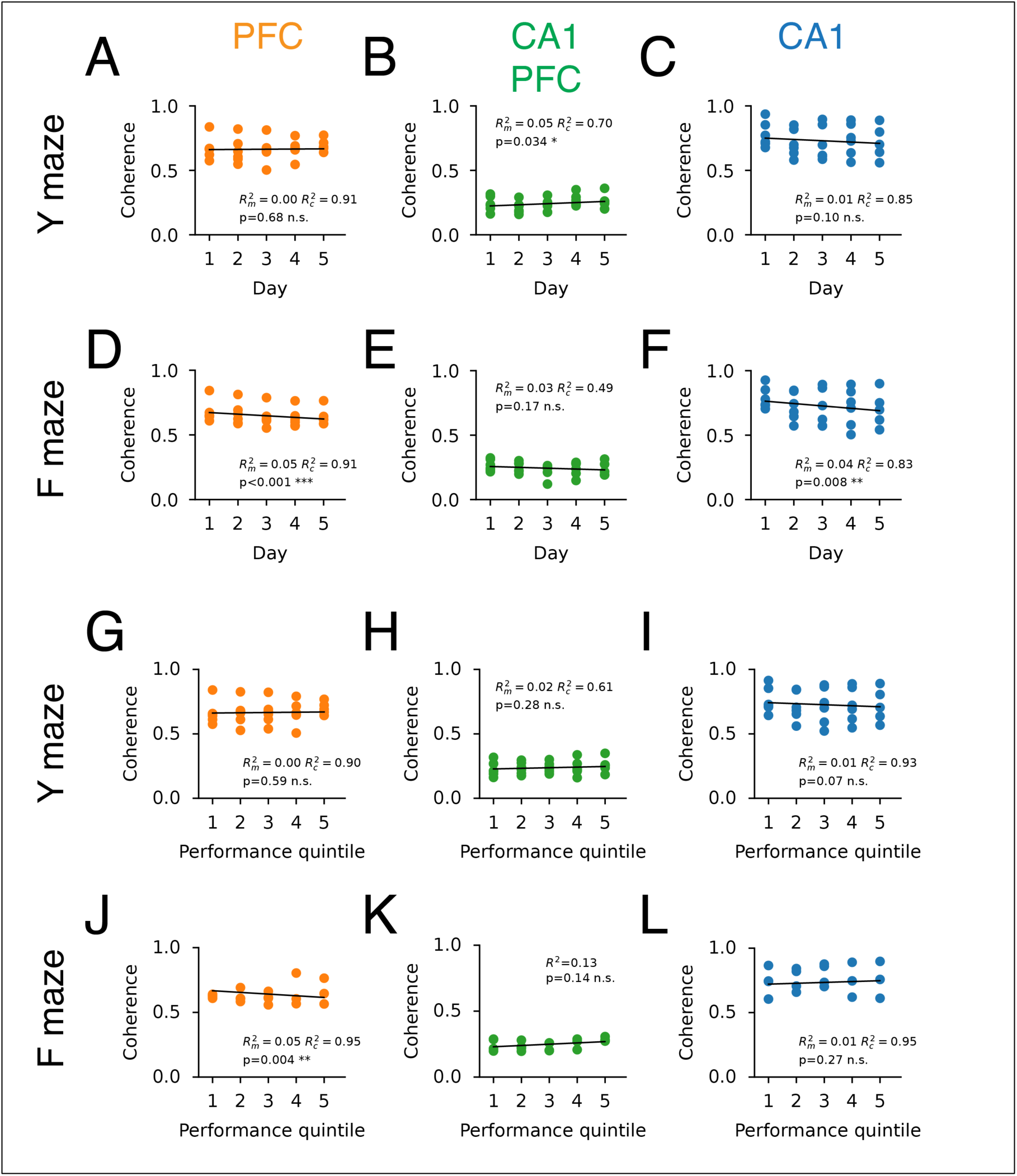
Beta coherence within and across brain regions by experiment day or performance, for each maze. **A.** PFC coherence by day in the Y maze task. **B.** PFC-CA1 coherence by day in the Y maze task. **C.** CA1 coherence by day in the Y maze task. **D.** PFC coherence by day in the F maze task. **E.** PFC-CA1 coherence by day in the F maze task. **F.** CA1 coherence by day in the F maze task. **G.** PFC coherence by performance in the Y maze task. **H.** PFC-CA1 coherence by performance in the Y maze task. **I.** CA1 coherence by performance in the Y maze task. **J.** PFC coherence by performance in the F maze task. **K.** PFC-CA1 coherence by performance in the F maze task. **L.** CA1 coherence by performance in the F maze task.

**Figure 3-3.**
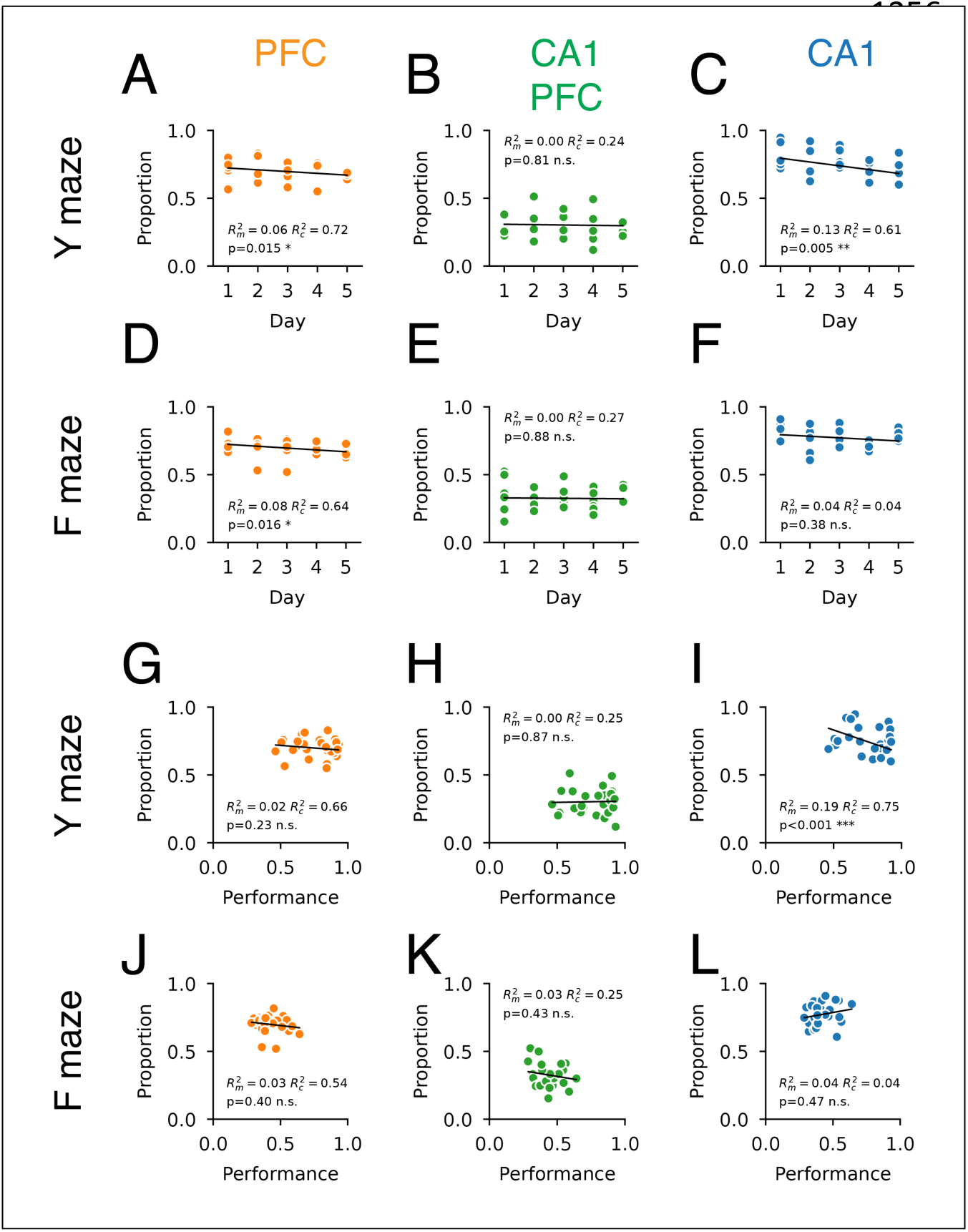
Beta burst coincidence within and across brain regions by experiment day or performance for each maze. **A.** PFC burst coincidence by day in the Y maze task. Value shows the proportion of the cross-correlation density within ±50 ms lag, similar to Fig. 2 E. **B.** PFC-CA1 burst coincidence by day in the Y maze task. **C.** CA1 burst coincidence by day in the Y maze task. **D.** PFC burst coincidence by day in the F maze task. **E.** PFC-CA1 burst coincidence by day in the F maze task. **F.** CA1 burst coincidence by day in the F maze task. **G.** PFC burst coincidence by performance in the Y maze task. **H.** PFC-CA1 burst coincidence by performance in the Y maze task. **I.** CA1 burst coincidence by performance in the Y maze task. **J.** PFC burst coincidence by performance in the F maze task. **K.** PFC-CA1 burst coincidence by performance in the F maze task. **L.** CA1 burst coincidence by performance in the F maze task.

**Figure 4-1.**
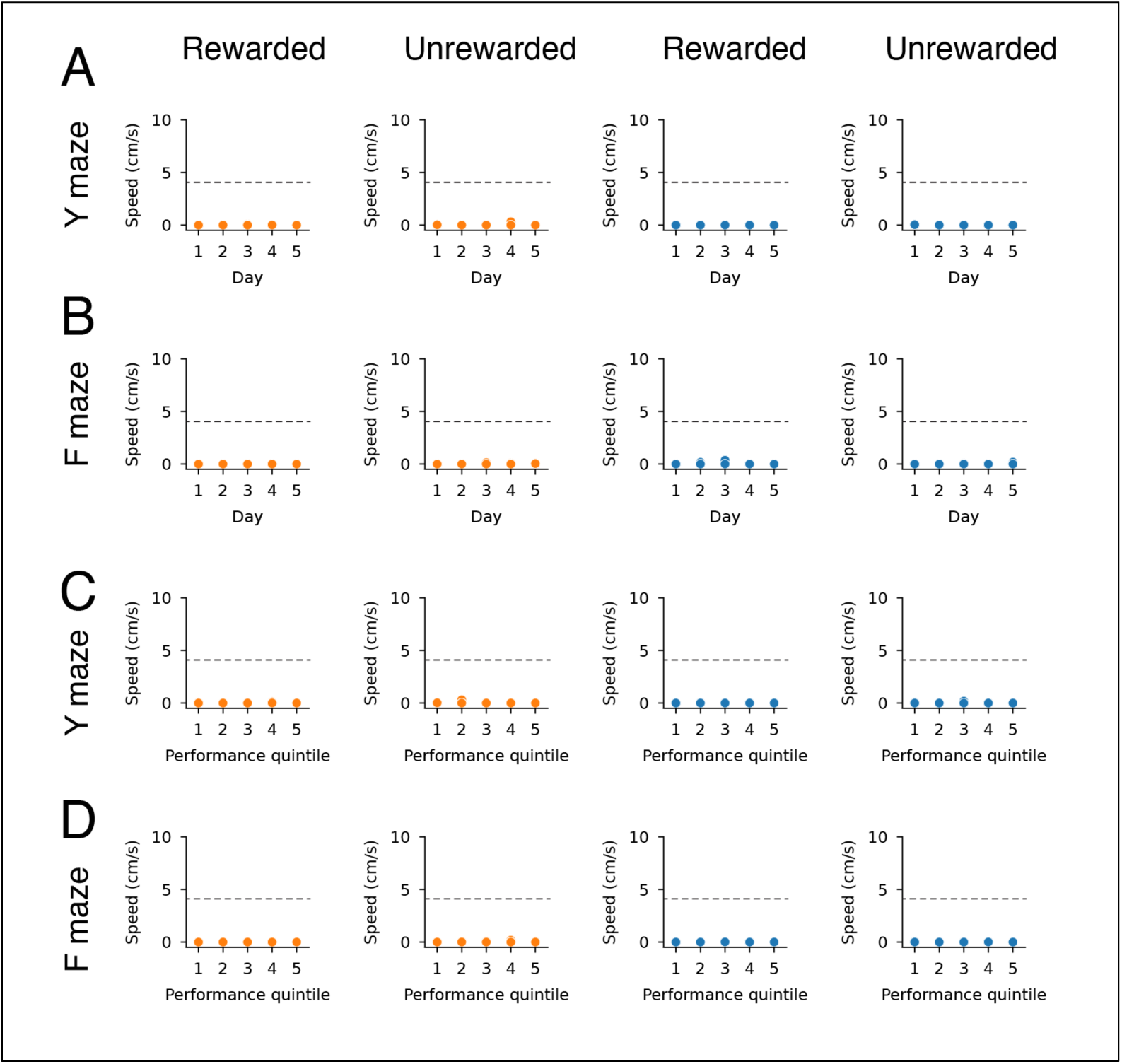
Animals are immobile at the time of beta bursts on rewarded and unrewarded trials. **A.** Animal speed during PFC (orange) and CA1 (blue) bursts in Y maze task versus day for unrewarded or rewarded trials. Dotted line shows the threshold for immobility (4 cm/s). **B.** Animal speed during PFC (orange) and CA1 (blue) bursts in F maze task versus day for unrewarded or rewarded trials. **C.** Animal speed during PFC (orange) and CA1 (blue) bursts in Y maze task versus performance for unrewarded or rewarded trials. **D.** Animal speed during PFC (orange) and CA1 (blue) bursts in F maze task versus performance for unrewarded or rewarded trials.

**Figure 4-2.**
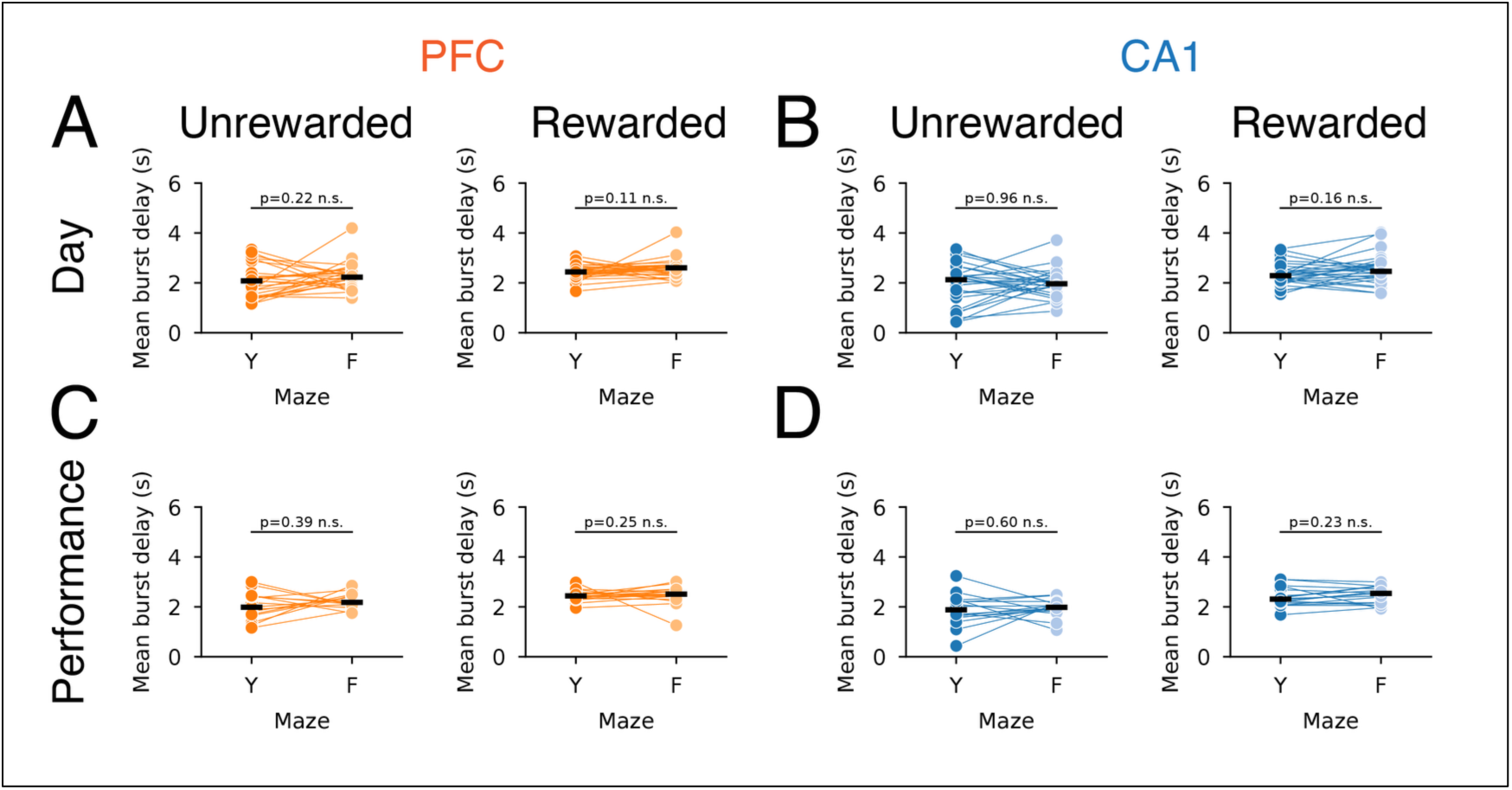
Burst delay by maze for unrewarded and rewarded trials. **A.** PFC burst delay by day (Fig. 4 A). **B.** CA1 burst delay by day (Fig. 4 B). **C.** PFC burst delay by performance (Fig. 4 C). **D.** CA1 burst delay by performance (Fig. 4 D).

**Figure 4-3.**
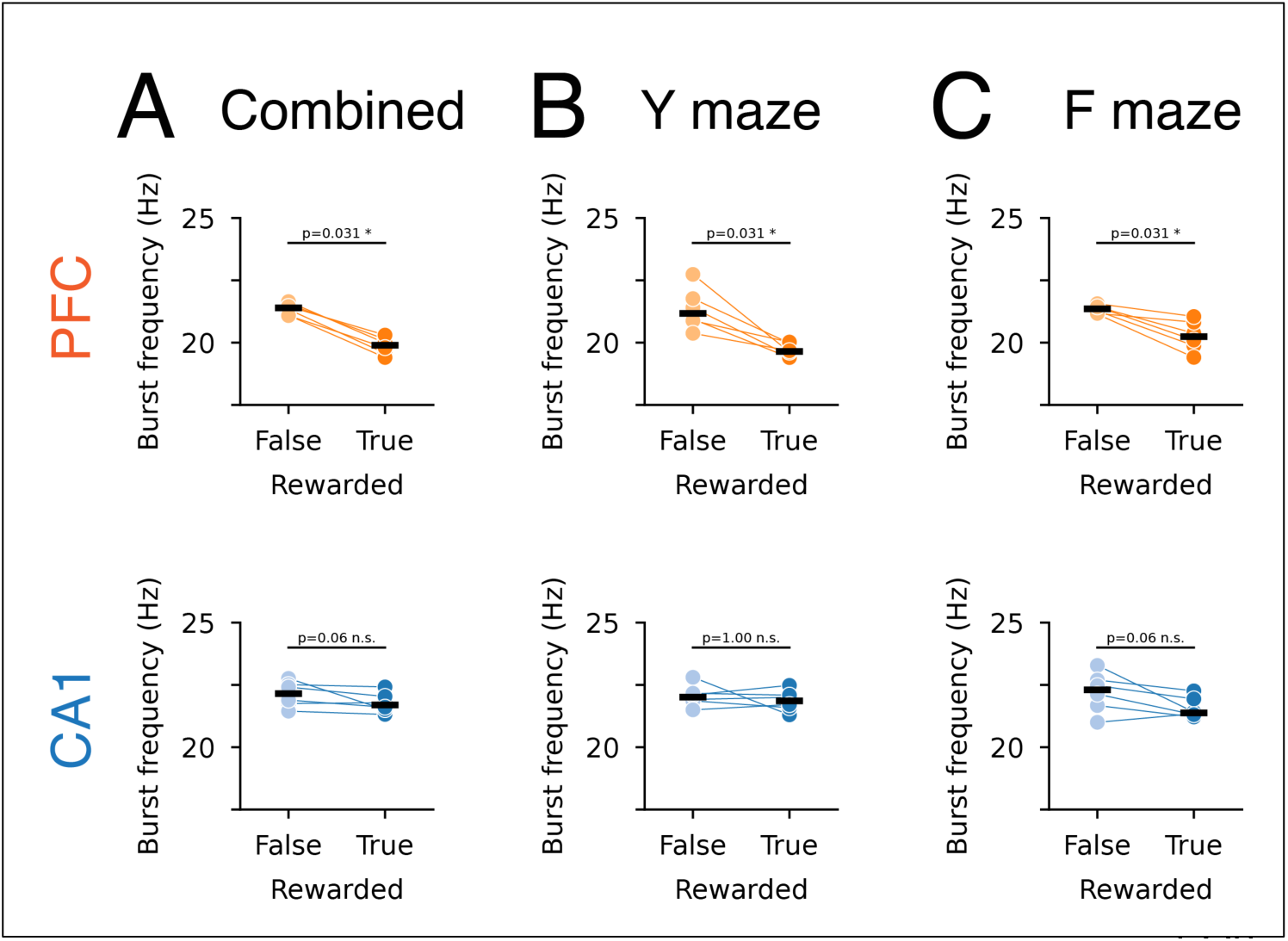
Burst frequency for unrewarded and rewarded trials. **A.** PFC (upper) and CA1 (lower) burst frequency for both maze tasks. The minimum p value for the Wilcoxon signed-rank test (two-sided, exact test without ties) is p=0.031 for a sample size of 6 pairs. **B.** PFC (upper) and CA1 (lower) burst frequency for Y maze task. **C.** PFC (upper) and CA1 (lower) burst frequency for F maze task.

**Figure 4-4.**
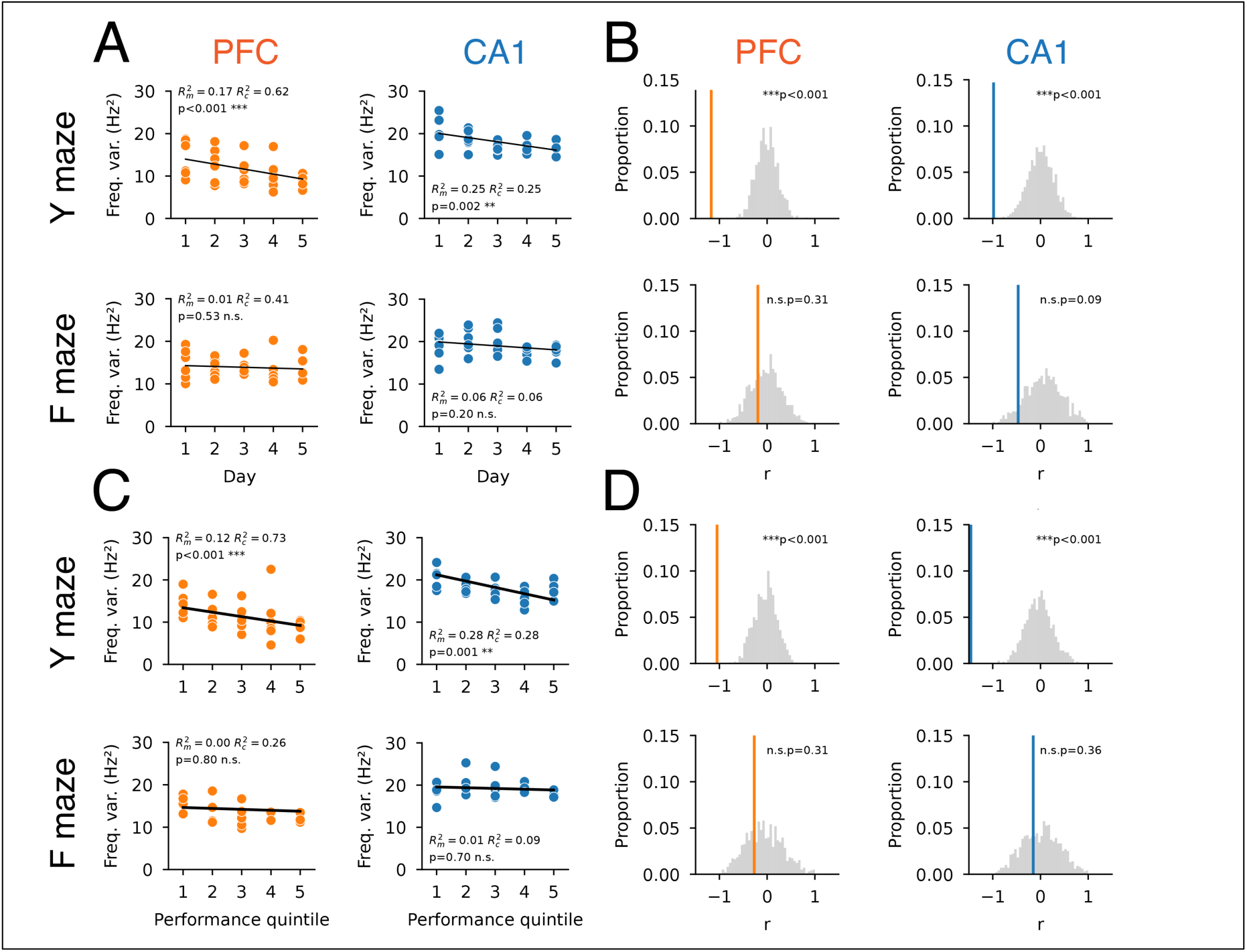
Permutation test for frequency variance across days. **A.** Burst peak frequency variance in each maze task by day. Linear model with fixed (day) and random (animal) effects. **B.** Permutation test for the slope of regression in A. Solid line shows the observed slope. Gray histogram shows the permuted distribution where the experiment day identity was shuffled (n=1000). **C.** Burst peak frequency variance in each maze task by performance quintile. Linear model with fixed (performance quintile) and random (animal) effects. **D.** Permutation test for the slope of regression in C. Solid line shows the observed slope. Gray histogram shows the permuted distribution where the performance quintile was shuffled (n=1000).

**Figure 5-1.**
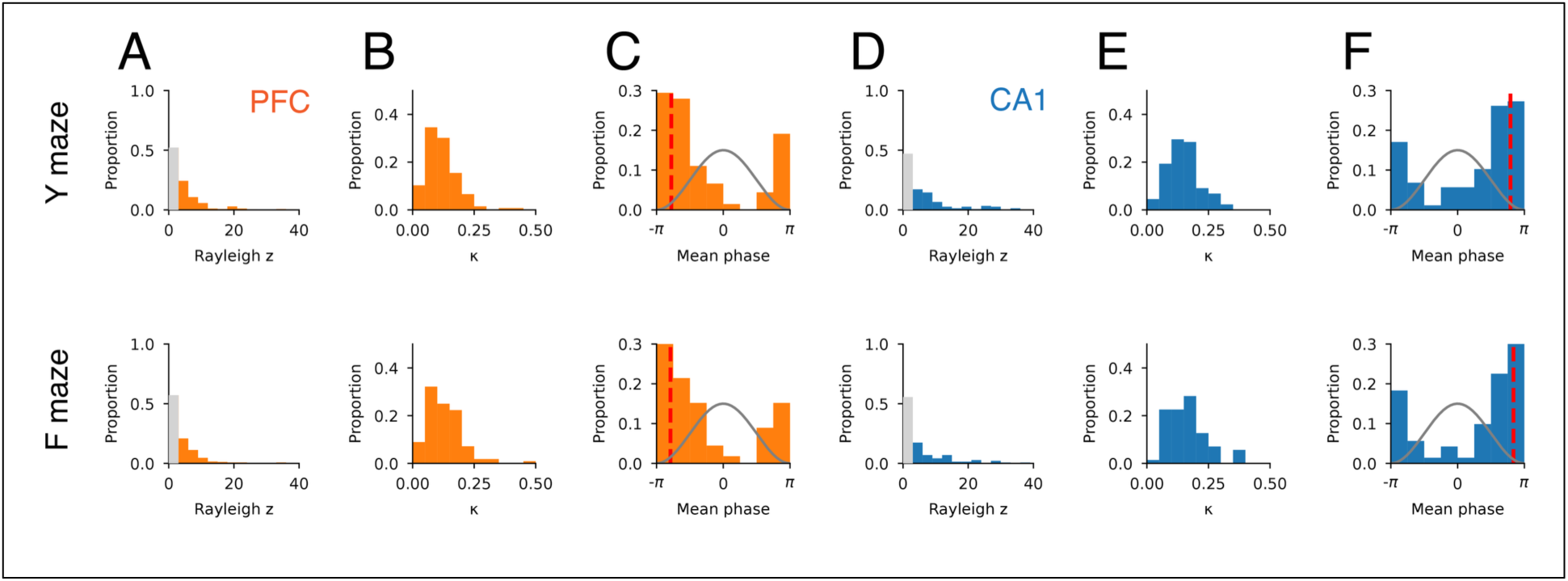
Beta-phase locking by maze task. **A.** Distribution of Rayleigh z for PFC. **B.** Distribution of k from a von Mises distribution fit for significantly beta-modulated cells (Rayleigh p <0.05 in panel A) in PFC. **C.** Distribution of phase preference for significantly beta-modulated cells in PFC. Red dotted line shows the mean population phase preference. The gray line indicates the peak and trough of beta. **D.** Distribution of Rayleigh z for CA1. **E.** Distribution of k from a von Mises distribution fit for significantly beta-modulated cells (Rayleigh p <0.05 in panel D) in CA1. **F.** Distribution of phase preference for significantly beta-modulated cells in CA1. Red dotted line shows the mean population phase preference. The gray line indicates the peak and trough of beta.

**Figure 5-2.**
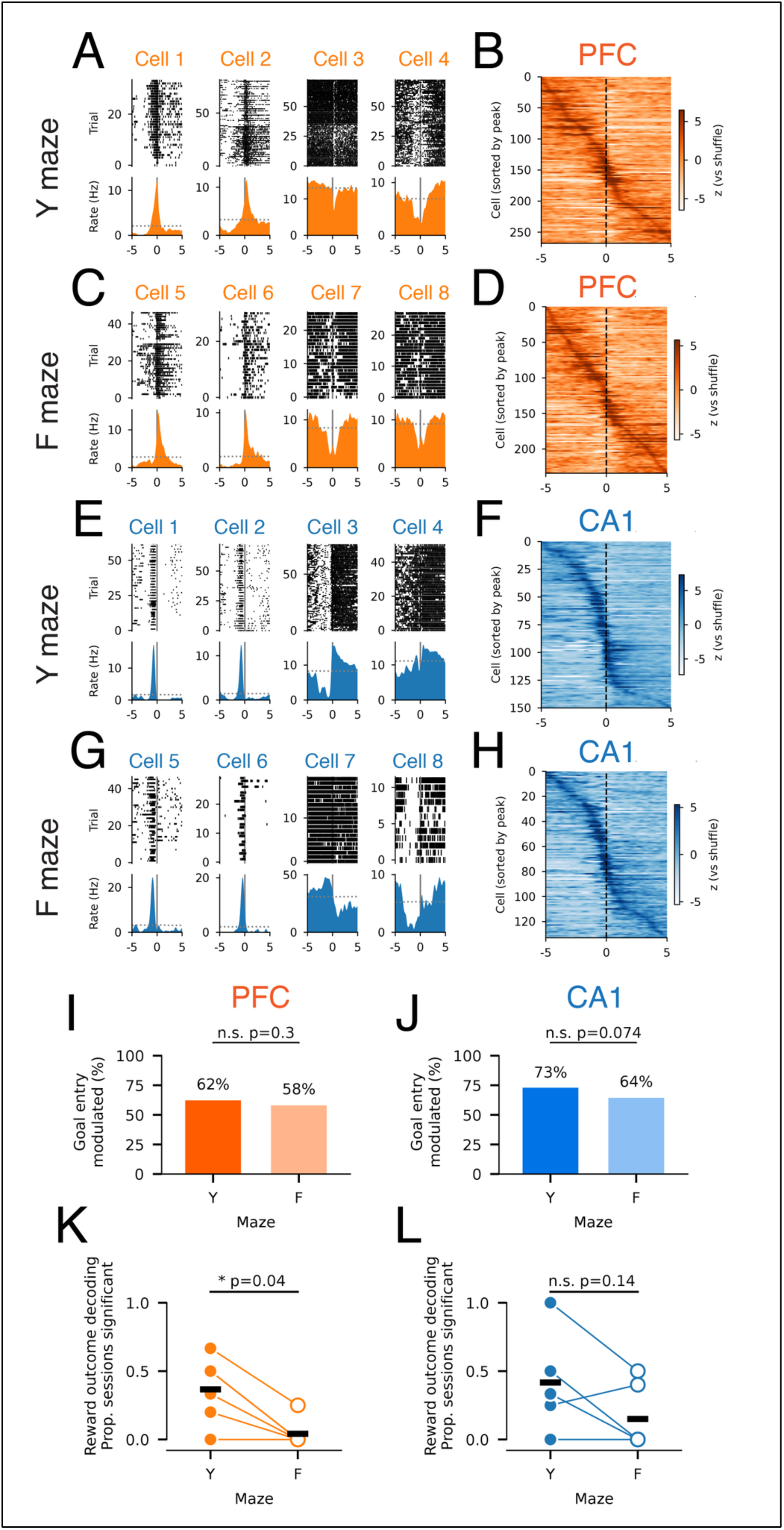
Goal-entry spiking patterns. **A.** Example PFC cells showing trial spike raster and firing rate aligned to goal entry on the Y maze. **B.** Normalized PFC mean firing aligned to goal entry on the Y maze. Cells are sorted by maximum firing rate relative to goal entry on the Y maze. **C.** Example PFC cells showing trial spike raster and firing rate aligned to goal entry on the F maze. **D.** Normalized PFC mean firing aligned to goal entry on the F maze. Same sorting as B. **E.** Example CA1 cells showing trial spike raster and firing rate aligned to goal entry on the Y maze. **F.** Normalized CA1 mean firing aligned to goal entry on the Y maze. Same sorting as B. **G.** Example CA1 cells showing trial spike raster and firing rate aligned to goal entry on the F maze. **H.** Normalized CA1 mean firing aligned to goal entry on the F maze. Same sorting as B. **I.** Proportion of PFC cells with goal-entry spiking modulation, defined as significant spiking deviation from the mean. The proportions for each maze are comparable, Fisher’s exact test: p=0.3. **J.** Same as (I) for CA1. Fisher’s exact test: p=0.074. **K.** Mean proportion of sessions for each animal with significant decoding of goal outcome using PFC population firing in the 1 s after goal entry, for the Y and F mazes. Wilcoxon signed-rank test: p=0.04. Data includes 55 sessions (26 Y-maze, 29 F-maze) from 6 rats; 9.5 ± 3.7 cells per session (mean ± STD); 24.2 ± 10.8 goal visits per session (17.1 rewarded, 7.1 unrewarded). Two animals had identical values for the Y and F mazes. **L.** Same as (K) for CA1. Wilcoxon signed-rank test: p=0.14. Data includes 44 sessions (21 Y-maze, 23 F-maze) from 6 rats; 12.0 ± 7.4 cells per session (mean ± STD); 23.6 ± 10.7 goal visits per session (16.4 rewarded, 7.2 unrewarded).

**Figure 5-3.**
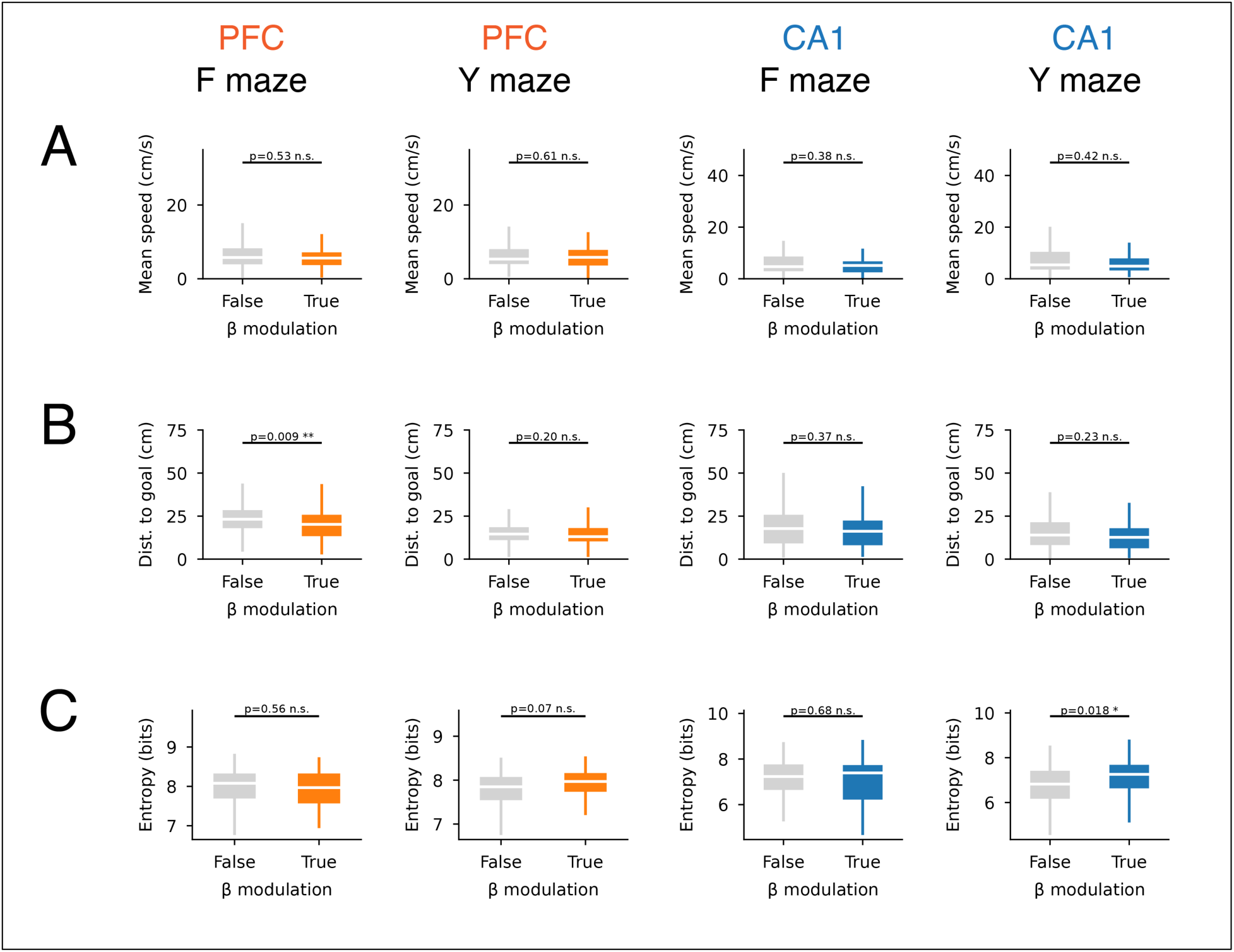
Spiking task correlates by maze task. **A.** Mean spiking speed for PFC and CA1 cells based on beta modulation. **B.** Spiking distance to nearest goal for PFC and CA1 cells based on beta modulation. **C.** Spatial firing entropy for PFC and CA1 cells based on beta modulation.

**Table 5-1.**
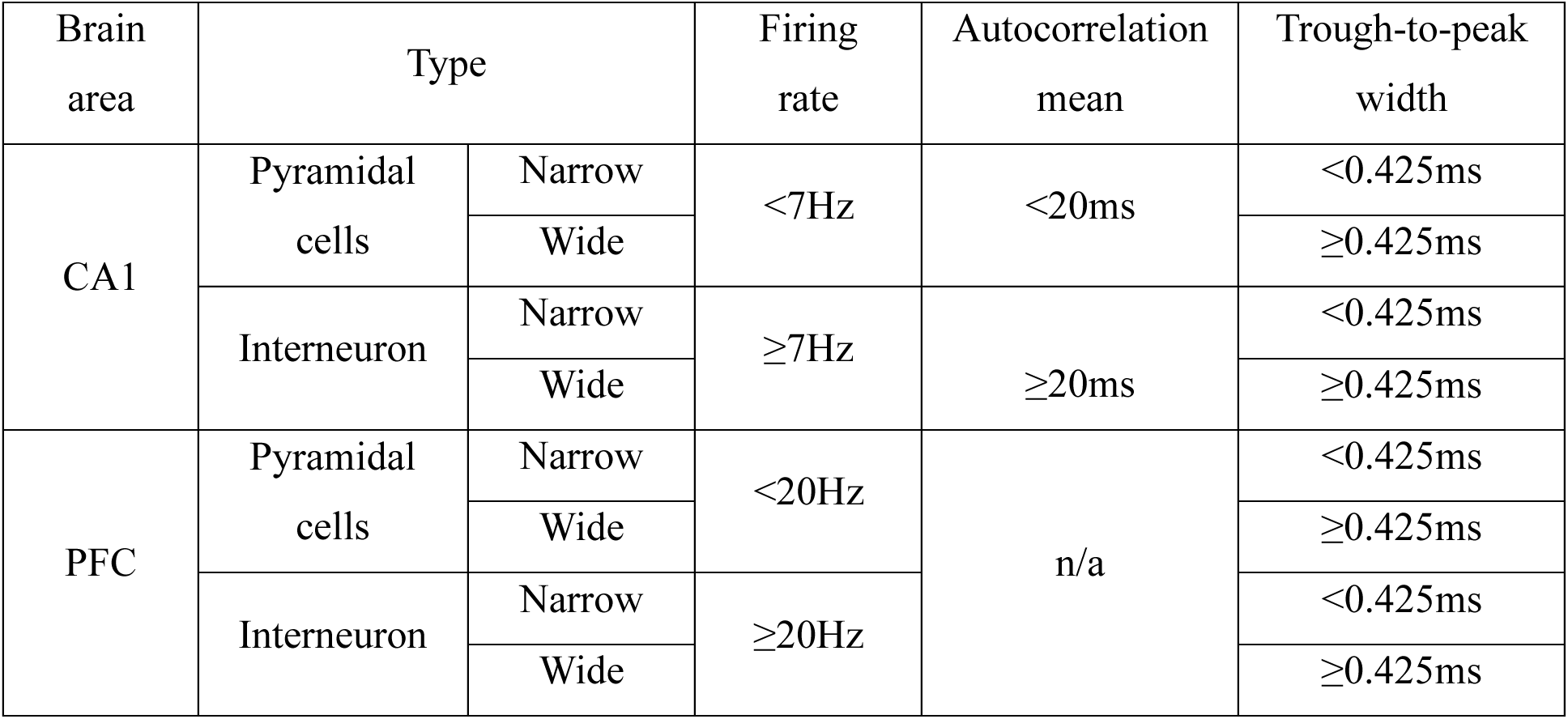
Cell-type classification. Thresholds for cell-type classification.

**Table 5-2.**
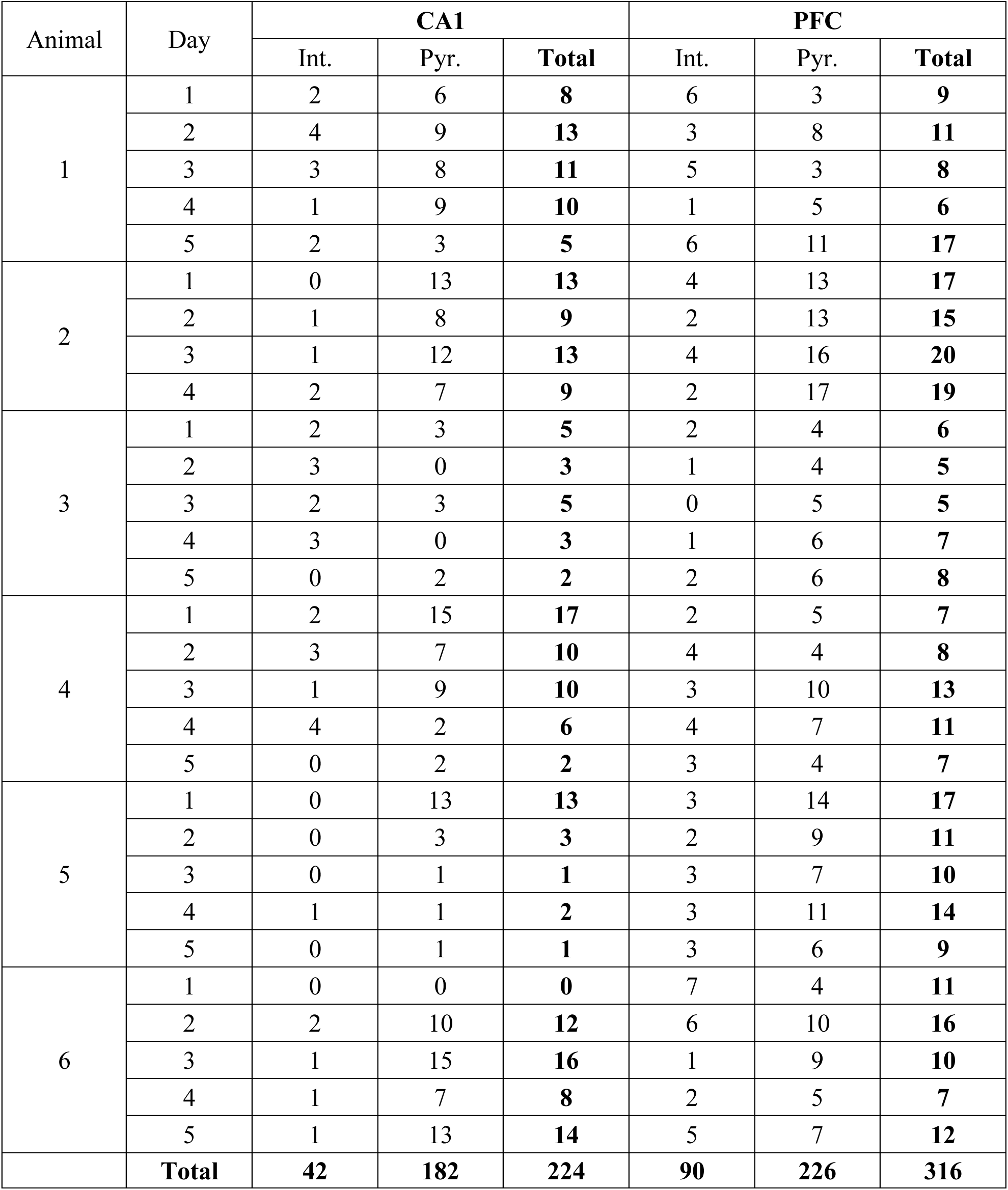
Cell count per day. Number of cells recorded per animal for each day by cell type.

| Animal | Day | CA1 |  |  | PFC |  |  |
| --- | --- | --- | --- | --- | --- | --- | --- |
|  |  | Int. | Pyr. | Total | Int. | Pyr. | Total |
| 1 | 1 | 2 | 6 | 8 | 6 | 3 | 9 |
|  | 2 | 4 | 9 | 13 | 3 | 8 | 11 |
|  | 3 | 3 | 8 | 11 | 5 | 3 | 8 |
|  | 4 | 1 | 9 | 10 | 1 | 5 | 6 |
|  | 5 | 2 | 3 | 5 | 6 | 11 | 17 |
| 2 | 1 | 0 | 13 | 13 | 4 | 13 | 17 |
|  | 2 | 1 | 8 | 9 | 2 | 13 | 15 |
|  | 3 | 1 | 12 | 13 | 4 | 16 | 20 |
|  | 4 | 2 | 7 | 9 | 2 | 17 | 19 |
| 3 | 1 | 2 | 3 | 5 | 2 | 4 | 6 |
|  | 2 | 3 | 0 | 3 | 1 | 4 | 5 |
|  | 3 | 2 | 3 | 5 | 0 | 5 | 5 |
|  | 4 | 3 | 0 | 3 | 1 | 6 | 7 |
|  | 5 | 0 | 2 | 2 | 2 | 6 | 8 |
| 4 | 1 | 2 | 15 | 17 | 2 | 5 | 7 |
|  | 2 | 3 | 7 | 10 | 4 | 4 | 8 |
|  | 3 | 1 | 9 | 10 | 3 | 10 | 13 |
|  | 4 | 4 | 2 | 6 | 4 | 7 | 11 |
|  | 5 | 0 | 2 | 2 | 3 | 4 | 7 |
| 5 | 1 | 0 | 13 | 13 | 3 | 14 | 17 |
|  | 2 | 0 | 3 | 3 | 2 | 9 | 11 |
|  | 3 | 0 | 1 | 1 | 3 | 7 | 10 |
|  | 4 | 1 | 1 | 2 | 3 | 11 | 14 |
|  | 5 | 0 | 1 | 1 | 3 | 6 | 9 |
| 6 | 1 | 0 | 0 | 0 | 7 | 4 | 11 |
|  | 2 | 2 | 10 | 12 | 6 | 10 | 16 |
|  | 3 | 1 | 15 | 16 | 1 | 9 | 10 |
|  | 4 | 1 | 7 | 8 | 2 | 5 | 7 |
|  | 5 | 1 | 13 | 14 | 5 | 7 | 12 |
|  | Total | 42 | 182 | 224 | 90 | 226 | 316 |

**Table 5-3.** Cell count by maze. Number of cells recorded per animal across all days in each maze task that met the inclusion conditions (see Methods: Cell-type classification). The “Unique” column counts each cell once per day, regardless of its activity in one or both maze tasks.

| Animal | CA1 |  |  | PFC |  |  |
| --- | --- | --- | --- | --- | --- | --- |
|  | Y | F | Unique | Y | F | Unique |
| 1 | 35 | 32 | **47** | 48 | 31 | **51** |
| 2 | 30 | 21 | **44** | 61 | 57 | **71** |
| 3 | 14 | 13 | **18** | 26 | 28 | **31** |
| 4 | 32 | 29 | **45** | 39 | 33 | **46** |
| 5 | 13 | 13 | **20** | 52 | 48 | **61** |
| 6 | 26 | 33 | **50** | 42 | 48 | **56** |
| **Total** | **150** | **141** | **224** | **268** | **245** | **316** |

**Table 5-4.** Cell count by beta-phase locking. Number of cells recorded per animal classified based on beta-phase-locking status.

| Animal | CA1 |  |  | PFC |  |  |
| --- | --- | --- | --- | --- | --- | --- |
|  | Not beta phase-locked | Beta phase-locked | Total | Not beta phase-locked | Beta phase-locked | Total |
| 1 | 31 | 16 | 47 | 41 | 10 | 51 |
| 2 | 32 | 12 | 44 | 48 | 23 | 71 |
| 3 | 11 | 7 | 18 | 26 | 5 | 31 |
| 4 | 34 | 11 | 45 | 38 | 8 | 46 |
| 5 | 14 | 6 | 20 | 46 | 15 | 61 |
| 6 | 30 | 20 | 50 | 43 | 13 | 56 |
| **Total** | **152** | **72** | **224** | **242** | **74** | **316** |

**Figure 6-1.**
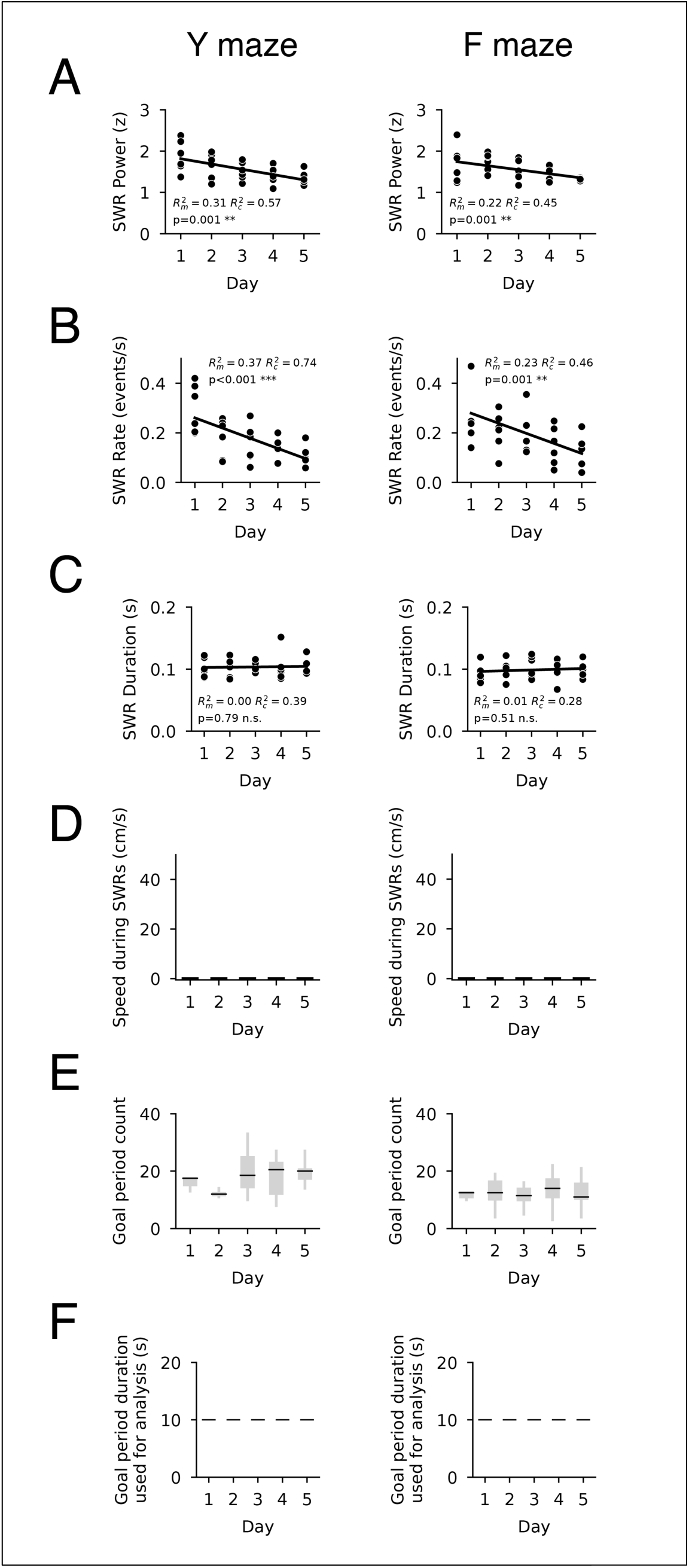
CA1 SWR properties versus day by maze task. **A.** SWR power. **B.** SWR rate. **C.** SWR duration. **D.** Animal speed during SWR. **E.** Goal period count per animal. **F.** Goal period duration.

**Figure 6-2.**
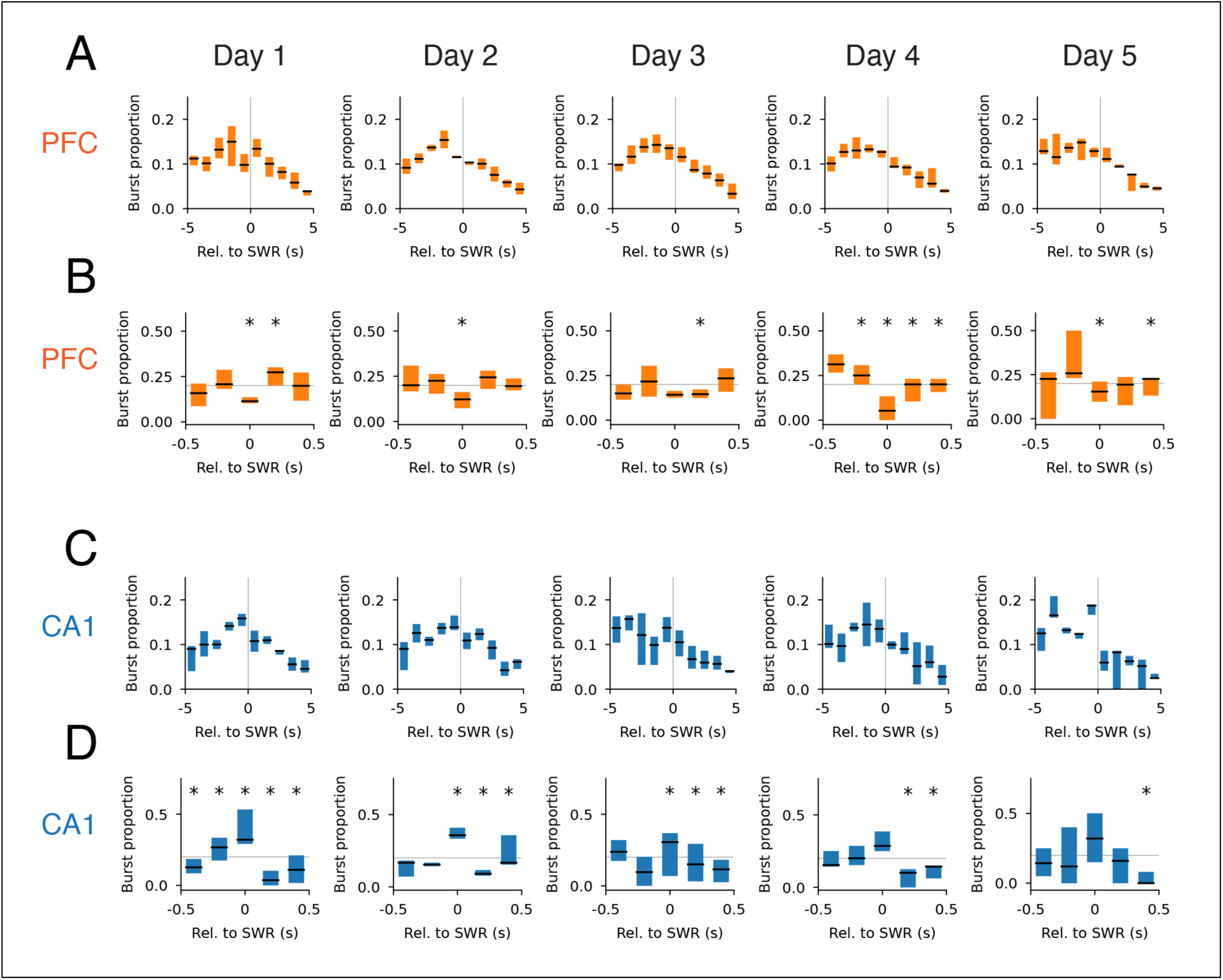
Beta burst timing relative to SWRs across days of learning. **A.** Cross-correlation between PFC beta bursts and CA1 SWRs (±5 s, 1 s bins) for days 1-5. The beta bursts occur before SWRs, Wilcoxon signed-rank test for burst timing relative to SWRs, p<0.01 for all days. **B.** Cross-correlation between PFC beta bursts and CA1 SWRs (±500 ms, 200 ms bins). Bins with significant deviations from the mean (gray line) are indicated with * (with Benjamini-Hochberg false discovery rate correction). **C.** Same as (A) for CA1. **D.** Same as (B) for CA1.

**Figure 6-3.**
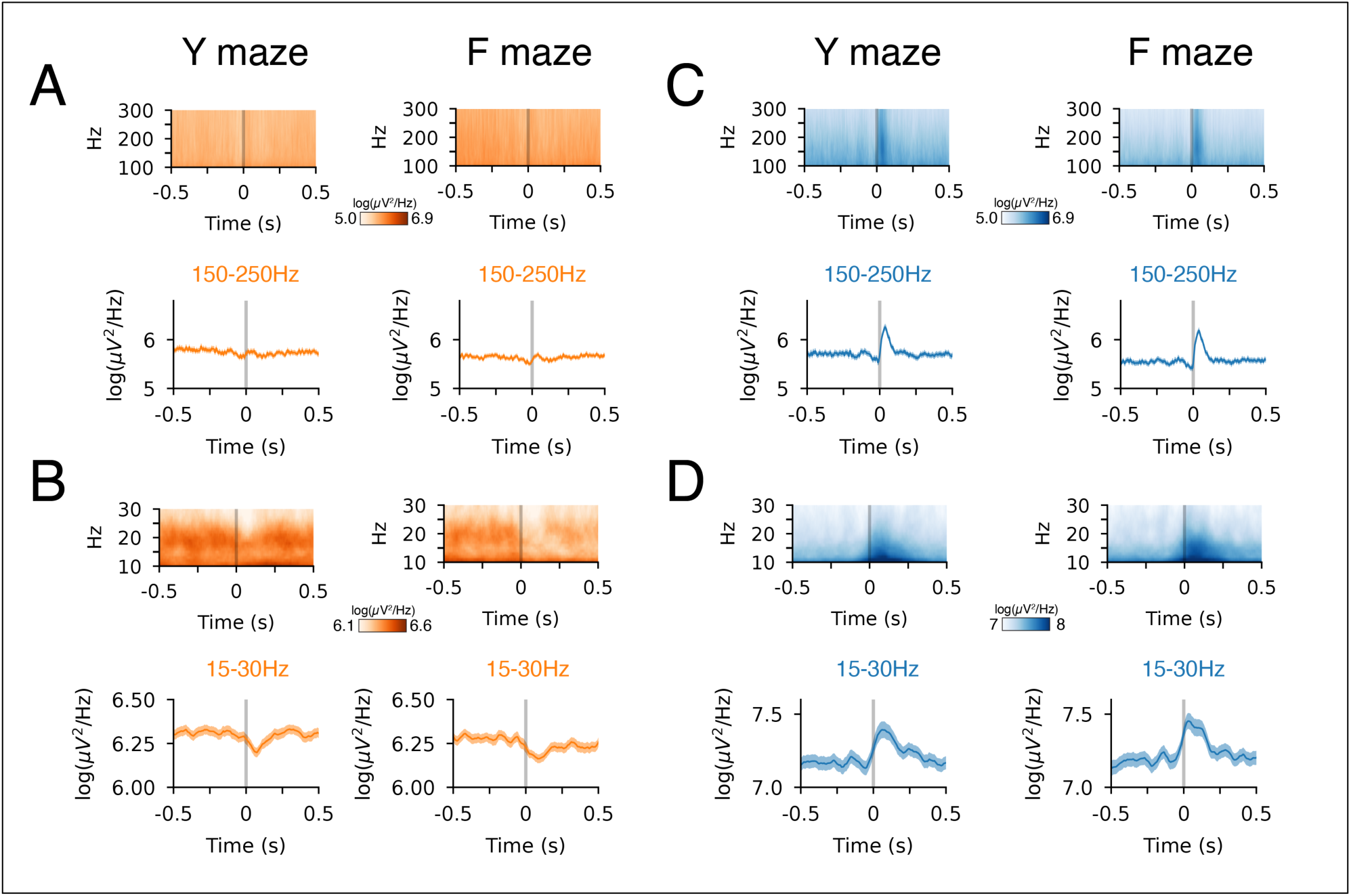
SWR-aligned spectrogram by maze task. **A.** SWR-aligned PFC spectrogram (100-300 Hz) and corresponding ripple band power (150-250 Hz). **B.** SWR-aligned PFC spectrogram (10-30 Hz) and corresponding beta-band power (15-30 Hz). **C.** SWR-aligned CA1 spectrogram (100-300 Hz) and corresponding ripple band power (150-250 Hz). **D.** SWR-aligned CA1 spectrogram (10-30 Hz) and corresponding beta-band power (15-30 Hz).

**Figure 6-4.**
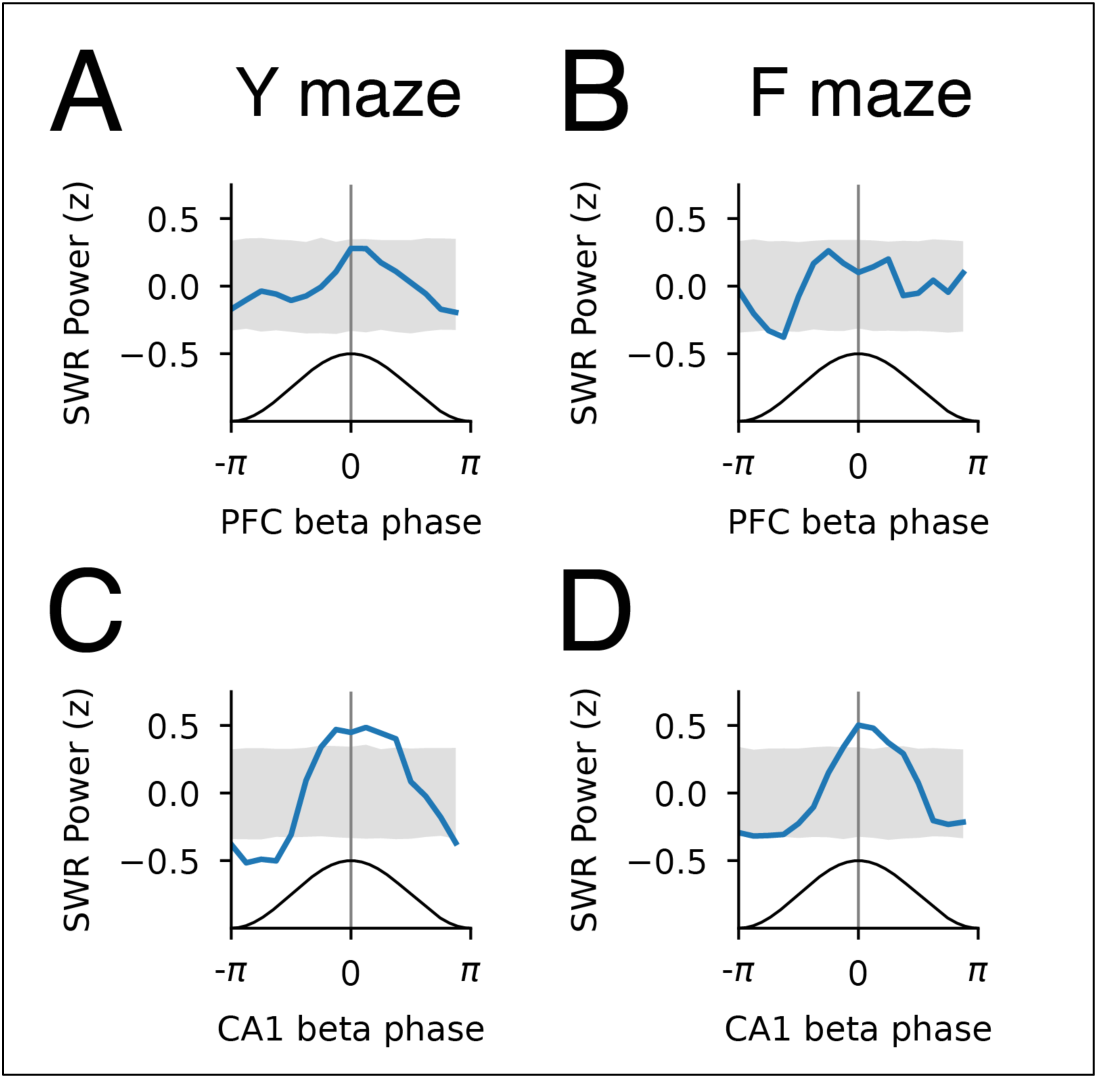
Phase amplitude coupling between SWR band power and beta phase during SWRs. **A.** Y maze SWR band power relative to PFC beta phase. The normalized mean SWR band power (z) across all sessions is shown in blue. The 99% confidence interval from a circular permutation test (5000 shuffles) is shown in gray. The permutation test circularly permutes the phase value for each session. **B.** F maze SWR band power relative to PFC beta phase. **C.** Y maze SWR band power relative to CA1 beta phase. SWR band power showed significant deviation from the shuffle bounds around the peak of beta. **D.** F maze SWR band power relative to CA1 beta phase. SWR band power showed significant deviation from the shuffle bounds around the peak of beta.

**Figure 7-1.**
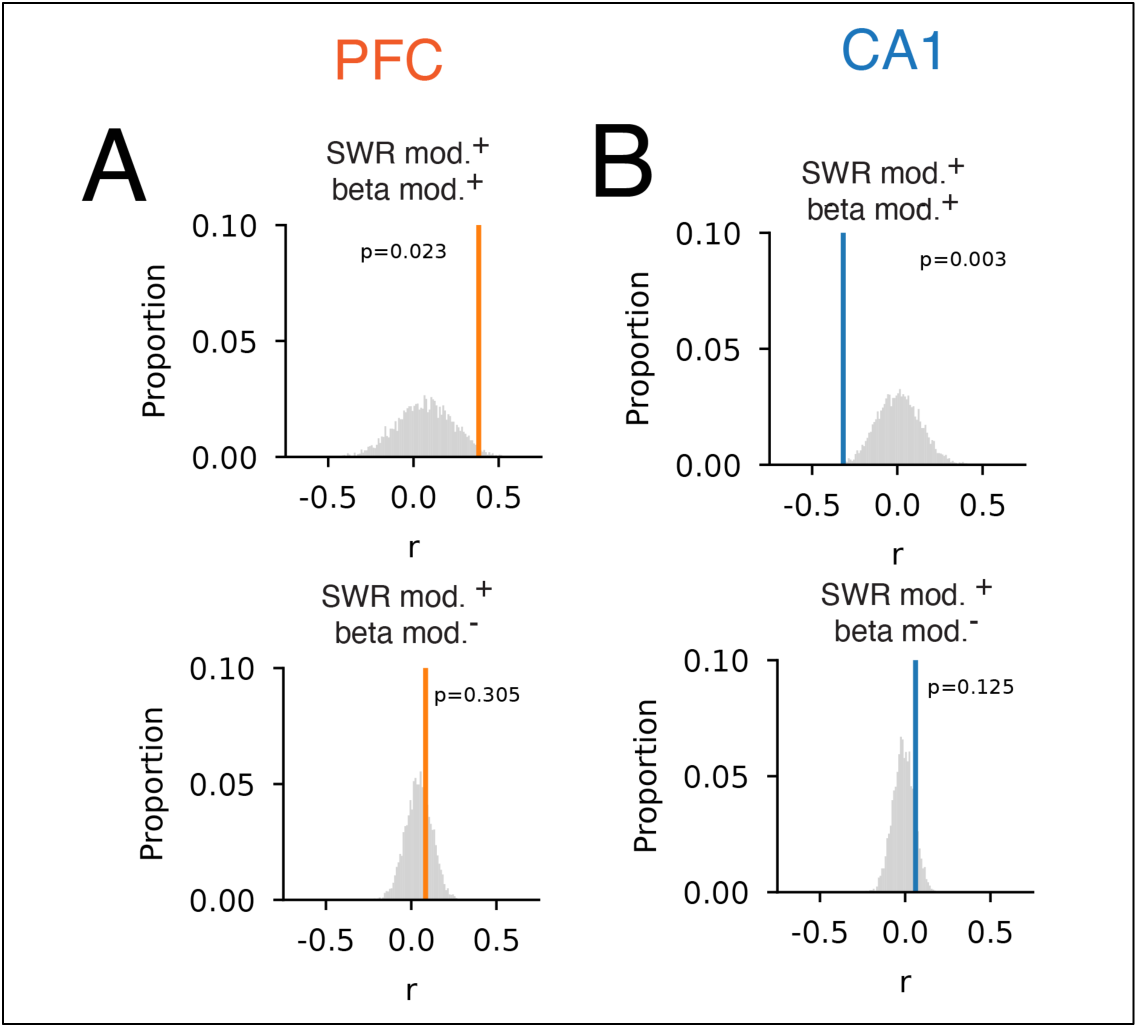
Permutation test for the significance of the correlation coefficient. **A.** Distribution of the permuted (gray) and observed (vertical line) correlation coefficients for PFC SWR-modulation index versus beta-phase-locking preference in Fig. 7 C. We performed this additional permutation test to verify that the observed value of the correlation coefficient is different from the chance distribution. **B.** Distribution of the permuted (gray) and observed (vertical line) correlation coefficients for CA1 SWR-modulation index versus beta-phase-locking preference in Fig. 7 I.

**Figure 7-2.**
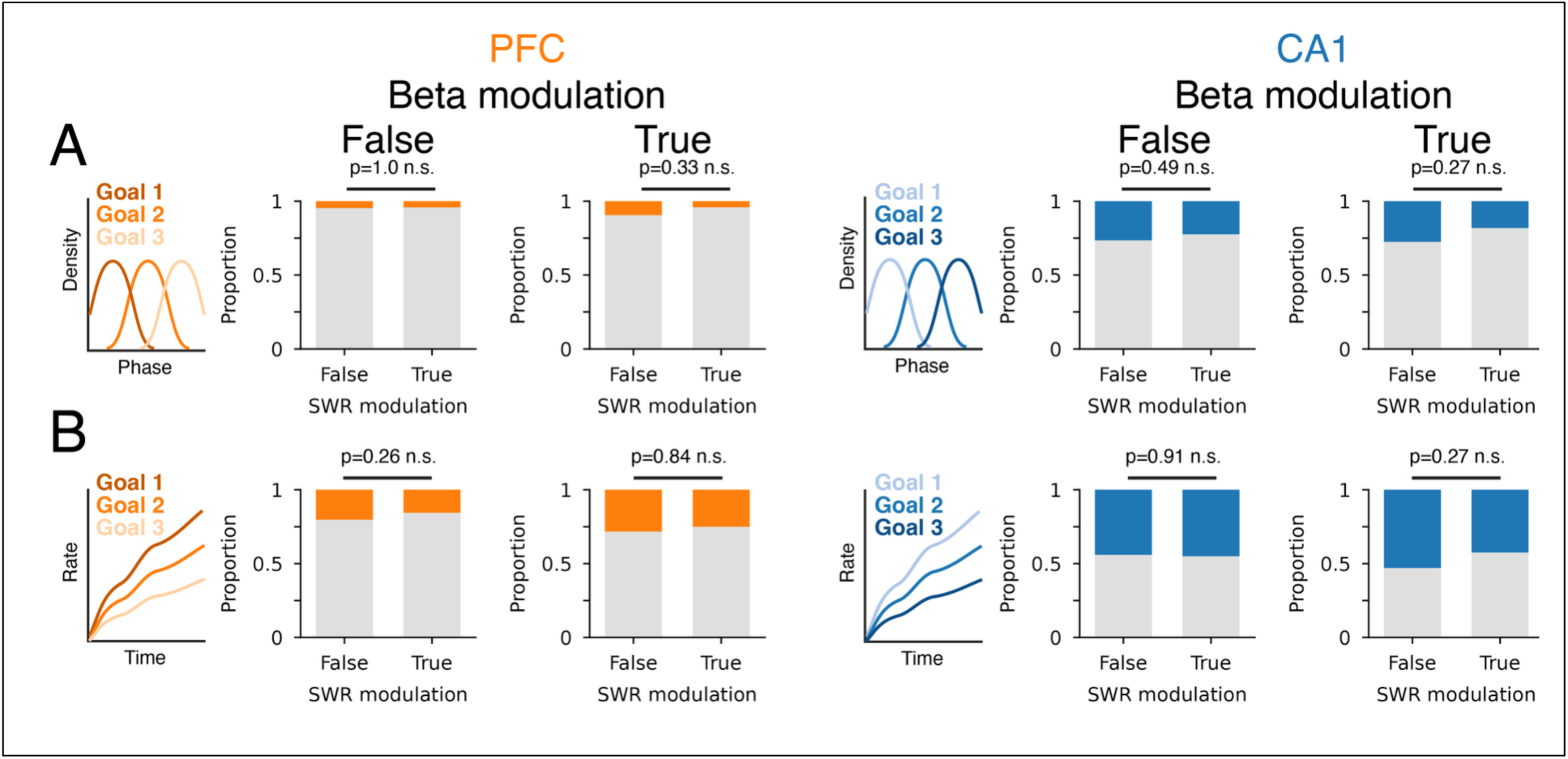
Task correlates by beta and SWR modulation. **A.** Proportion of each PFC (orange, left two columns) and CA1 (blue, right two columns) population that distinguished goal identity or did not (gray) based on phase preference at each goal. **B.** Proportion of each PFC (orange) and CA1 (blue) population that distinguished goal identity or did not (gray) based on mean firing rate at each goal.

## Notes

### Competing Interest Statement

The authors have declared no competing interest.

### Summary of Updates

Revised the text in response to reviewers' comments and to improved readability. Captions added for all tables. The scaling concern and corresponding edits are explained in the response. Table 2-1 has been corrected. The previous version did not show the correct trial count for all sessions for each day. Added two new figures (Figs. 5-2 and 6-2); former Figs. 6-2 and 6-3 are now Figs. 6-3 and 6-4.

